# Introgression dynamics of sex-linked chromosomal inversions shape the Malawi cichlid adaptive radiation

**DOI:** 10.1101/2024.07.28.605452

**Authors:** L. M. Blumer, V. Burskaia, I. Artiushin, J. Saha, J. Camacho Garcia, F. Campuzano Jiménez, A. Hooft van der Huysdynen, J. Elkin, B. Fischer, N. Van Houtte, C. Zhou, S. Gresham, M. Malinsky, T. Linderoth, W. Sawasawa, G. Vernaz, I. Bista, A. Hickey, M. Kucka, S. Louzada, R. Zatha, F. Yang, B. Rusuwa, M. E. Santos, Y. F. Chan, D. A. Joyce, A. Böhne, E. A. Miska, M. Ngochera, G. F. Turner, R. Durbin, H. Svardal

**Affiliations:** Department of Genetics, University of Cambridge, Cambridge, England, CB2 3EH, UK; Evolutionary Ecology Group, Department of Biology, University of Antwerp, 2020 Antwerp, Belgium; Department of Zoology, University of Cambridge, Cambridge, CB2 3EJ, UK; Wellcome Sanger Institute, Tree of Life, Wellcome Genome Campus, Hinxton, CB10 1SA, UK; Institute of Ecology and Evolution, Department of Biology, University of Bern, Baltzerstrasse 6, 3012 Bern, Switzerland; W.K. Kellogg Biological Station, Michigan State University, Hickory Corners, MI 49060, USA; Senckenberg Research Institute and Natural History Museum, 60325 Frankfurt am Main, Germany; LOEWE Centre for Translational Biodiversity Genomics, 60325 Frankfurt am Main, Germany; Friedrich Miescher Laboratory of the Max Planck Society, Max-Planck-Ring 9, 72076 Tübingen, Germany; Wellcome Sanger Institute, Wellcome Genome Campus, Hinxton, Cambridge CB10 1SA, UK; School of Natural and Applied Sciences, University of Malawi, Zomba, Malawi; School of Life Sciences and Medicine, Shandong University of Technology, Zibo, China; Groningen Institute for Evolutionary Life Sciences (GELIFES), University of Groningen, Nijenborgh 4, 9747AG Groningen, The Netherlands; Evolutionary and Ecological Genomics Group, School of Natural Sciences, University of Hull, Hull, UK; Leibniz Institute for the Analysis of Biodiversity Change, Museum Koenig Bonn, Adenauerallee 127, 53115 Bonn, Germany; Department of Biochemistry, University of Cambridge, Cambridge, England, CB2 3EH, UK; Department of Fisheries Headquarters, P.O. Box 593, Lilongwe, Malawi; School of Environmental and Natural Sciences, Bangor University, Bangor, Gwynedd, LL57 2TH, Wales, UK; Naturalis Biodiversity Center, 2333 Leiden, The Netherlands

**Author notes:** These authors contributed equally to this work (L. M. Blumer, V. Burskaia, I. Artiushin, J. Saha). Zoological Institute, University of Basel, 4051 Basel, CH. Department of Translational Genomics, University of Cologne, 50931 Cologne, Germany (M. Kucka). Laboratory of Cytogenomics and Animal Genomics, Department of Genetics and Biotechnology, University of Trás-os-Montes and Alto Douro, Vila Real, Portugal; BioSystems and Integrative Sciences Institute, Faculty of Sciences, University of Lisbon, Lisbon, Portugal (S. Louzada).

## Abstract

Chromosomal inversions contribute to adaptive speciation by linking co-adapted alleles. Querying 1,375 genomes of the species-rich Malawi cichlid fish radiation, we discovered five large inversions segregating in the benthic subradiation that each suppress recombination over more than half a chromosome. Two inversions were transferred from deepwater pelagic *Diplotaxodon* via admixture, while the others established early in the *deep benthic* clade. Introgression of haplotypes from lineages inside and outside the Malawi radiation coincided with bursts of species diversification. Inversions show evidence for transient sex linkage and a striking excess of protein changing substitutions points towards selection on neuro-sensory, physiological and reproductive genes. We conclude that repeated interplay between depth adaptation and sex-specific selection on large inversions has been central to the evolution of this iconic system.

## Main

Understanding how biodiversity evolves is a fundamental question in biology. While some evolutionary lineages remain virtually unchanged over hundreds of millions of years (*1*), others give rise to a great diversity of species over short evolutionary timescales (*2*). Adaptive radiations are particularly remarkable examples of explosive diversification, with many ecologically, morphologically, and behaviourally differentiated species emerging rapidly from a common ancestor. It is still not well understood how evolutionary lineages can produce such bursts of organismal diversity, but recent insights from genome sequencing point to a widespread contribution of “old” genetic variants (*3*), often introduced into populations by hybridisation (*4*), and reused in new combinations that provide adaptation to novel ecological niches (*5*). A conundrum, however, is the role of meiotic recombination in this process. On the one hand, recombination can create beneficial combinations of adaptive alleles (*6*). On the other hand, recombination can break adaptive combinations apart (*7*), especially in the face of gene flow, producing unfit intermediates (*8*), and impeding speciation (*9*).

Chromosomal inversions – stretches of DNA that are flipped in their orientation – provide a mechanism to break the apparent deadlock between the beneficial and detrimental effects of recombination on species diversification, by strongly suppressing recombination between the inverted haplotype and its ancestral configuration (*7, 10, 11*). Inverted haplotypes acting as “supergenes” can link together adaptive alleles that confer a fitness advantage in a specific environmental context or species background (*12*). In recent years, inversions have increasingly been found to contribute to adaptation (*13, 14*), genetic incompatibilities (*15*), assortative mating (*16*), sexual dimorphism (*17, 18*), mating systems (*19*), social organisation (*20*), life-history strategies (*21*) and other complex phenotypes (*11*). Inversions are more common between sympatric than allopatric sister species in fruit flies (*22*), rodents (*23*), and passerine birds (*24*), pointing to their involvement in speciation with gene flow. However, despite their evolutionary relevance in other systems, there is relatively little information on their role in shaping large vertebrate adaptive radiations (*25*–*28*).

With over 800 known extant species, Lake Malawi cichlids constitute the most species-rich recent vertebrate adaptive radiation (*29, 30*). The radiation was able to unfold and generate extraordinary morphological and ecological diversity, despite repeated hybridisation (*31, 32*) and conserved fertility across species (*33*). Intriguingly, previous studies found broad genetic association peaks for a behavioural phenotype important in assortative mating (*34*) in genomic regions that showed suppressed recombination in crosses of Malawi cichlid species (*35*). This raises the question of whether recombination-suppressing mechanisms such as inversions contributed to the adaptive diversification of Malawi cichlids.

Here we show that five large inversions segregate across and within many species and groups in the Lake Malawi radiation, and systematically investigate their evolutionary histories and functions. By suppressing recombination, large chromosomal inversions can cause affected genomic regions to show evolutionary histories consistently distinct from the rest of the genome (*36, 37*). To detect regional deviations from the genome wide evolutionary history we obtained whole genome sequencing (WGS) data from 1,375 individuals of 240 Malawi cichlid species (table S1), detected 84 million single nucleotide polymorphisms (SNPs), and first inferred genome-wide relationship patterns as a backbone (Fig. 1, fig. S1) (see materials and methods). Previous work suggested that the Malawi radiation evolved through serial diversification of three subradiations from a riverine-like ancestor (*31*): 1. A pelagic grouping of the mostly mid-water *Rhamphochromis* and mostly deep-living *Diplotaxodon*; 2. an ecologically and morphologically highly diverse *benthic* subradiation consisting of three subgroups – *deep benthics, shallow benthics*, and semi open-water *utaka*; and 3. the predominantly rock-dwelling *mbuna*. The generalist-like stem lineage is represented today by the Malawi cichlid species *Astatotilapia calliptera* (which provided the reference genome for the present study) (*31, 38*). Phylogenetic inference on our larger dataset confirms these major groupings and supports the branching order (Fig. 1).

**Fig. 1:**
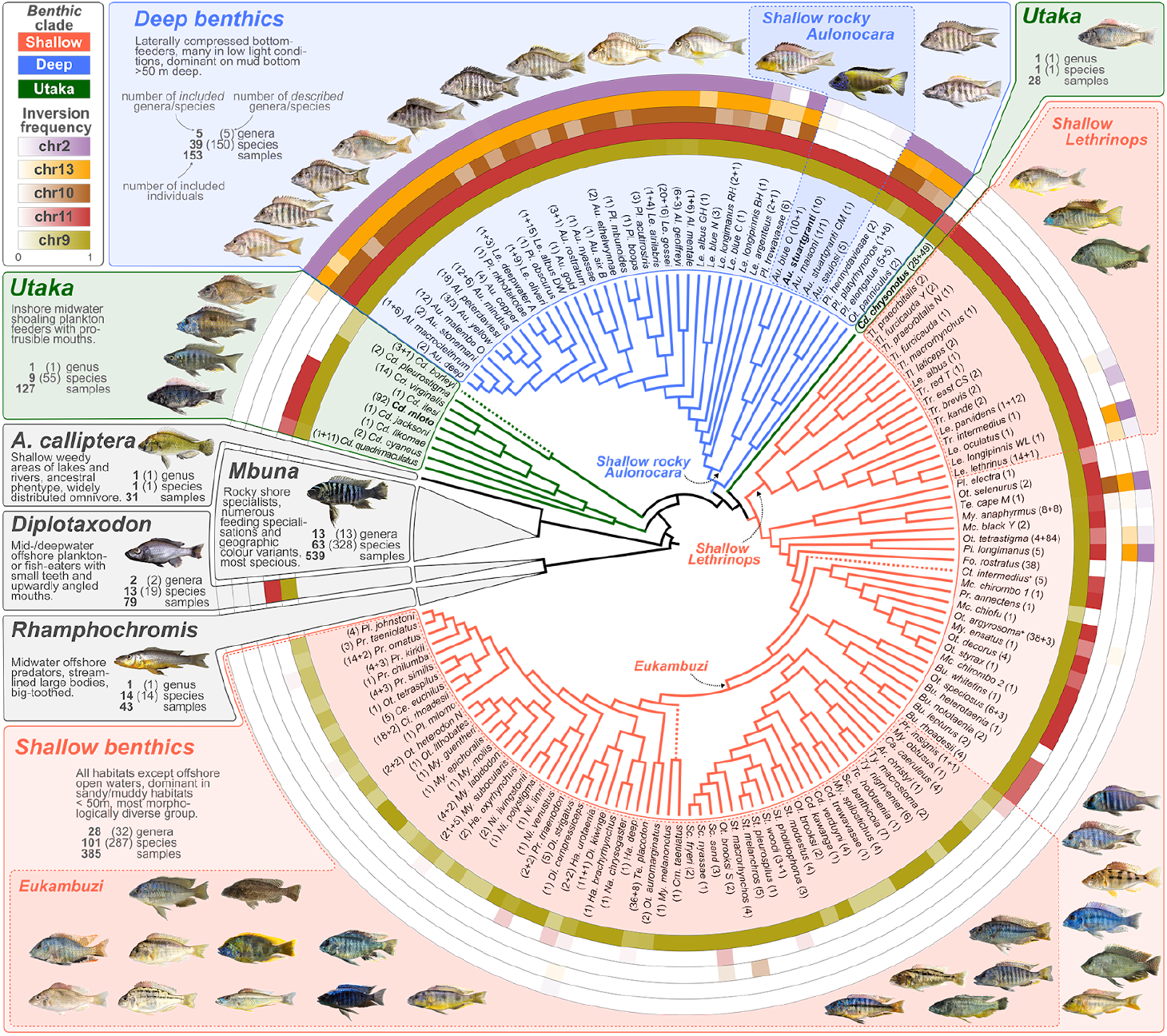
Study system and prevalence of five large inversions. Consensus phylogeny of Malawi cichlid species used in this study (data S1) with inversion frequency based on WGS and PCR-typing (see materials and methods) shown in rings around the phylogeny (the same colours are used throughout the article). The *benthic* subradiation is expanded to show the phylogenetic position of each species and to highlight subclades that we refer to in the main text (*Shallow rocky Aulonocara, Shallow Lethrinops, Eukambuzi*). Note that *utaka* are not monophyletic in this phylogeny. Non-*benthic* groups of Malawi cichlids (i.e., the *pelagic* subradiations of *Rhamphochromis* and *Diplotaxodon*, the subradiation of predominantly rock-dwelling *mbuna*, and *Astatotilapia calliptera* – a species distributed in rivers and margins around the lake that shares its putatively ancestral characteristics and genus assignment with riverine haplochromines outside of the radiation) are each represented by a single grey triangle approximately reflecting species richness relative to each other. Dashed lines indicate branches with unstable placement. See data S1 and S2 for full phylogenies with branch lengths and support values. Annotations next to species/clade names provide the numbers of sequenced/inversion-genotyped samples (additional samples which were inversion-genotyped with PCR are indicated as *+ n* in the annotation). Two taxa are annotated with ‘^+^’ to denote polyphyletic groups: *Otopharynx argyrosoma* contains a single *Cyrtocara moorii* individual and *Ctenopharynx intermedius* contains two *Ctenopharynx pictus* individuals. Full species names are given in table S3 and inversion frequencies by species in table S2. Species names for representative photographs are given in fig. S1. Tree files are given in data S1 and S2. Species subject to further experimental investigation (see text) are highlighted in bold.

### Large inversions suppress recombination

We identified extensive genomic outlier regions consistent with polymorphic inversions on five chromosomes (2, 9, 10, 11 and 13), each spanning more than half a chromosome (between 17 and 23 Mbp), using a clustering approach on our SNP data set (Fig. 2A, top panel, figs. S2, S3) (see materials and methods). A windowed principal component (PC) analysis (*37, 39*) of genetic variation revealed that these regions showed relationships among species of the diverse *benthic* clades of Malawi cichlids which were dramatically different from the rest of the genome (Fig. 2A, bottom panel, fig. S4). While in the rest of the genome PC1 tended to separate the three *benthic* subclades, in the focal regions three distinct clusters emerged that separated different groups of individuals and explained a much higher proportion of the genetic variance (fig. S5). Individuals in the intermediate cluster showed strongly increased heterozygosity as expected for the heterozygous state of two divergent haplotypes (fig. S6). An exception to this was chromosome 9, where only two clusters (one with increased heterozygosity) emerged, consistent with the absence of one homozygous state. Overall, with double crossover events in only two *deep benthic* individuals, the clustering was consistent with nearly complete recombination suppression between inverted and non-inverted haplotypes (fig. S7) as observed in other systems (*22, 37, 40*).

**Fig. 2:**
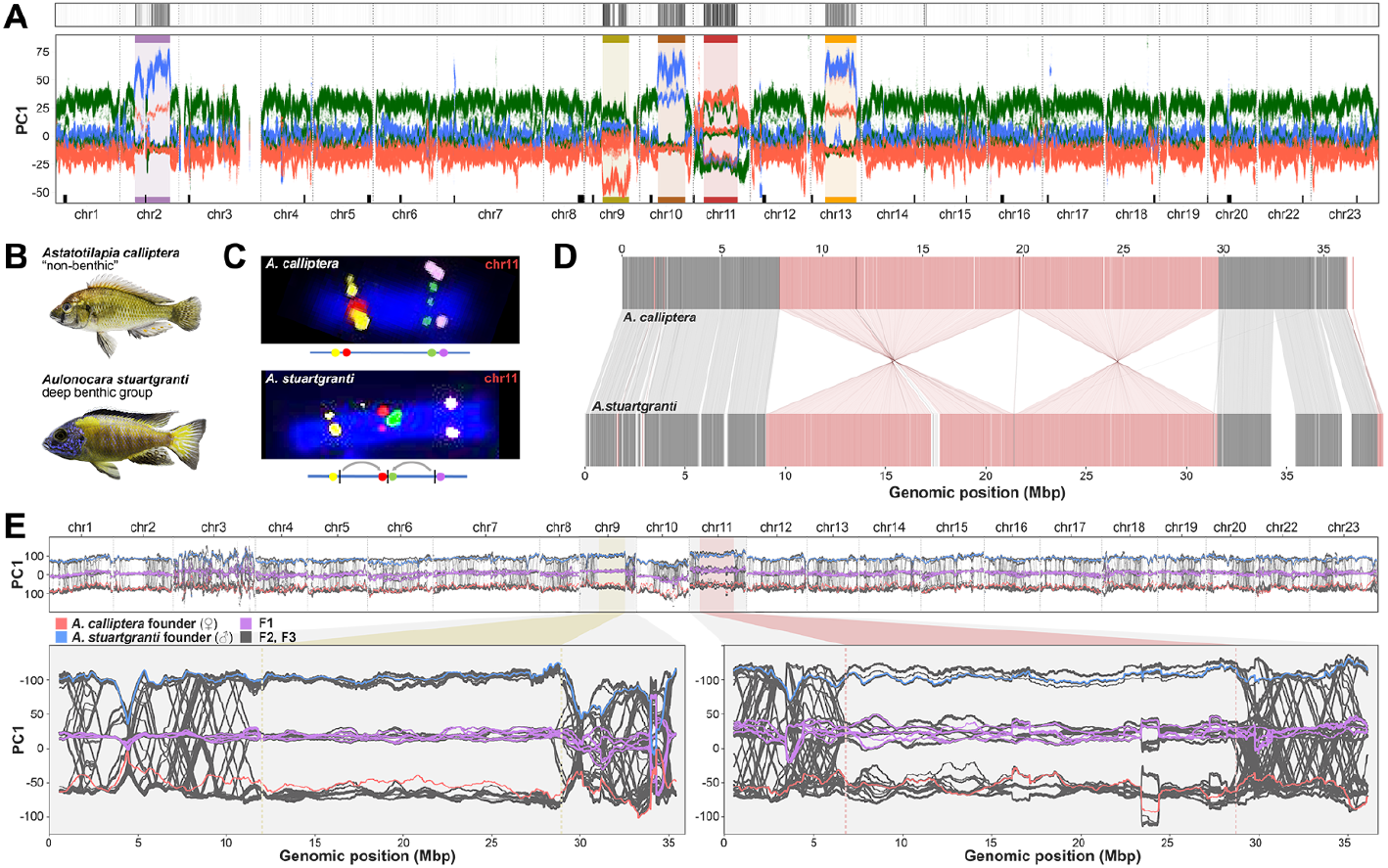
Characterisation of inversions. (**A**) (Top panel): Identification of genomic regions from clusters of aberrant phylogenetic patterns (see materials and methods). (Bottom panel): First genetic principal component in overlapping 1 Mbp windows along chromosomes, using the same colours for the *benthic* subclades as in Fig. 1. Outlier regions from the top panel are highlighted and colour-labelled. Centromeric satellite regions (for inference see materials and methods, text S1, fig. S17, table S6) are indicated as black rectangles on top of the X axis. (**B**) Representative photographs of the species used in panels C-D: *Astatotilapia calliptera*, a lineage of the Malawi radiation distinct from *benthics* from which the reference genome was produced, and *Aulonocara stuartgranti*, a species that genetically belongs to the *deep benthic* group, but lives in shallow rocky habitats (clade *Shallow rocky Aulonocara* in Fig. 1). According to WGS-typing, the species are expected to show opposite orientations for the chromosome 9 and 11 inversions. (**C**) Fluorescence in situ hybridisation (FISH) of markers on chromosome 11 left and right of the putative inversion breakpoints show the expected non-inverted orientation (upper panel) in *A. calliptera*. In *Au. stuartgranti* we see a double inversion (lower panel; see fig. S8 for FISH of chromosome 9). (**D**) Whole genome alignment of an ONT duplex long-read assembly of *Au. stuartgranti* to the *A. calliptera* reference assembly (which was re-scaffolded with chromosome conformation capture (Hi-C) data, see materials and methods) confirms the double inversion on chromosome 11 (for other chromosomes see fig. S9). (**E)** Top: Windowed PC1 values of whole genome sequenced founders and progeny of an interspecific cross. Among 290 F2 and F3 individuals no crossing-over events were observed in the inversion regions of chromosomes 9 (bottom left) and chromosome 11 (bottom right), while recombination was frequent in the flanking regions and on other chromosomes.

Using a combination of cytogenetics with long and linked read sequencing and *de novo* chromosome-level assembly of all five major clades of Malawi cichlids allowed us to confirm and characterise inversions in the regions on chromosomes 2, 9, 11 and 13 (but not chromosome 10 due to the lack of appropriate samples, see materials and methods) (Fig. 2B-C, table S4, fig. S8 to S19) and revealed additional smaller inversions that went undetected in the SNP analysis, including: (1) a small inversion nested inside the large inversion on chromosome 2, located next to the centromere (text S1, fig. S15 and (2) two adjacent inversions on chromosome 20 (text S1, fig. S16).

To confirm the suppression of recombination between inverted and non-inverted haplotypes, we performed an interspecific cross between *A. calliptera* and *Au. stuartgranti* and whole genome sequenced 290 individuals up to generation F3 (table S5). The absence of switching between the inversion-state clusters on chromosomes 9 and 11 on genomic PC1 axis in F2 and F3 individuals confirmed that recombination was fully suppressed in inversion regions of heterozygous F1s (Fig. 2E). Segregation ratios in F2s were Mendelian, except for the chromosome 11 inversion which had a moderate deficiency of homozygotes for the *A. calliptera* haplotype (genotype proportions 20:66:43; χ^2^ test on Mendelian ratios p=0.016).

### Inversions segregate within *benthic* subradiation

Next, we investigated the distribution of inversion states across the phylogeny based on a multi-step PC approach to infer inversion genotypes for all 1,375 sequenced individuals (“WGS-typing”) (Fig. 1, tables S7 and S8; fig. S20 and fig. S21), denoting as non-inverted or *ancestral* the orientation of the outgroup species *Pundamilia nyererei* (fig. S22) and *Oreochromis niloticus* (fig. S23). To further increase the number of genotyped individuals, we identified TE insertions highly correlated with inversion state and PCR-typed these insertions in an additional 401 individuals (see materials and methods, fig. S24, tables S9 to S11). Together, these analyses revealed that all specimens of the *mbuna* and *Rhamphochromis* subradiations and *A. calliptera* were fixed for the non-inverted, ancestral orientation for all five large inversions. All *Diplotaxodon* specimens also lacked the inversions on chromosomes 2, 10 and 13 but localized closer towards the cluster of inverted haplotypes than other non-*benthic* clades in the PCA-based typing of the chromosome 9 and 11 inversions. In our *de novo* assembly for *D. limnothrissa* both inversions are present (figs. S10, S25), suggesting that *Diplotaxodon* are fixed for the inverted chromosome 9 and 11 haplotypes.

Among the *benthic* clades, the five inversions showed strikingly different frequencies across the species: for chromosomes 2, 10, and 13, the inverted state was fixed or at high frequency in most *deep benthic* species, but almost absent among *shallow benthic* species and *utaka*. For chromosomes 9 and 11, the inverted states were fixed in most *benthics*, with the major exception of a large monophyletic subclade of *shallow benthics* in which chromosome 11 was mostly fixed for the ancestral non-inverted state and chromosome 9 mostly polymorphic. We will refer to this group as “*eukambuzi*”, inspired by the local name “Kambuzi” for some members of this group (Fig. 1). In summary, the distribution of inversion frequencies is consistent with a scenario in which inversions on chromosomes 2, 10, and 13 rose to high frequencies in an ancestor of the *deep benthic* lineage, while the inversions on chromosomes 9 and 11 rose to high frequency in the ancestors of two non-sister groups – pelagic *Diplotaxodon* and *benthics* – but with one monophyletic subgroup of the *benthics* (*eukambuzi*) retaining or re-gaining the non-inverted ancestral state.

### Origins and introgression patterns of the inversions

To better understand the evolutionary histories in inversion regions, we estimated genetic divergence times between Malawi cichlid species for the five inversion regions (both the inverted and non-inverted haplotypes) as well as for the remaining non-inverted regions of the genome (Fig. 3A, fig. S26). Surprisingly, we found that, outside the inversions, *benthics* were least divergent from *Diplotaxodon* and *A. calliptera* (top row in Fig. 3A), a pattern that is inconsistent with the inferred phylogenetic position of *benthics* as a sister group to *mbuna* and *A. calliptera* (e.g. ref. (*31*), Fig. 1, data S2), but rather suggests that *benthics* arose through admixture between the *Diplotaxodon* and *A. calliptera* lineages after their respective splits from *Rhamphochromis* and *mbuna* (Fig. 3B, see text S2 for more detailed discussion). Such a hybrid benthic origin model also provides a parsimonious explanation for the sharing of the chromosome 9, 11 and 20 inversions between all *Diplotaxodon* and most *benthics* (figs. S9 to S12) (as indicated by Ⓐ in Fig. 3B) and explains the general strong affinity of all inverted haplotypes to *Diplotaxodon* (Fig. 3A, right panels).

**Fig. 3:**
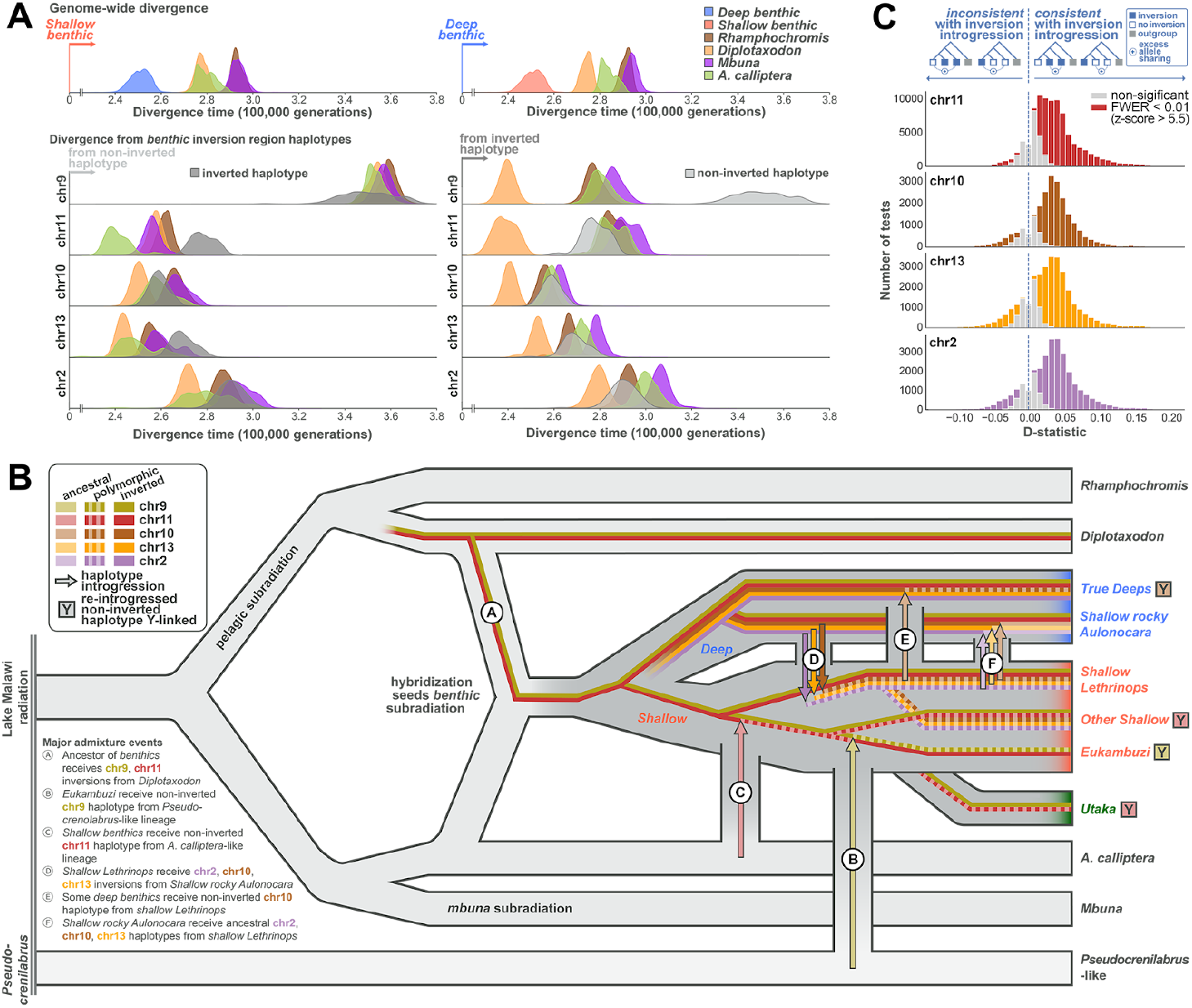
Evolutionary history scenario of inversion haplotypes. **(A**) Density plots of pairwise sequence divergence translated into divergence (coalescence) times assuming a mutation rate of 3×10^−9^ bp per generation (*31*). The top panels show results for the genome outside the five large inversions, comparing all major clades against *shallow benthics* (left) and *deep benthics* (right). Panels below the top row show divergence in inversion regions for the non-inverted (left) and inverted (right) *benthic* haplotypes. (**B**) A simplified model for the evolutionary history of the Malawi cichlid radiation, which includes several inversion haplotype transmission events. Vertical grey connections indicate major gene flow events. Letter-labelled arrows indicate transfer of inversion-region haplotypes. For further events see fig. S34. Lineages in which re-introgressed inversion region haplotypes of ancestral orientation apparently play a Y-like role in sex determination (see Fig. 5 and main text) are indicated by 🅈. (**C**) Evidence for transfer of inversion haplotypes through introgressive hybridisation. Histograms of ABBA-BABA statistics *D(P1, P2, P3, Outgroup*) calculated outside the inversions. For the different panels, we selected those ABBA-BABA tests for which the inverted state of the respective chromosome is present in one of the two more closely related species P1 and P2 but absent in the other and ordered them such that P2 shared the inversion state (presence/absence) with P3. In such a configuration, significantly positive values are suggestive of gene flow outside of inversions between the species sharing inversion states, while significantly negative values suggest gene flow between species not sharing inversion states. Under the null hypothesis of no inversion introgression, the statistic would be symmetric around zero.

The origins of non-inverted *benthic* haplotypes appear to be more diverse. The non-inverted *benthic* chromosome 11 haplotype is closest to *A. calliptera* (Fig. 3A, row 3), as expected if this haplotype was contributed from *A. calliptera* in the original founding of *benthics*. However, previously inferred signals of gene flow between *shallow benthics* and *A. calliptera* relative to *deep benthics (31*) and the relatively low heterozygosity of this haplotype (fig. S27), which is mostly present among *eukambuzi*, could alternatively point to its later introgression from *A. calliptera* (event Ⓒ in Fig. 3B).

For chromosome 9, *benthics* are almost fixed for the inversion with only some individuals, mainly *eukambuzi*, being heterozygous. However, it is striking that the non-inverted haplotype found in *benthics* is much more divergent from the rest of the Malawi radiation than any other inversion haplotype and the rest of the *benthic* genome (Fig. 3A, row 2). To follow this up, we produced a second SNP callset including a wide variety of related African cichlid species (“haplochromines”) and computed ABBA-BABA tests (*32, 41*) (text S2, fig. S28, tables S12 and S13, materials and methods). This revealed strong excess allele sharing of the non-inverted *benthic* chromosome 9 haplotype with *Pseudocrenilabrus philander*, one of the few outgroup species present today in the catchment of Lake Malawi (D = 0.45, block-jackknifing z-score 6.5; FWER corrected p = 4×10^−9^). We conclude from this that the chromosome 9 non-inverted *benthic* haplotype is not closely related to other Malawi haplotypes, but instead arrived in an ancestor of *eukambuzi* through admixture with a lineage containing *Pseudocrenilabrus*-like genetic material (Ⓑin Fig. 3B).

The remaining inversions on chromosomes 2, 10 and 13 are all common among *deep benthics*, and rare or absent among *shallow benthics* and *utaka*, suggesting that they rose to high frequency early in the deep *benthic* lineage (Fig. 3B). This is also consistent with the phylogenetic relationships within the inversion regions (figs. S29 to S33), which suggest that the few cases of *deep benthics* with non-inverted states and *shallow benthics* with inverted states are due to a limited number of later gene flow events, the most consequential of which are indicated in Fig. 3B (events Ⓓ-Ⓕ) (see text S2 and fig. S34 for a more comprehensive analysis). Most of these events transmitted more than one inversion haplotype and also genetic material outside the inversion regions (Fig. 3C) (*41*).

While there is evidence for each of the admixture and introgression events described in Fig. 3B and fig. S34, other more complex scenarios are also possible. Furthermore, additional minor introgression events and/or incomplete lineage sorting (ILS) are required to explain the final patterns of occurrence of the inversions. However, the alternative hypothesis of random segregation through incomplete lineage sorting giving rise to the observed patterns is not supported. First, inference based on coalescent calculations yields a probability of only 0.02% for retaining shared polymorphism among deep and shallow benthics at three inversions through ILS (materials and methods) (*28*). Second, an ABBA-BABA analysis confirmed that signatures of introgression outside of inversions were much more common when *deep benthic* and *shallow benthic* species shared their inversion state, compared to cases where they differed in inversion state (Fig. 3C), a pattern that is not expected under ILS.

### Inversion divergence between deep and shallow-living lineages

All five inverted states are found at higher frequencies in deepwater-living species (fig. S35, table S14). The corresponding regions show increased relative divergence and reduced cross-coalescence rates (a measure of genetic exchange) compared to the rest of the genome, and, for chromosomes 2, 10 and 13, also disproportionally high effect sizes in a population-structure-corrected genome-wide association study between *deep* and *shallow benthics* (materials and methods, Fig. 3A, figs. S36 to S40). Further, the inversion transmission events inferred above (Fig. 3B, fig. S34) were often such that species living at depth atypical for their clade (e.g., shallow-living rocky *Aulonocara* of the *deep benthic* clade and deep-living *shallow benthic* species like *Trematocranus* sp. ‘Cape Maclear’) had received the inversion haplotypes of species living at similar depths (fig. S35). Together, these observations support the hypothesis that inversion haplotypes contributed to divergence along a depth gradient.

### Pervasive signatures of adaptation on inversion haplotypes

Next, we investigated whether the evolution of inversion haplotypes was driven by adaptive processes and could potentially constitute “supergenes” of co-adapted alleles. To identify genetic variants relevant for the early evolution of inversion haplotypes, we computed correlation coefficients and significance scores (−log_10_ p-value) between SNP and inversion genotypes (figs. S41 and S42), where positive correlation coefficients correspond to derived SNP alleles on inverted haplotypes and negative coefficients to derived SNP alleles on non-inverted haplotypes. We expected the most highly inversion correlated SNPs (“ICS”) to contain variants relevant in early inversion evolution.

ICS were much more likely to be located in protein coding regions compared to other (non-ICS) SNPs on all five chromosomes (Fig. 4A) and showed a strong excess of non-synonymous divergence in McDonald–Kreitman type tests (fig. S43), indicating that positive and/or relaxed purifying selection contributed to inversion haplotype divergence. To confirm that there was a component of positive selection, and not just drift due to relaxed purifying selection as expected when a single haplotype rapidly raises in frequency (*42*), we calculated the normalised ratio of non-synonymous to synonymous mutations (dN/dS). While dN/dS ratios can approach 1.0 when selection is ineffective (complete relaxation of selection), values larger than 1.0 are only consistent with adaptive evolution (text S2) (*43*). We found that dN/dS ratios increased with increasing inversion correlation, with highly positive ICS showing dN/dS ≥ 1 for all five chromosomes (Fig. 4B, table S15). We confirmed with evolutionary simulations that such a pattern is only expected in the presence of substantial numbers of positively selected variants (text S2, figs. S44 and S45). Therefore, we conclude that widespread adaptive evolution contributed to diversification between ancestral and inverted haplotypes on all five chromosomes.

**Fig. 4:**
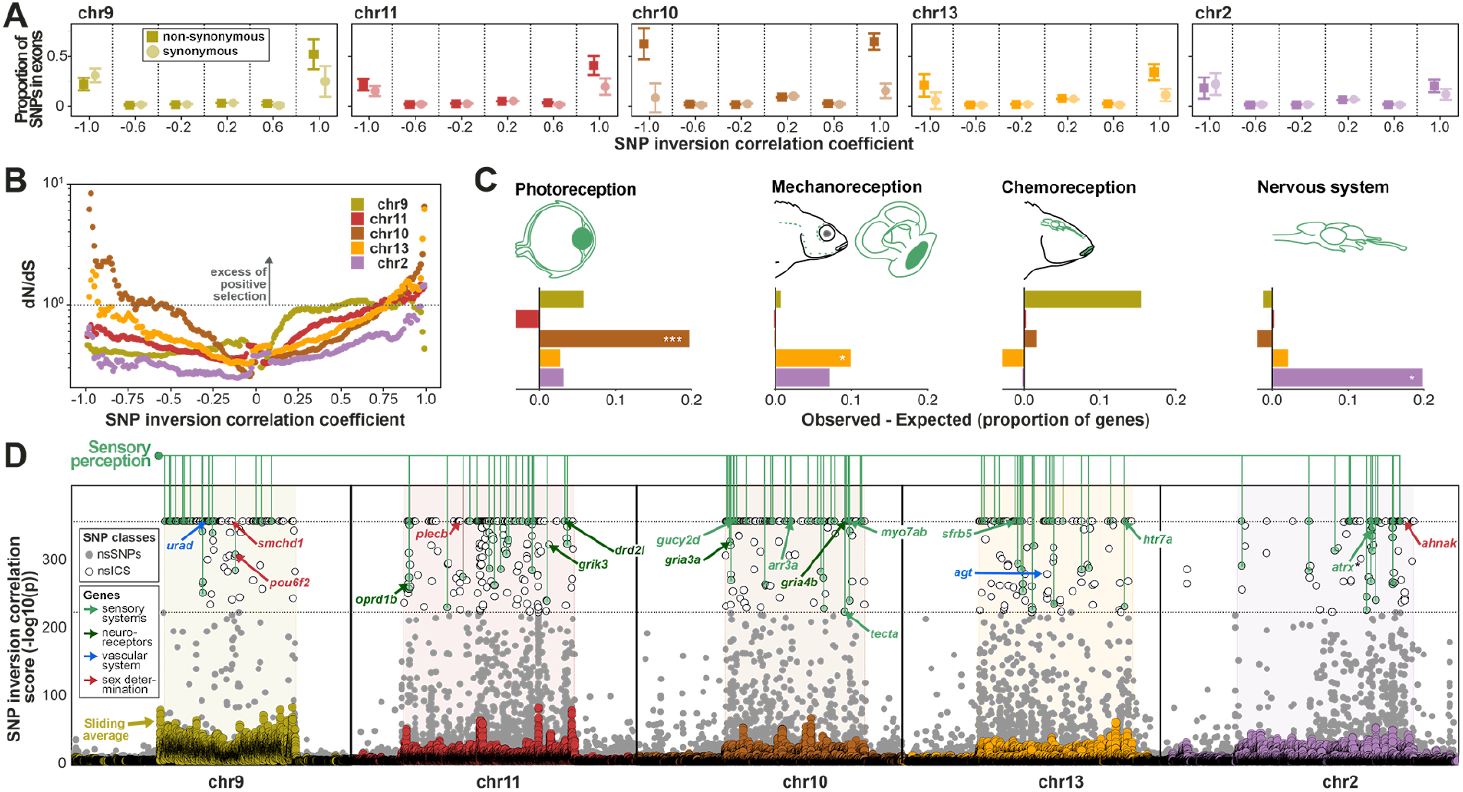
Adaptive evolution of inversion haplotypes. (**A**) Proportion of exonic SNPs, grouped by inversion correlation coefficient intervals, relative to all SNPs within the same interval. A positive correlation coefficient corresponds to the derived SNP allele being more common on the inverted haplotype, while a negative coefficient corresponds to the derived SNP allele being more common on the non-inverted haplotype (ancestral orientation). (**B**) dN/dS measured for SNPs as a function of inversion genotype correlation (see materials and methods). (**C**) Excess of genes containing non-synonymous highly inversion correlated SNPs (nsICS) among all genes highly expressed in the main sensory (green) and nervous system (blue). Expression data is based on the single-cell expression atlas of developing zebrafish Daniocell (*44*). The tissues were grouped into the functional categories: vision (eye), mechanoreception (lateral line, ear), chemoreception (taste, olfaction), and nervous system (neural). (**D)** Non-synonymous SNPs (grey dots; if ICS: empty dots) and averaged ICS scores in 100 SNP rolling windows (markers colour-coded according to inversion) on the five inversion chromosomes. We annotated nsICS located in genes with high expression in zebrafish tissue groups related to sensory perception (same as in (C)). nsICS in candidate genes discussed in the main text are annotated with arrows (if several were located in the same gene, the highest nsICS is annotated) and color-coded by functional category (table S24).

Interestingly, despite their general high differentiation, for many ICS both alleles were present in at least one copy on both inverted and non-inverted haplotypes (37-72% of ICS, table S16). As expected, this pattern of shared polymorphism across inversion orientations is even more apparent for other (non-ICS) common variants (table S16) and suggests that despite their excess divergence, recombination between inversion region haplotypes is not uncommon at evolutionary timescales, potentially providing a mechanism to concentrate adaptive alleles on such haplotypes.

### Inversions contribute to sensory and physiological adaptation

To study functional roles of genes involved in inversion adaptations, we first analyzed expression of the 315 genes from inversion regions with non-synonymous ICS (nsICS) in a multi-tissue single-cell gene expression atlas of the zebrafish model (Daniocell database (*44*), text S3). We found that individual inversions showed elevated expression in tissues related to vision, mechanoreception, and the nervous system (FDR-corrected p = 2×10^−4^, 0.02, 0.04, respectively) (Fig 4C, figs. S46 and S47, tables S17 to S20). As most of the associations were related to neural and sensory tissues, we checked for overrepresentation of all neural and sensory tissues across all five inversions, and found them to be significant (FDR-corrected p = 6×10^−4^, fig. S48, tables S21 and S22). We additionally tested for gene ontology (GO) enrichment of genes near ICS (see materials and methods) and found sensory system-related categories in all five inversions (table S23). Finally, we found vascular system-related categories functionally linked to responses to hypoxia stress to be enriched in three inversions. All these findings are consistent with adaptations related to changes in light, oxygen, and hydrostatic pressure along a depth gradient, as observed in many aquatic organisms (*45*–*47*) including cichlids (*31, 48, 49*).

Among the 83 genes with two or more nsICS were strong candidate genes for depth adaptation (tables S24 and S25), including genes involved in signal transduction in photoreceptor cells (*arr3a, gucy2d, pou6f2*), otolith tethering (*tecta*), sound perception (*myo7ab*), kidney function and blood pressure regulation (*urad*) and a master regulator of vasoconstriction (*agt*) (Fig. 4D, text S4, table S24). Interestingly, some genes showed a close similarity between the amino acid sequence coded by the inverted haplotype and that of *Diplotaxodon*, even when the relevant inversion was not present in *Diplotaxodon* (e.g., *arr3a* on chromosome 10, see also Malinsky et al. (*31*)) (fig. S49), which is consistent with a hybrid ancestry of *benthics* and subsequent differential selection between recombination-suppressed inversion haplotypes. Intriguingly, several genes harbour both positive and negative ICS, as expected under diversifying selection.

Consistent with their enrichment among sensory and neural tissues, we also found nsICS to be significantly overrepresented among neuroreceptor genes that have been previously associated with social (glutamate) and affiliative (oxytocin and arginine vasopressin/vasotocin, opioid receptors, dopamine, serotonin) behaviour (*50*) (6 out of the 46 candidate genes, Fisher’s exact test p=0.0011) (table S26). These are three glutamate receptors (*gria3a* and *gria4b* on chromosome 10, *grik3*, on chromosome 11), one opioid receptor (*oprd1b* on chromosome 11), one dopamine receptor (*drd2l* on chromosome 11), and one serotonin receptor (*htr7a* on chromosome 13) (fig. S49).

It is notable that the identified neurotransmitters are not only associated with fish social behaviour in general, but have been specifically associated with bower building behaviour in Malawi cichlids (*51*), a behavioural phenotype important for assortative mating (*52*), which has previously been linked to the existence of supergenes (*50*) and associated with genetic divergence peaks inside our chromosome 2 and 11 inversion regions (*34*). Following this up, we found no significant correlation between bower type and the presence of the five inversions when accounting for phylogeny (materials and methods; fig. S50, table S27; but see text S1), suggesting that previously detected associations might be due to phylogenetic confounding.

Overall, our selection analyses suggest that widespread functional divergence in genes related to sensory, vascular, and nervous systems occurred during the early evolution of inversion haplotypes.

### Inversions contribute to sex determination

Considering segregation patterns of inversion genotypes within species, we observed a notable excess of inversion heterozygotes for chromosomes 9, 10, and 11 (deviation from within-species Hardy-Weinberg-equilibrium, HWE, p < 10^−4^, 0.0048, and 0.0133, respectively). This pattern was most extreme for chromosome 9, for which despite the presence of 77 heterozygous individuals across twelve species, not a single homozygous ancestral (non-inverted) state was present in any *benthic*.

Since inversions are a common feature in the evolution of suppressed recombination on sex chromosomes (*53, 54*), we hypothesized that the observed excess of heterozygotes could be due to sex-linked inheritance (fig. S51). In the two species for which we had gonad-examination-based sex assignment, we found a perfect correlation of sex with chromosome 11 inversion state in *Copadichromis chrysonotus* (Fig. 5A, B; n=28, Fisher’s exact test p-value = 4.7×10^−8^), while the other species, *Copadichromis mloto*, was not variable for any inversion. We further confirmed a significant chromosome 11 inversion–sex association among 107 laboratory-bred individuals from 11 broods of three species (Fig. 5C-E). Notably, in a second laboratory population of one of these species (*O. tetrastigma*) with different geographic origin, the chromosome 11 inversion was fixed for the inverted state, but there was a correlation of sex with the chromosome 9 inversion state (Fig. 5E). This is consistent with previous observations of multiple sex determination systems acting even within single Malawi cichlid species (*55, 56*). In each case where there was an association, males tended to be heterozygous and females homozygous for the respective inverted state as expected for XY-like sex determination systems.

**Fig. 5:**
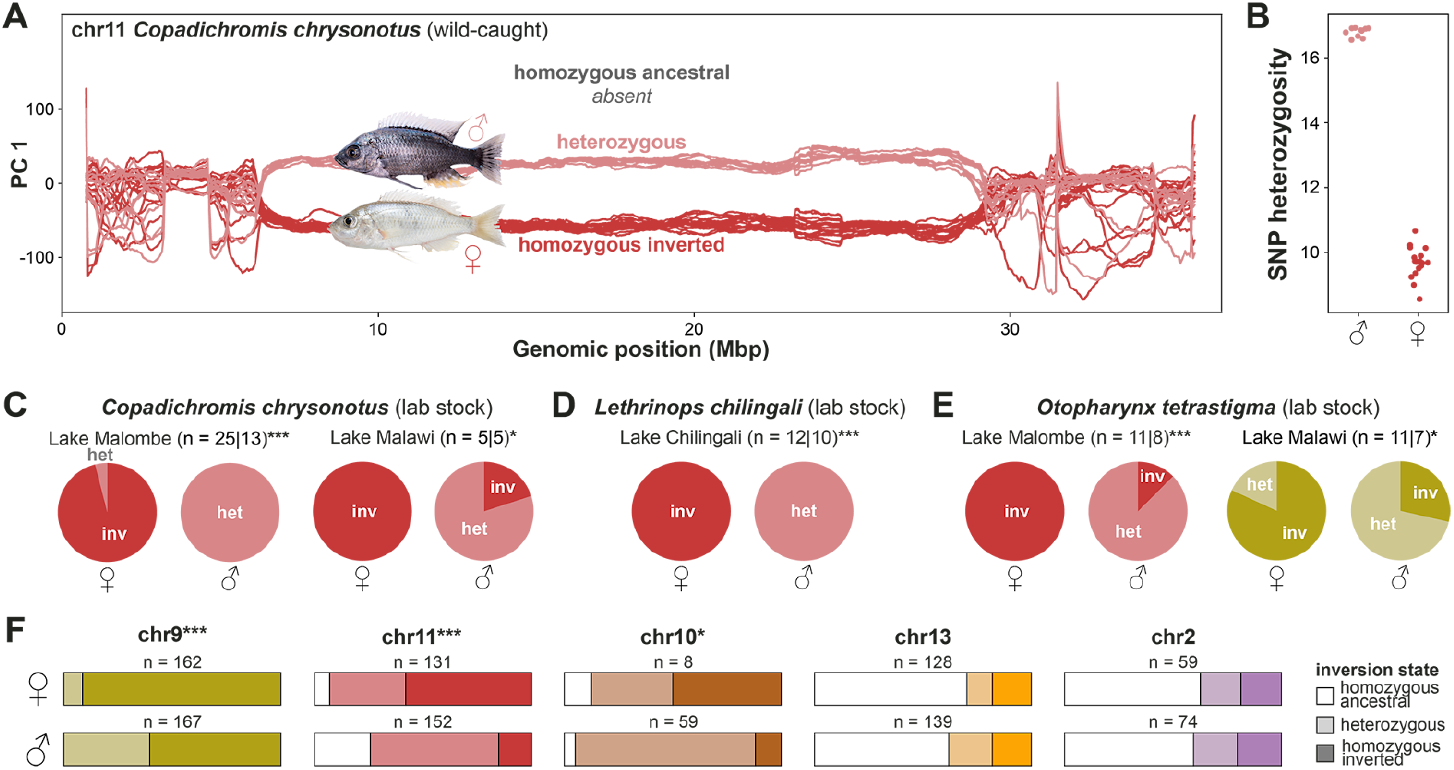
Sex association of inversions. (**A**) Windowed PC analysis along chromosome 11 demonstrates perfect association of inversion genotype with sex in our sample of 28 wild-caught *Copadichromis chrysonotus*. (**B**) SNP heterozygosity among our sample of wild-caught male and female *C. chrysonotus*, measured as number of het SNPs per 10 kbp. (**C**-**E**) Sex-inversion associations in lab-raised populations. Per population, the number of females and males is given (separated by ‘|’) and asterisks denote significance levels of Fisher’s exact tests of inversion genotype–sex correlation (*: p < 0.05, ***: p < 0.001). Inversion genotype per sex (confirmed through gonad examination) in lab-raised broods of (**C**) *C. chrysonotus* from Lake Malombe (p < 0.001, left) and from Lake Malawi (p = 0.048, right), (**D**) *Lethrinops chilingali* from satellite lake Chilingali (p < 0.001) (**E**), *Otopharynx tetrastigma* from Lake Malombe at the outflow of Lake Malawi (p < 0.001, left) and from the northern part of Lake Malawi (p = 0.049, right). (**F**) Proportions of homozygous/heterozygous inversion genotypes in males and females of species with heterozygotes present, according to WGS and PCR typing of 809 samples (67 species). Asterisks denote significance levels of Fisher’s exact tests of inversion genotype–sex correlation (*: p < 0.05, ***: p < 0.001).

To further examine the extent of sex-linkage of inversions, we pooled data for each inversion across species with at least one heterozygous sample (table S28). This revealed that, while females tended to be homozygous for the inverted state, there was a significant association of male sex with the heterozygous state for chromosomes 9, 10, and 11 (Fisher’s exact test p-value = 3.7×10^−11^, 0.014, and 5.3×10^−11^, respectively) (Fig. 5F, table S28) consistent with a widespread role of the inversions in sex determination. That said, many species were not polymorphic for inversions and even within polymorphic species associations were usually not perfect (data S3) suggesting additional genetic or environmental effects contributing to sex determination and a rapid turnover of sex-linked function.

Given that the evolution of sex-determining regions often involves changes in gene expression between male and female haplotypes, we obtained transcriptomic data of five tissues (muscle, liver, brain, gills, gonads) for 11 males of *C. chrysonotus*, the species in which males were heterozygous for the chromosome 11 inversion while females were fixed for the inverted state, and investigated allele specific expression (ASE). Among genes with significant ASE (fig. S52, text S3) there was a moderate bias towards lower expression of the Y-like non-inverted haplotype, a pattern seen in many organisms (*57*). Further, lacking access to appropriate female samples, we obtained equivalent data for eleven male *C. mloto*, the congeneric species fixed for the derived chromosome 11 inversion state, to perform differential gene expression (DE) analysis between the two inversion states (fig. S53). Several of the significant ASE and DE genes (FDR < 0.05) were implicated in sex determination, sex specific expression or gonad function in other (fish) species (table S24) and ICS were significantly overrepresented among genes with significant allele specific expression (Fisher’s exact test p = 1.6×10^−7^). Furthermore, we found candidate genes related to sex and reproduction among the strongest candidate genes for adaptive evolution (i.e., those with the largest number nsICS) (Fig. 4D; text S3).

It is notable that for each of the three inversions with evidence for sex-linkage, the Y-like, non-inverted haplotype arrived in the affected *benthics* through introgression events, affecting mainly *eukambuzi*, other *shallow benthics/utaka*, and *deep benthics*, for chromosomes 9, 11, and 10, respectively (events Ⓑ, Ⓒ, and Ⓔ in Fig. 3B). For chromosome 9, this is further supported-by results from a lab based hybrid cross between females of *A. calliptera* and males of the *eukambuzi Protomelas taeniolatus, which* showed a QTL peak for sex in the chromosome 9 inversion region (*58*). This is consistent with a dominant male-determining function of the non-inverted *benthic* haplotype (with origin external to the Malawi radiation), even when paired with an *A. calliptera* haplotype of the same orientation.

To further investigate the generality of the role of inversions in sex determination of haplochromine cichlids, we applied our SNP-based inversion detection approach to publicly available sequencing data for the Lake Victoria adaptive radiation (*59*). The results suggest that the sex-linked regions identified by Feller et al. (*58*) on chromosomes 9 and 23 are in fact chromosomal inversions (the one on chromosome 9 being distinct from the one present in Malawi) (figs. S22 and S54), pointing to a wider relevance of chromosomal inversions in cichlid sex determination.

## Discussion

In this article we identify large chromosomal inversions present in the Lake Malawi cichlid radiation and present evidence that their evolutionary history was shaped by introgression, ecological adaptation, and the turnover of sex determination systems. Our results are consistent with a recent preprint that independently identified the inversions described here and their sex-linkage based on optical mapping and chromosome-level de novo assemblies (*60*). Given the evolutionarily independent presence of sex-linked inversions in Lake Victoria cichlids (fig. S54), which are potentially also involved in adaptive introgression (*59*), and large scale differences in male/female DNA sequence in Lake Tanganyika cichlids (*61*), we suggest that such rearrangements may be a common feature of adaptive radiation in cichlids.

Chromosomal inversions have been implicated in adaptation, sex determination, and speciation in many systems, especially in the context of adaptive divergence with gene flow (*7, 11, 15*–*22, 28*) likely because of their ability to lock together adaptive alleles (*7*). The chromosomal inversions we identified in Malawi cichlids were involved in gene flow events at different stages of the radiation, most prominently a founding admixture event of the species rich *benthic* clade and introgression from a distantly related lineage outside the Malawi radiation (chromosome 9 inversion). These events coincide with bursts of eco-morphological diversification of the resulting lineages. In the latter case, this concerns the *eukambuzi*, which show exceptional diversity in eco-morphology, body patterning, and colouration (*30*).

We found evidence for inversion transmission between *deep* and *shallow benthic* species caught at similar depths (fig. S35). At the same time, when not introgressed, inversions seemingly helped to suppress gene flow and thereby contributed to adaptive divergence. However, despite their excess divergence compared to the rest of the genome and the complete recombination suppression we observed in a cross (Fig. 2E), most common genetic polymorphism is shared across orientations suggesting that inversions are a barrier to genetic exchange rather than completely suppressing it. On the one hand, this facilitates the purging of genetic load, which often hinders the spread of inversions (*62*), while on the other hand, it provides an additional mechanism for the creation of combinatorial diversity (*5*).

The genetic variants most differentiated between inverted and non-inverted haplotypes (ICS) show a strong relative excess of amino acid changing mutations, as expected only under adaptive evolution at many loci (text S3). These mutations showed enrichment in genes related to and expressed in tissues involved in sensory and behavioural functions. This makes sense, because sensory systems mediate sound perception, mechanoreception and vision, essential for navigation and feeding in fishes (*63*), making them important targets of ecological adaptation to differing underwater environments (*64*). Although further experiments – most promisingly within species polymorphic for inversions – will be necessary to dissect the precise phenotypes of the adaptive alleles within inversion regions, our results point towards widespread, multigenic adaptation along a depth gradient which is a frequent axis of differentiation in fishes (*65, 66*).

We found evidence for XY-like sex linkage of the inversions on chromosomes 9, 10, and 11, in which introgressed haplotypes of ancestral orientation act as Y chromosomes in some extant species, with inversion-region genes related to sex and reproduction being under allele specific expression in XY males and showing signatures of selection. Consistent with the highly dynamic nature of sex determination in many fishes (*67*) and specifically cichlids (*58, 61, 68*), our results point to a relatively easy recruitment of sex determination loci (SDLs), possibly as a direct consequence of introgression of relatively divergent haplotypes affecting a sex determination threshold, or due to heterozygote advantage of introgressed inversions selecting for the recruitment of SDLs (*69*).

Sexual selection has been identified as a major predictor of successful radiation in cichlids (*70*), and assortative mating is a main driver of cichlid reproductive isolation (*71*). Both of these processes rely heavily on the same sensory systems (e.g., vision (*71*), olfaction (*72*), and hearing (*73*)) that are also relevant for adaptation to depth and feeding niches, and that we identified as candidates for adaptive evolution. Assortative mating and the evolution of sex linked regions are both forms of sex-specific selection (*74*). Although the interplay between these forms of selection is not well understood, it can give rise to synergistic evolutionary dynamics, potentially mediated by sexually antagonistic selection (*75, 76*).

While our analysis focused on genetic variants nearly fixed between inversion haplotypes, we expect that additional variants specific to particular species groups will prove important for further diversification. Furthermore, alongside the inversions that identify admixture events, we also see a signal of introgressed material in the rest of the genome, providing a potential substrate for further selection and adaptation. Indeed, while we focussed on large inversions segregating across many species, there are expected to be many more inversions and other structural genetic variants that are smaller or have a more limited taxonomic distribution, as suggested by our whole genome alignments and in Talbi et al. 2024 (*77*). Surely, more structural variants with relevance in adaptive diversification are to be found in future studies.

In conclusion, the haplotypes of five chromosomal scale inversions in the Malawi cichlid adaptive radiation show supergene-like signs of adaptive evolution and repeated introgression associated with speciation. Together with the repeated transient sex-linked nature of introgressed haplotypes, this provides a substrate for rich evolutionary dynamics around the interactions between natural, sexual, and sexually antagonistic selection.

## Supporting information

Supplmentary tables in .tsv format

## Acknowledgments

We thank the Department of Fisheries of the Government of Malawi for facilitating Malawi cichlid specimen collection and Martin Genner, Alix Tyers, Mingliu Du, Mexford Mulumpwa, Magnus Nordborg, Karl Svardal and the staff of the Monkey Bay Fisheries Research Unit for support with sampling, Shane McCarthy for bioinformatics support, Walter Salzburger for access to facilities and data, and Patrick Gemmell for input on the project. We are grateful to Daniel Shaw, Regev Schweiger, Victoria Caudill, Peter Ralph, Jonas Lescroart, and Els De Keyzer for useful comments on the manuscript.

## Authors contributions

H.S. and R.D. conceived the study, with support from L.M.B. H.S. organised cichlid collections in Malawi with support from B.R., R.Z., W.S., J.C.G, M.N., M.M., R.D., G.V., E.A.M., and G.F.T. G.F.T. undertook final taxonomic assignment. I.B. and V.B. performed DNA extractions. T.L. produced the original variant callset. H.S., L.M.B., V.B., I.A., J.S., and J.C.G. analysed the SNP data. Inversion detection was accomplished by L.M.B. (PCA-based) and I.A. (clusterization, phylogenetic reconstructions). H.S. performed hybridisation analysis. V.B. made selection tests (together with H.S.), zebrafish-based functional analysis and candidate genes investigation. J.C.G. performed enrichment analysis and candidate gene investigation. J.S. and H.S. performed population history reconstructions. J.E., A.H., M.E.S., and G.F.T. performed the species cross. B.F. designed the PCR essays and provided molecular lab support. V.B. performed PCR typing. C.Z., L.M.B. and B.F. produced assemblies and analysed long read and Hi-C data with support from J.S. F.C.J. performed and analysed time-foward simulations. A.H.H. raised and collected lab populations for sex determination analysis. N.H. performed PCR-typing of lab populations. L.M.B. detected sex-linked inversions in the Lake Victoria radiation. M.K. and Y.F.C. performed haplotagging. I.A. analysed haplotagging data. W.S., I.A., and H.S. analysed RNA-seq data. S.L. and F.Y. performed FISH. S.G. produced and analysed the outgroup variant callset with support from H.S. M.M. produced Hi-C data and the ancestral sequence. D.A.J. contributed the *Diplotaxodon* long read samples. A.B. provided support for sex chromosome and adaptation related analyses. L.M.B. designed the main figures, with input from H.S., V.B., J.C. and I.A. H.S. wrote the initial manuscript. H.S., R.D., L.M.B, V.B., and J.C. edited the manuscript with contributions from all authors. All authors read and approved the final manuscript.

## Data and materials availability

Supporting data is made available in the online supplementary on an open-access basis for research use only. ENA Accession numbers of raw sequencing data *<will be added to Supplementary Table 1 before final publication>*. VCF are shared in Dryad repository <*DOI added before final publication*>. Data was collected under appropriate ethical and sampling permits and genetic material and sequences are subject to an Access and Benefits Sharing (ABS) agreement with the Government of Malawi. Any person who wishes to use this data for any form of commercial purpose must first enter into a commercial licensing and benefit-sharing arrangement with the Government of Malawi.

## Supplementary Materials

Materials and Methods

Supplementary Text

Fig. S1 to Fig. S54

Table S1 to S31

References (78-177)

Data S1 to S8

