## Supplementary methods, text, figures, tables for "Introgression dynamics of sex-linked chromosomal inversions shape the Malawi cichlid adaptive radiation"

**The PDF file includes:**

Materials and Methods
Supplementary Text
Fig. S1 to Fig. S54
Tables S1 to S31
References 78 to 177

**Other Supplementary Materials for this manuscript include the following:**

Data S1 to S8

### List of contents

**Introgression dynamics of sex-linked chromosomal inversions shape the Malawi cichlid adaptive radiation...1**
Abstract...2
Main...2
Large inversions suppress recombination...3
Inversions segregate within benthic subradiation...6
Origins and introgression patterns of the inversions...8
Inversion divergence between deep and shallow-living lineages...9
Pervasive signatures of adaptation on inversion haplotypes...11
Inversions contribute to sensory and physiological adaptation...13
Inversions contribute to sex determination...14
Discussion...17
References and Notes...19
Acknowledgments...25
Funding...25
Authors contributions...25
Competing interests...26
Data and materials availability...26
Supplementary Materials...26
**List of contents...2**
**Materials and Methods...6**
Sample collection for short-read sequencing...6
Short read sequencing, alignment, SNP detection, and statistical phasing...6
Ancestral state inference...7
Phylogenetic reconstruction...8
Clustering-based inversion detection...8
Windowed principal component analysis (PCA)...9
Windowed heterozygosity...9
Genome assembly...9
Comparative genomics analysis...11
Cichlid cell lines establishment and FISH...11
Linked read analysis (haplotagging)...12
Species cross...12
Centromere localization in the reference genome...12
Structural variants and transposable element scan...13
Breakpoint estimation...14
Inversion genotyping of whole genome-sequenced samples...14
PCR assay to genotype inversions...15
Test for structured Hardy-Weinberg equilibrium...16
Sex association...16
Lake Victoria radiation inversion scan...16
Inversion haplotype rephasing...17
Haplotype-resolved local phylogenetic inference in inversion region...17
Pairwise differences and divergence times...17
Investigation of chromosome 9 haplotype origin...18

Inversion-wide heterozygosity ... 19
Probability of shared polymorphism ... 19
Gene flow analysis ... 20
Principal component analysis of benthic samples ... 20
Habitat depth and inversion state ... 20
Fixation index (FST) population branch statistic (PBS) in windows ... 21
MSMC analysis ... 21
Genome-wide association analysis ... 22
SNP–inversion-correlation ... 22
Simulations of selection on inversion haplotypes ... 22
Gene ontology enrichment analysis ... 24
Selection tests (NI, DoS, dN/dS) ... 24
Expression patterns in zebrafish homologs ... 24
Association between bower building and inversion state ... 25
Allele-specific expression analysis ... 26
RNAseq and differential gene expression ... 26
Gene haplotype analysis ... 26
**Supplementary text ... 28**
Text S1: Comparative genomics and small inversions ... 28
Text S2: Evolutionary history of inversions ... 30
Text S3: Inversion-correlated SNPs (ICS) and selection ... 32
Text S4: Signals of adaptation in inversions ... 35
**Figs. S1 to S54 ... 40**
Fig. S1: Representative photos used in Fig. 1 ... 41
Fig. S2: Clustering of distance matrices ... 42
Fig. S3: Contiguous runs of cluster identity ... 43
Fig. S4: Clade-wise windowed PC analysis ... 44
Fig. S5: Variance explained by PC1 ... 45
Fig. S6: Windowed SNP heterozygosity ... 46
Fig. S7: Double recombination events in wild samples ... 47
Fig. S8: Fluorescence in situ hybridization results for chromosome 9 ... 48
Fig. S9: Structural rearrangements between Astatotilapia calliptera and Aulonocara stuartgranti ... 49
Fig. S10: Structural rearrangements between Astatotilapia calliptera and Diplotaxodon limnothrissa ... 50
Fig. S11: Structural rearrangements between Astatotilapia calliptera and Rhamphochromis sp. 'Chilingali' ... 51
Fig. S12: Structural rearrangements between Astatotilapia calliptera and Tropheops sp. 'mauve' ... 52
Fig. S13: Hi-C maps of the chromosome-level genome assemblies ... 53
Fig. S14: Proportion of tags shared between breakpoints ... 54
Fig. S15: Nested inversion on chromosome 2 ... 55
Fig. S16: PCA of chromosome 20 ... 56
Fig. S17: Centromere identification in the reference genome ... 57
Fig. S18: Differences in TE length between inverted and non-inverted haplotypes ... 58
Fig. S19: Transposable element content along inversion chromosomes ... 59
Fig. S20 Inversion breakpoint estimation ... 60
Fig. S21: PCA-based genotyping of inversion state ... 61
Fig. S22: Structural rearrangements between Astatotilapia calliptera and Pundamilia nyererei ... 62
Fig. S23: Structural rearrangements between Astatotilapia calliptera and Oreochromis niloticus ... 63
Fig. S24: PCR-based inversion genotyping ... 64

### Materials and Methods

#### Sample collection for short-read sequencing

Of the 1,375 Malawi cichlid individuals in the short-read whole genome sequencing dataset, 118 samples were already sequenced as part of our earlier study (*31*), while the majority of remaining samples derive from specimens collected in the wild in Malawi during field trips in 2016 and 2017. Four samples were acquired in the UK through the aquarium trade. The 20 *Astatotilapia calliptera* from Lake Kingiri and some *Maylandia fainzilberi* samples were kindly shared by Martin Genner. Specimens were collected either by chasing them into small-meshed monofilament gill nets during scuba-diving or snorkelling, purchased from local fishermen, or by bottom trawling using the RV Ndunduma research vessel of the Malawian Department of Fisheries. Before tissue acquisition, live samples were euthanized using clove oil before collecting fin clips. Specimens were photographed. Fin clips were preserved in 100% (dehydrated) ethanol in individually labelled sample tubes and stored at -80 °C. For detailed per-sample information see table S1.

#### Short read sequencing, alignment, SNP detection, and statistical phasing

DNA was extracted from ethanol-preserved fin tissue using the Wellcome Sanger Institute’s automated DNA extraction pipeline or manual DNA extraction using standard protocols. Whole-genome sequencing libraries were prepared using the Wellcome Sanger Institute's standard IHTP WGS NEB UltraII library process. Specifically, samples were quantified with Biotium Accuclear Ultra high sensitivity dsDNA Quantitative kit using Mosquito LV liquid platform, Bravo WS and BMG FLUOstar Omega plate reader and cherry picked to 200ng / 120ul using Tecan liquid handling platform. Cherry Picked plates were sheared to 450bp using a Covaris LE220 instrument. Post sheared samples were purified using Agencourt AMPure XP SPRI beads on Agilent Bravo WS. Libraries were constructed using ‘NEB Ultra II custom kit’ on an Agilent Bravo WS automation system. PCRs were set-up using KapaHiFi Hot start mix and IDT UDI 96 PCR (sets A-D) barcodes on Agilent Bravo WS automation system. Post PCR plates were purified using Agencourt AMPure XP SPRI beads on a Beckman BioMek NX96 liquid handling platform. Libraries were quantified with Biotium Accuclear Ultra high sensitivity dsDNA Quantitative kit using Mosquito LV liquid handling platform, Bravo WS and BMG FLUOstar Omega plate reader, pooled on a Beckman BioMek NX-8 liquid handling platform and normalised. Libraries were sequenced on Illumina HiSeq platforms (100-150 bp paired-end) to genomic coverages of 5.6–48.3 fold (median 17.3 fold; for per-sample information see table S1).

The samples were aligned to the reference genome fAstCal1.2 (RefSeq: GCF_900246225.1) using BWA-MEM (version 0.7.17) (*78*) (command: bwa mem) and then sorted and converted to CRAM format using samtools sort (*79*) and then choosing -C option to output as a CRAM file (*80*). We identified genetic variants segregating among all 2,198 individuals with BCFtools (version 1.12) (*79*) by first using the mpileup function to calculate genotyping likelihoods for each individual based on the subset of all primary mapped, non-PCR duplicate reads with no QC fail bits set and with minimum Phred-scaled base and map qualities of 20 and 13, respectively. We used the genotype likelihoods in the BCFtools multiallelic variant calling model (BCFtools call -m) to call variants jointly across all individuals, specifying a heterozygosity prior (-P) of 0.001 and a Hardy-Weinberg equilibrium prior for genotype frequencies (-G) within, but not across, major clades (*Astatotilapia calliptera*, *deep benthics*, *shallow benthics*, *mbuna*, *Diplotaxodon, Rhamphochromis*, and *utaka*) to balance sensitivity for identifying rare variants with robustness to noise. We required that individuals have at least two gapped reads (and that this was at least 5% of their total reads at a site) to consider candidate indels (mpileup -p -m 2 -F 0.05). We left-aligned and normalized indels with respect to the fAstCal1.2 reference genome assembly using BCFtools norm. We performed tests of excess heterozygosity across all samples and calculated the number of samples with data at each site using BCFtools fill-tags, and added the resulting values to the VCF INFO field. We annotated the VCF FILTER field with genomic accessibility information which we used to mask regions of the genome containing sites refractory to short read sequencing and mapping from downstream analyses. Specifically, for a subset of 255 individuals representing the specimen sequenced to the highest depth of coverage for unique taxa (herein referred to as the QC subset), we calculated quality metrics related to depth, base accuracy, and mapping using all sequencing reads contained in their BAM files with the program bamstats (*81*). We added flags indicating that a site failed quality control if the total depth (based on all sequence reads) was outside of the genome-wide 95 interpercentile range for the QC subset, the root mean square (RMS) map quality was below 35, the fraction of reads with either map quality and/or base quality of zero exceeded 2x the genome-wide average for the QC subset, the RMS base quality was below 20, or no reads covered the site. In addition to quality control based on the QC subset of individuals, we also flagged and filtered sites for which the total site depth considering only high-quality reads used in genotyping (VCF DP) was more extreme than the genome-wide median +/- 25% for all samples used for variant calling. Lastly, we extracted all variants and the subset of biallelic SNPs for downstream analyses from a VCF containing all sites, denoting SNPs within deletions (VCF FILTER/VT) using the extractVariants.pl and extractBiallelicSnps.pl scripts available on github (*82*). Finally, the biallellic SNP set was phased using Eagle (version 2.4) (*83*) with parameters set as --Kpbwt=40000, --expectIBDcM=0.5 to suit the cichlid model and missing genotypes were not imputed. Unless stated otherwise, only biallelic SNPs on the that passed all filter criteria were used for further analysis. Filtering resulted in an accessible genome size of 666,872,383. To annotate the SNPs in our vcf we used SnpEff (version 5.0) (*84*). A custom SnpEff database for the fAstCal1.2 reference was built using the GTF (release version 99) available on the Ensemble genome database (command: snpEff.jar build -gtf22). After which the vcf was annotated using appropriate SnpEff command (snpEff -Xmx16g fAstCal1.2.99 input.vcf > annotated.vcf). VCFs are accessible at Dryad <added_prior_to_publication> (see *Data availability* for ABS restrictions).

#### Ancestral state inference

We used whole genome alignments between the *A. calliptera* fAstCal1.2, the *Cyphotilapia frontosa* fCypFro1.1 (NCBI REF<added_prior_to_publication>), and the *Oreochromis niloticus* UMD/NMBU (NCBI GenBank: GCA_001858045.3) reference genomes to probabilistically reconstruct the ancestral sequence at the *A. calliptera* and *C. frontosa* divergence using the phyloFit and prequel tools from the PHAST software package (*85*). This sequence provides a good basis for polarising ancestral vs. derived alleles in our dataset because *A. calliptera* and *C. frontosa* diverged about 8.7 Mya, preceding the first split among the haplochromine species we are considering in this study, which was estimated to be 7.1 Mya (*86*). The whole genome alignments were generated following the UCSC paradigm (*87*), using lastz

(*88*) (version 1.02, “B = 2 C = 0 E = 150 H = 0 K = 4500 L = 3000 M = 254 O = 600 Q = human_chimp.v2.q T = 2 Y = 15,000”). Multiple alignments were generated from pairwise alignments with the MULTIZ (version 11.2) program (*89*), using default parameters and the following pre-determined phylogenetic tree: ((*A. calliptera*, *C. frontosa*), *O. niloticus*) in agreement with e.g. (*86*). The resulting reconstructed ancestral sequence is available at Dryad <added_prior_to_publication>.

#### Phylogenetic reconstruction

To reconstruct the phylogenetic relationships among all included 684 *benthic* samples we split the biallelic VCF files into windows of 100 kbp length, removing windows with less than 80% of sites passing all quality filters. We furthermore excluded the entire (extremely repetitive) chromosome 3 from the analysis, as well as the inversion regions on chromosomes 2, 9, 10, 11, 13 and 20. For the remaining 5,158 windows, we produced FASTA alignments with a modified version of vcf2phylip (*90*) and constructed trees with IQtree2 (*91*). We selected the least constrained substitution model (GTR+I+G) because the window size and substitution count were large enough (*92*). The trees were then summarised using ASTRAL III (*93*) (data S1). For the phylogeny shown in Fig. 1 we collapsed all monophyletic species to their average branch lengths. While all other taxonomic species formed monophyletic clusters, a single *Cyrtocara moori* individual fell within the otherwise monophyletic *Otopharynx argyrosoma*, rendering them polyphyletic, which we indicate with a ‘+’ at the *O. argyrosoma* tip label. We furthermore manually added non-*benthic* Malawi clades to the tree, based on a second phylogeny that was constructed analogously, but with fewer *benthic* samples (n=54) and representatives of each outgroup clade (7 *Diplotaxodon* sp., 2 *A. calliptera*, 24 *mbuna*, 5 *Rhamphochromis* sp.) (data. S2). All downstream tree operations including plotting Fig. 1 were done in python using the ETE toolkit (*94*).

#### Clustering-based inversion detection

To detect regions with suppressed recombination we identified clusters of windows which demonstrate phylogenetic patterns consistently distinct from the rest of the genome (code available at (*95*)). Pairwise distance matrices were computed in non-overlapping 100 kbp windows (as described in the methods section “Pairwise differences and divergence times”). Windows with less than 80% of positions passing all VCF filters were excluded. We flattened each distance matrix, converting it to a vector of length N of taxa squared and further used these vectors as data points. This means that pairwise distances between each pair of individuals were considered as variables and windows as observations. To reduce collinearity we applied PCA to this dataset and only retained the first 10 principal components. At this point we additionally filtered windows, excluding outliers (z-score > 3) on each of the ten PCs and then repeated the PCA. PCs for each window were normalised and clustered by k-means with N from 2 to 20 (fig. S2). To identify regions with long contiguous stretches of points falling into the same cluster, we calculated for each window the number of adjacent points within the same cluster. We considered only 10 points to the right and to the left of a focal point separately. Then we chose the greatest value of the two to avoid blurring of the abrupt transitions between spans of clusters (fig. S3). Windows having more than 8 of 10 neighbours from the same cluster (for N clusters = 8) on either side are marked with bars in the top track of Fig. 2A.

The rationale behind this approach is that the distance matrices for windows, which have the same phylogenetic history should ideally differ only by a local evolutionary rate and ancestral Ne variation which can be considered scaling coefficient and stochastic noise, PCA should extract scaling coefficient to one high weight component and noise on many low weight components. Major differences in the tree topology will introduce major changes in the distances which cannot be compensated just by changing the scaling coefficient, and these changes will be assigned to separate high weight components. Moreover, as the topology change produces discrete patterns of distances we expect to see discrete points on these topology-related PCs corresponding to individual topologies. Clustering then separates groups of windows sharing the same topology. Large stretches of the genome deviating in evolutionary rate or ploidy (e.g Y chromosome) will be probably also identified by the algorithm.

#### Windowed principal component analysis (PCA)

Genetic structure along the genome was analysed using a windowed PCA approach (*96*), which applies the pca function of scikit-allel (*97*) to the genotypes in a VCF file. Windowed principal components of 1 Mbp with a step size of 10 kbp were computed along chromosomes, applying a minor allele frequency threshold of 0.01. The resulting windowed PC1 values of the featured samples were plotted along the genome to identify regions with changes in genetic structure that are characteristic for inversions. To polarise PC scores along chromosomal windows we used the automatic guide sample selection approach implemented in the method. If this was insufficient, suited guide samples were manually determined and in some cases PC scores of individual windows were manually flipped. Initially, windowed PCAs were conducted for all seven clades and for combinations of these (fig. S4). After inversion identification and genotyping, two additional windowed PC analyses were conducted per inversion chromosome: (1) for all *benthic* samples homozygous for the inversion and (2) for all *benthic* samples homozygous for the non-inverted state. For visualisation in Fig. 2A, only a subset of 406 *benthic* individuals (species with ≥ 10 individuals sequenced) was included, and only every 5th window was plotted (50 kbp display step size), while the top 0.3% outlier windows for the % variance explained by PC1 are not shown.
We performed an additional windowed PCA for 27 *shallow rocky Aulonocara* individuals to investigate the possibility of an inversion polymorphism segregating among that group (fig. S16G, H).

#### Windowed heterozygosity

The number of heterozygous sites per sample was calculated in 1 Mbp windows along chromosomes (step size: 10 kbp) for a subset of 406 *benthic* individuals (species with ≥ 10 individuals sequenced) (*96*). For plotting (fig. S6), every 5th window was included, counts of heterozygous sites were divided by 100,000 and windows were normalised by the chromosome-wide average (for chromosomes with inversions, both the inversion region and the flanking regions were normalised by the average of the flanking regions).

#### Genome assembly

*Taxonomic coverage*
To evaluate structural rearrangements across the five major Lake Malawi cichlid clades *A. calliptera*, *mbuna*, *benthics*, *Diplotaxodon* and *Rhamphochromis*, we generated new Hi-C libraries and chromosome level genome assemblies for one representative species per clade: *A. calliptera*, *Tropheops* sp. 'mauve', *Aulonocara stuartgranti* (*deep benthic* subclade), *Diplotaxodon limnothrissa* and *Rhamphochromis* sp. 'Chilingali', respectively. In addition, we included contig-level long read genomes to represent the two

remaining benthic subclades: *Otopharynx argyrosoma* to represent *shallow benthics* and *Copadichromis chrysonotus* to represent *utaka*. All assemblies are based on read sets from different long read sequencing platforms: PacBio Continuous Long Reads (CLR), PacBio High Fidelity reads (HiFi) and Oxford Nanopore (ONT). In the following, we summarise the key characteristics of these seven assemblies, and how they were used to investigate structural rearrangements.

##### *Hi-C library generation*

Samples for *A. calliptera*, *T.* sp. 'mauve' and *Au. stuartgranti* were obtained from aquarium stocks of W. Salzburger in Basel, Switzerland. Per individual, ~100k sperm cells were used as input material for Hi-C libraries following the DovetailOmni-C library protocol. Libraries were sequenced at the Duke University Sequencing Center (Illumina Novaseq 6000 platform). Samples for *D. limnothrissa* and *R.* sp. 'Chilingali', were collected from aquarium stocks of D. Joyce (Hull, UK) and E. Santos (Cambridge, UK) respectively. Individuals were euthanized in a 0.1% MS222 solution and fin clips were immediately flash frozen and transferred to -80 °C for storage after collection. Library preparation (Hi-C - Arima v2 kit) and sequencing (Illumina NovaSeq 6000 platform) were performed at the Wellcome Sanger Institute (Hinxton, UK) according to the standard Darwin Tree of Life workflow (*98*). The raw sequencing data is accessioned at BioProject PRJNA1133007, diplim:<added_prior_to_publication> and ERR12668762.

##### *Long read sequencing*

We generated new ONT duplex data (3 PromethION R10 flow cells) for *Au. stuartgranti*: To reduce processing time on the GPU computing cluster, pod5 reads > 1 kbp were split into individual sequencing channels using Pod5 software (https://github.com/nanoporetech/pod5-file-format) (version: 0.2.2) and batched. Duplex basecalling was performed with dorado (https://github.com/nanoporetech/dorado) (version: 0.3.1) using model dna_r10.4.1_e8.2_400bps_sup@v4.2.0. The resulting duplex reads from 3 flow cells were then concatenated. We furthermore generated new HiFi data for the *D. limnothrissa* assembly using the standard Darwin Tree of Life sequencing workflow (*98*).

##### *Genome assembly and curation*

For *T.* sp. 'mauve' and *R. sp. 'Chilingali'* we started from our previously published PacBio CLR contig assemblies (*99*). Due to its insufficient structural integrity, we broke the *A. calliptera* reference genome (fastCal1.2) (*100*) into contigs at assembly gaps (stretches of exactly 100 'N's) for re-scaffolding. We assembled the newly generated ONT duplex and PacBio HiFi reads for *A. stuartgranti* and *D. limnothrissa* into contigs with hifiasm (*101*) (parameters: '--primary') and removed contigs representing haplotypic duplications with purge_dups (*102*) (default parameters).

For scaffolding, we mapped the Hi-C data to the corresponding contig assemblies using BWA-MEM2 (version 2.2.1, parameters: -5 -S -P -C -p) (*103*) and further processed with samtools (version 1.15.1, fixmate [parameters: -u -m -p], sort [parameters: -l 0], markdup [parameters:: -u -c]) (*79*) . Yahs (*104*) (default parameters) was then used to scaffold assemblies into chromosomes. For curation, we visualised Hi-C contact maps in JuiceBox (*105*) and generated pairwise whole genome alignments (see below). Taking into account Hi-C maps and pairwise alignments we adjusted the orientation of contigs if the Hi-C signal was inconclusive with respect to contig orientations. Especially when contigs are small or near chromosome ends, the scaffolding software may make mistakes about the orientations due to insufficient

Hi-C signal. In such cases, we used synteny information from pairwise alignments between five chromosome level assemblies to harmonise contig orientation, while prioritising assemblies of higher local structural integrity. This process resulted in five chromosome-level assemblies: fAstCal1.6 (*A. calliptera*, BioProject <added_prior_to_publication>), troMau2.1, (*T.* sp. 'mauve', Bioproject <added_prior_to_publication>) , fAulStu2.1 (*Au. stuartgranti*, BioProject <added_prior_to_publication>), fDipLim1.1 (*D. limnothrissa*, BioProject PRJEB77457) and fRhaChi1.1 (*R.* sp. 'Chilingali', BioProject <added_prior_to_publication>). We additionally included our previously published contig-level assemblies fOtoArg1.0 (*O. argyrosoma*) and fCopChr1.0 (*C. chrysonotus*). Assembly stats and accessions are provided in Table S4.

#### Comparative genomics analysis

We generated whole genome alignments between fAstCal1.6 and all other assemblies with minimap2 (parameters: -ax asm5 --MD) (*106*), and for the final chromosome-level assemblies re-aligned the Hi-C data exactly as described above. In addition, to represent outgroups, we aligned the publicly available assemblies for *Oreochromis niloticus* (Nile tilapia, GCF_001858045.2) and *Pundamilia nyererei* (representative of the Lake Victoria sister radiation) (*107*) to fAstCal1.6. We visualised pairwise alignments with a custom script (plot_bam_alignments.py) (*96*) and additionally used Hi-C maps (if available) to infer the presence/absence of the five focal inversions, as well as additional smaller or less frequent ones (see text S1).

#### Cichlid cell lines establishment and FISH

One captive bred male of *A. calliptera* and *Au. stuartgranti* were each euthanised, photographed, and dissected.

Primary cell cultures were established from caudal fin tissue, grown in DMEM/F12 media (Gibco, Life Technologies) supplemented with 10% FBS (Sigma- Aldrich), 1% antibiotic antimycotic solution (Sigma-Aldrich) and 1% non-essential amino acids solution (Gibco, Life Technologies), at 28 °C in an incubator. Metaphase chromosomes were harvested after incubation of cells with 50 ng/mL of nocodazole (Thermo-Fisher) for 2–3 h. Subsequently, cells were treated with a buffered hypotonic solution (0.4% KCl in 10 mM HEPES, pH 7.4) for 8–12 min at 37 °C and fixed with 4:1(v/v) methanol glacial fixative.

Fluorescent in situ hybridization probes were derived from long-range PCR products covering two sides of inferred inversion breakpoints, using GenomePlex™ Whole Genome Amplification Kits (Sigma-Aldrich) as described previously (*108*), and using CF532-dUTPs (Biotium), TexasRed-, atto-488-, Cy5-dUTPs (Jenna Bioscience) as the labels. Probes were prepared by mixing the labelled PCR products together with sonicated genomic DNA from cichlid fish and hybridization buffer. Freshly-prepared metaphase slides were immersed in acetone for 10 min and then baked at 62 °C for 1 hour. Slides were denatured in an alkaline denaturation solution (0.5 M NaOH, 1.5 M NaCl, Sigma-Aldrich) for 10 min. Probes were denatured at 65°C for 15 min and kept at 37 °C for 25 min. Hybridisation and post-hybridisation washes followed Louzada et al., 2017. FISH images were captured with a Zeiss Axio Imager D2 fluorescence microscope and images were processed using SmartCapture software (Digital Scientific UK).

#### Linked read analysis (haplotagging)

To investigate the structure of haplotypes we used the haplotagging molecular barcoding protocol (*109*), which attaches unique 24 nt tags to reads originating from the same DNA fragment of several kbp in size. After sequencing, tagged reads were mapped to the fAstCal1.2 reference using the ema aligner v0.6.2 ([https://github.com/arshajii/ema](https://github.com/arshajii/ema)). We computed the share of all tags in the window, which have at least one read with the same tag mapping to (a) the 1 Mbp region centred around the second inversion breakpoint ('second breakpoint'), (b) in the 100 kbp region next to the window under consideration (upstream and downstream separately: 'near left' and 'near right'), (c) more than 100 kbp apart from the window under consideration ('other distant'). For the chromosomes for which more than one sample with inversion was available we averaged these metrics over samples with the same inversion genotype. We applied rolling window smoothing to these values with a window of 5 data points for plotting.

#### Species cross

An interspecific cross was conducted between an *Au. stuartgranti* male from a lab strain originally established from a population sourced from Lake Malawi near the town of Usisya, and two *A. calliptera* females from a lab strain established from a population sourced from the Itupi river in Tanzania. Nine F1 individuals were reared from two different broods, with different mothers. These F1s were intercrossed to generate 129 F2 individuals, which were in turn intercrossed to generate 165 F3s. Sex was determined by coloration and only male F2s and F3s were sequenced. All individuals to be sequenced were kept in isolation upon reaching sexual maturity. Specimens were euthanized in a 0.1% MS222 solution and fin clips were collected and stored in 100% ethanol at -20 °C. DNA extraction and library preparation followed the same protocols as for the wild samples above. Libraries were sequenced (150 bp paired end reads) on a NovaSeq 6000 S4 or or NovaSeq X 10B machine to a mean coverage 4.5x to 8.6x (F1 - F3 samples) and 17.4x and 21.1x (*A. calliptera* and *Au. stuartgranti* G0 samples) (for per sample-information refer to table S5). The raw sequences are available at BioProject <added_prior_to_publication>. Sequencing reads were mapped to the fAulStu2.1 assembly using bwa-mem (*78*) (for code see (*110*)) and biallelic variants were called with bcftools (*79*) and quality-filtered (for code see (*111*)). Windowed principal components were computed and plotted exactly as described above for the full dataset of wild samples.

#### Centromere localization in the reference genome

Because centromeres were mostly unassembled in the PacBio CLR fAstCal1.2 assembly, we instead focused on the more recent *D. limnothrissa* HiFi assembly, which is based on PacBio HiFi technology. First, we used TRASH (version: 1.2) (*112*) with default settings to identify high order repeats (HOR). We found multiple HORs with 237 ±10 bp unit size across different chromosomes, consistent with centromeric repeats. To obtain a genome-wide centromeric repeat consensus we filtered the TRASH output for high quality HORs (score >= 80 and size 237 ±10 bp, ignoring unplaced scaffolds), resulting in 42 HORs. We concatenated the corresponding 42 repeat unit consensus sequences, rotated them with rotate (*113*) version: 1.0, parameters: CATGTTTATGGCTTTATCT -m 2 -rc, one dropout) and aligned them with MAFFT (v7.309, default parameters) (*114*), resulting in a 237 bp genome-wide centromeric repeat consensus sequence after removing alignment columns with <50% missingness:

TTTGTCTGAGTCTTCTATCAAAAGTTACAGCTATTTATATGAAGTTGTACAAAAACGCTTTCTTT CGCCAAGACAGTGCGTTTCTCACTGTTACATGCATTTGAATGGGGAACTCGACCGAAACAAA GTATAGGTTTTCATTATGTGAATAACTTGAAAATCTTAGCTCAAACAGCTGCAAAACCTGTTTT CCCAGCACGGGGACCTTGAGTTCTATAAGATAAAGCCATAAACATG.

To identify centromeres in the fAstCal1.2 assembly, we used BLAT (v38x) (*115*) to search for alignments of the 237 bp repeat consensus sequence. In addition, we extracted the flanking regions around centromeric HORs in the *D. limnothrissa* assembly (TRASH score ≥ 50, 237 ±10 bp unit size) and aligned them to the fAstCal1.2 assembly with minimap2 (v2.28-r1209, parameters -a, sorted with samtools sort). Taking all available evidence into account (BLAT hits ≥ 100 bp of the centromeric repeat unit, alignments of the centromere-flanking regions from the *D. limnothrissa* assembly, assembly gaps), we then determined the likely locations of centromeres in the fAstCal1.2 assembly with different levels of certainty (fig. S17, table S6).

#### Structural variants and transposable element scan

To compare the transposable element (TE) composition of corresponding inverted and non-inverted haplotypes we assessed structural variants between the genomes of two species with opposite inversion states for the chromosome 9 and 11 inversions. For this purpose, we aligned our *Au. stuartgranti* (fAulStu2.1) assembly to the (re-scaffolded) *A. calliptera* (fAstCal1.6) reference with minimap2 (version 2.24, parameters: -ax asm5, --eqx) (*106*) and sorted the output with samtools (version 1.14) (*79*). The resulting BAM file was then used as an input to call structural variants with SVIM-asm (*116*) in haploid mode. The SVIM-asm output comprises insertions, deletions and duplications in the query (*Au. stuartgranti*) compared to the reference (*A. calliptera*).

We extracted insertions and deletions ≥50 bp and annotated them with Repeatmasker (version 4.0.7) (*117*) using a combined cichlid TE library from repbase (*118*) and additionally a de novo annotated TE libraries for fAstCal1.2*,* which was produced with RepeatModeler2 (*119*) and EarlGrey (*120*). Since assembly quality varied considerably between the two genomes (table S4) we limited the downstream analysis to fAulStu2.1 regions with alignments to fAstCal1.6 (considering only minimap2 primary alignments with mapping quality ≥30), to restrict the results to regions assembled in both genomes. For those regions, we calculated mean TE lengths for inserted and deleted sequences, both genome wide (ignoring inversion chromosomes) and for inversion regions. We split the results at a 0.5% divergence threshold (bin 1: divergence < 0.5%, bin 2: divergence > 0.5%), which reflects between-species-divergence of *A. calliptera* and *Au. stuartgranti* – following the rationale that within the lower divergence bin (< 0.5%) we expect to capture predominantly recent TE diversity after the species split, while within the higher divergence bin (> 0.5%) we expect predominantly to capture shared TE diversity that built up before they separated. To compare TE content within and outside of the inversion regions we furthermore calculated Z-scores based on the genome wide mean (using only chromosomes without inversions) for the inverted chromosomes 9 and 11 of *Au. stuartgranti* in windows of 1 Mbp with step size of 100 kbp.

To look for potential translocations of sex determining genes in the inversion regions of chromosome 9 and 11 of *Au. stuartgranti*, we performed tblastn (BLAST+ suite version 2.14.1) (*121*) of all the insertion and deletion sequences ≥ 50 bp against a list of 124 known sex determining genes (*61*, *76*, *122*), with an E-value cut-off of 1e-5. When we filtered the results for best hits per query we found that the genes that were coded by these sequences were not present in the insertion and deletion sequences in the inversion chromosomes of *Au. stuartgranti* (table S29).

#### Breakpoint estimation

Approximate locations of the boundaries of inversions ("breakpoints") on the reference genome (fAstCal1.2) were determined by conducting windowed PC analyses at higher resolution (step size: 5 kbp, window sizes 1 Mbp, 100 kbp, 50 kbp), using python functions implemented in ref (*96*). Based on this, approximate breakpoint regions were determined around the onset (left breakpoint) and cessation (right breakpoint) of the inversion-characteristic stratification of samples into inversion genotypes, with final approximated regions spanning 50 to 80 kbp each. We used two sets of start and end positions throughout to delineate inversions throughout analyses: In the majority of analyses, inversions were defined as the region between the midpoints of the breakpoint regions. Some analyses, which required us to be more conservative, use only the "inner" region between the right boundary of the left breakpoint and the left boundary of the right breakpoint, which is indicated in the corresponding methods sections. The genomic positions of the breakpoint regions and the midpoints are reported in table S8.

#### Inversion genotyping of whole genome-sequenced samples

To genotype all 1,375 sequenced individuals and to infer which homozygous state is ancestral we designed a multi-step PCA approach using biallelic variants in each inversion region (fig. S21). All PCAs were performed using scikit-allel (*97*) on biallelic variants with no missing genotype calls. First we used an initial PCA of of all *benthic* samples ('*PCA I*') to select two sets of individuals representing the two homozygous inversion genotypes (these groups are equivalent to the upper and lower PC1 bands of inversions in the windowed PC analyses shown in Fig. 2A). To be precise, we selected all samples from the band with fewer individuals and a random subset of the same size to represent the other homozygous group. With these two sets of samples, we conducted a second PCA of each inversion region ('*PCA II*', not shown in fig. S21). Then, using only the subset of sites that were variable among the two homozygous sample sets in *PCA II*, but including all sequenced individuals (i.e. also non-*benthics*), we performed a third PCA ('*PCA III*'). We used *PCA III*, which emphasises inversion-related over phylogenetic variation to assign inversion genotypes (homozygous vs. heterozygous) to all 1,375 samples based on the inversion-related stratification on PC1. We repeated the entire procedure multiple times to confirm that sample selection from *PCA I* as input to the *PCA II* step did not affect the final genotyping outcome in *PCA III*, and always obtained identical inversion results. We then assigned the ancestral orientation to the genotype cluster containing non-*benthic* clades (except *Diplotaxodon*, see below) in *PCA III*. For confirmation of ancestral state assignments we examined whole genome alignments between *A. calliptera* and two outgroup species for ancestral state assignment (see text S1, fig. S22 and fig. S21). We furthermore conducted a PCA equivalent to *PCA I* but for the flanking regions of the same chromosome instead of the inversion region. For chromosome 9, where only the homozygous inverted genotype was present among *benthics*, the same steps were performed except that the heterozygous genotype cluster was used in the *PCA II* step in place of the missing homozygous cluster.
Notably, for the inversions on chromosomes 9 and 11, all *Diplotaxodon* samples localized closer to the inverted genotypes on PC1 than all other outgroups. We thus used our new chromosome-scale genome

assembly for *Diplotaxodon* (see below) and found both inversions. Accordingly, we assigned homozygous inverted genotypes to all *Diplotaxodon* individuals for the chromosome 9 and 11 inversions. For chromosome 20, which contains a differentially fixed inversion between *Diplotaxodon* and all *benthics* we conducted the *PCA I* step and visualized PC1 to PC4 to support the absence of an inversion polymorphism segregating among *benthic* clades (fig. S16A-F).

#### PCR assay to genotype inversions

To design the assay, we identified polymorphic TE insertions that were correlated with inversion state. First, to identify a set of candidate TE insertion sites, we visually examined BAM files for a small set of samples (four individuals for each homozygous state) in IGV (v2.11.4) (*123*), considering only the proximal 1 Mbp regions adjacent to the left and right breakpoint of each inversion. We then queried candidate TE polymorphisms across BAM files of all sequenced samples, counting clipped and spanning reads. Per inversion, we selected the most correlated TE insertion polymorphism. Among 1,375 samples we found only 4 (chromosome 9, but see below for additional details on this specific assay), 5 (chromosome 11), 5 (chromosome 10), 2 (chromosome 13) and 2 (chromosome 2) cases of mismatch between TE and inversion genotype (> 99% concordance).

Therefore, we went ahead and developed diagnostic sets of three primers per inversion to assay the presence or absence of each diagnostic TE. Two complementary primers were designed to flank the TE insertion site and would amplify in the absence of the TE. Additionally, a third primer complementary to one of flanking primers was designed to be located within the TE, so that it would amplify in the presence of the TE. For that matter, we assembled the relevant TE sequences with MEGAHIT (v1.2.9) (*124*) using the paired-end reads that mapped ± 1000 bp of the candidate site in at least 30 individuals. Due to the size difference between the alternative amplicons we were able to infer inversion genotypes through multiplex PCA. Per assay, amplification of only the longer or only the shorter fragment was diagnostic of homozygous presence or absence of the inversion while amplification of both fragments indicated inversion presence in heterozygous state. The primer sequences and a schematic of the primer design are provided in table S9 and fig. S24.

Importantly, for the chromosome 9 inversion, we selected a TE insertion specific to non-inverted *benthic* haplotype. Since this haplotype only occurs in heterozygous state among all WGS-typed samples, we classified *benthic* individuals without the TE insertion as homozygous for the inversion and non-*benthic* individuals without the TE insertion as homozygous for the ancestral Malawi state.

We used this assay to estimate the association between sex and inversion states in 401 additional Malawi cichlid samples belonging to 57 *benthic* species. For inversions on chromosomes 11 and 13, the assay yielded clear genotyping results for all samples. For the remaining three inversions (chromosome 2, 9 and 10) some PCR assays yielded ambiguous results. In these cases we repeated the PCR once and only considered results if unambiguous bands were visible in the second iteration. This allowed us to type inversion states of chromosome 2 inversion in 387 samples, states of chromosome 9 inversion in 359 samples and states of chromosome 10 inversion in 396 samples (tables S10 and S11).

#### Test for structured Hardy-Weinberg equilibrium

Given the generally low sample count per species (median 2, mean 5.7), our dataset lacks power to test for Hardy-Weinberg equilibrium (HWE) of (inversion) genotype frequencies within species. We therefore implemented a test for HWE equilibrium under population structure proposed by Hao and Storey (*125*), by using per-species sample allele frequencies as estimates of species allele frequencies, and resampling genotype frequencies following Hao and Storey's Algorithm 1 with 10,000 replicates. We consider this procedure conservative for detecting heterozygote excess, because non-accounted population structure within species (we effectively assume random mating) will bias tests towards an excess of homozygotes. Consistent with this, a significant excess of homozygotes was observed for the two inversions without evidence for sex linkage (p-value = 0.029 and 0.0016 for chromosomes 2 and 13, respectively).

#### Sex association

The biological sex of specimens was assessed either through visual examination of fresh specimens, based on breeding colouration, presence of mouth-brooding in females, or through dissection and examination of gonads. This led to the assignment of whole genome sequenced samples to the states: male (M, 669 specimens), female (F, 135), potential female (F?, 56), potential male (M?, 9), unclear (U, 506). We note that the distinction of females from subdominant males is challenging in visual sex-assessment, which is reflected in the large number of unclear assignments. Sex was assessed visually for the majority of wild-caught samples with the notable exception of *C. chrysonotus* from Lake Malombe which show perfect sex association (see main text), and for which gonads were examined. See table S1 for per-specimen info. For the 141 samples used in PCR-based inversion typing (tables S10 and S11) sex was assessed through gonad inspection.

To test for association between inversion genotype and sex, we merged PCR-genotyped and whole genome-sequenced samples and discarded samples with unambiguous sex determination (table S28). Considering only species with at least one heterozygous individual, we conducted Fisher's exact tests for each combination of inversion-genotype and sex using the scipy.stats package (version 1.10.0) (*126*). Associations reported in the main text remained significant when WGS and PCR-typed samples were considered independently (except for chromosome 10, which was not significant for WGS samples alone).

Windowed PCA and inversion-wide heterozygosity (Fig. 5A, B) were re-calculated separately for a subset of 28 *C. chrysonotus* from Lake Malombe as described for the full dataset, but using a window size of 1.5 Mbp in the PC analysis.

#### Lake Victoria radiation inversion scan

We fetched 104 short read sequencing datasets from SRA (PRJNA626405) with broad phylogenetic coverage of the closely related Lake Victoria cichlid radiation (*59*). We downloaded the read sets from NCBI using sratools (https://github.com/ncbi/sra-tools) (prefetch with default settings, fastq-dump [parameters: --gzip --skip-technical --read-filter pass --dumpbase --split-e]) and mapped them to the Lake Victoria *Pundmilia* reference genome using bwa (index and mem, default settings). We further processed BAM files with samtools (sort [parameters: -n -l 0], fixmate [parameters: -u -m], sort [parameters: -l 0], markdup [parameters: markdup [parameters: -u -S -T])(*79*), converted them to CRAM, adding NCBI

RunID to the read group SM tag (samtools addreplacerg [parameters: -w -r ID:${run_id} -r SM:${run_id} -O CRAM --reference]) and indexed the BAM files (samtools index, default parameters). We called and quality-filtered biallelic SNP variants (*111*) and computed and visualised windowed principal components exactly as described above for the full dataset of wild samples, using metadata available in the paper (taxonomy) and from the bio sample information on NCBI (sex assignment). We furthermore replotted LOD scores from published interspecific QTL crosses (*58*).

#### Inversion haplotype rephasing

Despite statistical phasing (see above), both haplotypes in inversion heterozygotes did initially show intermediate phylogenetic clustering consistent with frequent phasing errors. We therefore rephased heterozygous sites in inversion heterozygous individuals by calculating for each SNP Pearson's correlation coefficient between inversion and SNP derived allele count across all benthic samples and assigning the allele showing a positive Pearson's correlation to haplotype 2 and the other allele to haplotype 1 (see section SNP–inversion-correlation below for more details). SNPs with $|r| < 0.05$ were assigned to haplotypes randomly. To confirm that violations of the normality assumption of Pearson's r did not have a major impact on phasing, we also calculated Kendall's $\tau$, which showed the same sign as Person's r (and would thus have led to the same phasing) for > 99.99% of rephased SNPs.

#### Haplotype-resolved local phylogenetic inference in inversion region

For each inversion region we computed separate phylogenetic tree, following the same steps as for the 684 *benthic* samples tree (see *Phylogenetic reconstruction*), but using the rephased SNP data for the inversion regions (see *Inversion haplotype rephasing*) and treating the (phased) haplotypes of the same sample as separate OTUs in the tree. To reduce computational burden we selected no more than 10 samples for *benthic* and *Diplotaxodon* species and added one sample per species for *Astatotilapia*, *mbuna*, and *Rhamphochromis,* resulting in 488 OTUs in total. The counts of local inversion region trees were as follows: 228 on chromosome 2, 170 on chromosome 9, 181 on chromosome 10, 221 on chromosome 11 and 200 on chromosome 13.
To build co-phylogenetic trees we pruned the haplotype-resolved trees and the corresponding genome-wide phylogeny to the same set of samples using the ETE toolkit package in python. A co-phylogeny was then generated per inversion with the genome-wide tree on the left and the inversion region haplotype-resolved phylogeny on the right using the cophylo() function from the phytools R library (*127*), but rotating only the right tree. Connector lines were colour-coded by *benthic* sub-clade.

#### Pairwise differences and divergence times

To estimate pairwise divergence (coalescent) times in inversion regions and in the rest of the genome, we first calculated pairwise SNP differences between both haplotypes of all samples using custom python scripts in python 3.9.7, specifically the function get_pwd_and_trees in the python3 module diversity, which is available at (*128*). This function is based on scikit-allel's pairwise_distance function with option metric='cityblock', but corrects for missing genotype calls by assuming they correspond to the inferred ancestral allele.

The function was applied to rephased biallelic SNP VCFs separately for each of the inverted regions on chromosomes 2, 9, 10, 11, 13 and to the non-inverted regions (i.e., all other chromosomes and all regions outside of the inversion breakpoints on chromosomes 2, 9, 10, 11, 13). The latter were summed to obtain total "non-inverted" pairwise differences for all combinations of haplotypes. Pairwise differences were normalised to obtain mean absolute divergence ($d_{xy}$, for haplotypes of distinct individuals) and nucleotide diversity ($\pi$, haplotypes within individuals) by dividing total difference counts by the accessible genome size of the respective genomic region (non-inverted or inverted). These values were further translated into divergence times ($T_{coal}$) using our previous mutation rate estimate of $\mu=3*10^{-9}$ /(bp*generation) (*31*) according to the relation $d_{xy}=2*T_{coal}*\mu$.

The resulting estimates are displayed in Fig. 3A, where pairwise comparisons are grouped by inversion state for *benthics* and only inversion-homozygous samples are considered. An exception to this is chromosome 9 for which the homozygous non-inverted state is absent among *benthics*, and therefore the rephased haplotypes of heterozygotes were used in comparisons. Fig. 3A is based on the matplotlib hist function (*129*) with option density=True and 100 bins between 220,000 and 380,000. Haplotype divergence times across all *benthics* and the difference of estimates for inversion region compared to the remaining genome are shown in fig. S36.

#### Investigation of chromosome 9 haplotype origin

To investigate the origin of the non-inverted chromosome 9 haplotype present in *benthics*, we created a second SNP callset containing representatives of all major Malawi clades, as well as wide selection of other African haplochromine riverine and lacustrine cichlid species. We selected 11 species from the Lake Tanganyika radiation, 29 from the Lake Victoria Region Superflock (LVRS), and 32 species from other lake and riverine lineages within the Zambezi, East Central Coast, Congo and Nile Basins, giving a total of 112 non-Malawi African haplochromine samples. For the Malawi radiation, we selected 500 samples from 239 species, representing each of the Malawi ecomorphological clades (*Diplotaxodon*, *Rhamphochromis*, *shallow benthics*, *deep benthics*, *utaka*, *mbuna* and *A. calliptera*) and, with the exception of *A. calliptera* (number of samples = 113), included a maximum of two samples per species. For non-Malawi samples, we acquired Illumina short-read sequence data from the Sequence Read Archive (from studies ERP107954, SRP065582, SRP086337, SRP094973, SRP098665, SRP116725, SRP148476, SRP156808, SRP213801, SRP215535, SRP276092, SRP285149, and <added_prior_to_publication>).

Sequences were aligned to the *A. calliptera* (fAstCal1.2) reference genome (NCBI GenBank: GCA_900246225.3) (*100*) using using BWA-MEM (version 0.7.17-r1188) (*78*) and genomic variants were called using bcftools (version 1.14) (*79*). We removed variants within 10 bp distance to another variant, sites with an overall mapping quality (MQ) below 50, and sites where the forward and reverse strands had a significantly different MQ ($p<0.001$ in a Mann-Whitney U test). We also filtered out sites where >10% of mapped reads have an MQ = 0, sites with an inbreeding coefficient <0.2, sites with >20% missing genotypes, sites with abnormally high sequencing depth across all samples (>1.5 standard deviation above the mean), and heterozygous sites exhibiting significantly biased read depth between the reference and alternative alleles (PHRED score >20 in a binomial test). Finally, we filtered only for biallelic SNPs, and added to the final VCF callset the ancestral state sequence as an additional sample (see ancestral state inference methods).

To infer the phylogenomic relationships between Malawi and the other African haplochromine cichlids in our callset, we constructed a maximum likelihood (ML) tree using the GTR+G+I model in IQ-TREE (version 2.2.0-beta) (*130*), with the ancestral state sample used as an outgroup. We created trees in 100 kbp non-overlapping windows across the entire genome, which were then used to build a consensus tree using ASTRAL-III version 5.15.5 (*93*). The final consensus ML tree was visualised using FigTree version 1.4.4 (http://tree.bio.ed.ac.uk/software/figtree/), and rooted to the ancestral state sample.

The non-inverted benthic chromosome 9 haplotype is only present in heterozygous state. To amplify the signal pertaining to this haplotype, we calculated a SNP–inversion correlation (see *SNP–inversion-correlation*), but treating both "native" Malawi haplotypes (inverted and non-inverted) as the same state and calculating correlation of SNP genotypes with chromosome 9 inversion heterozygous state across all Malawi clades. We used this measure to identify for each SNP the "foreign" (introgressed) allele in chromosome 9 inversion heterozygotes as the one rare among other Malawi cichlid haplotypes and produced two separate genotype calls for each heterozygous individual, one homozygous for foreign alleles (chr9het_foreign), one for Malawi alleles (chr9het_malawi), respectively. We then used Dsuite Dtrios (*131*) to compute ABBA-BABA tests across the chromosome 9 region, treating all chr9het_foreign and all chr9het_malawi as well as each non-Malawi species as a separate population. We then focused on the 76 tests for which P3 was a non-Malawi species and P1 and P2 were among chr9het_foreign and chr9het_malawi. For 46 of these trios, chr9het_malawi was in the P2 position (i.e., closer to the P3 outgroup), whilst for 30 trios chr9het_foreign was in the P2 position and thus closer to the P3 Malawi species. When testing for significance we used the Bonferroni method to control for the family-wise error rate.

#### Inversion-wide heterozygosity

Bcftools stats (*79*) was executed on each inversion region independently and additionally on the remainder of the genome for the same subset of 406 benthic individuals used in Fig. 2A (species with at least ten sequenced samples). Per sample counts (PSC) were extracted from the output and the number of heterozygous sites ("nHets") per sample was normalised by the proportion of accessible SNPs (see *SNP detection*) of the inversions or of the rest of the genome, respectively. For each inversion, samples were binned by inversion state and clade adherence, and the genome-wide average heterozygosity value per sample was subtracted.

#### Probability of shared polymorphism

To better understand whether the sharing of inversion polymorphism between deep and shallow benthics for the chromosome 2, 10 and 13 inversions is likely to be the results of incomplete lineage sorting (ILS; i.e., in the absence of gene flow), we followed an approach used by Knief et al. (*28*), who used coalescent first principles to derive a formula for the sharing of polymorphism between two reproductively isolated species under neutrality. Under some simplifying assumptions (see ref (*28*)), this probability can be approximated by $p_{ILS} = \frac{2e^{-2\tau}}{11+12\tau}$, where τ is the species split time. We estimated τ for deep and shallow benthic divergence as $\tau = \frac{mean(d_{XY}) - median(\pi)}{median(\pi)} = 0.38$, where $d_{XY}$ is pairwise sequence divergence between *deep* and *shallow benthic* samples and π is nucleotide diversity (heterozygosity) within *benthic* samples. This yields a sharing probability of 6% for an individual inversion. Given that the three inversions segregate on independent chromosomes, the probability of sharing polymorphism at all three of them is $0.06^3 = 0.0002$. We also note that this estimate is conservative in the sense that this probably would even be lower if selection had acted on the inversion haplotypes, for which we present evidence in Fig. 4.

#### Gene flow analysis

To test whether inversion sharing is linked to gene flow signals in the rest of the genome, we grouped all benthic samples by species and compiled a Sets.txt as described in the manual of Dsuite (*131*) assigning as outgroup the ancestral sequence inferred above (which we had added as a sample to the VCF file). We then used Dsuite Dtrios to compute ABBA-BABA tests (D statistics, f4 ratio) across all chromosomes except those with large inversions (2, 9, 10, 11, 13), using 20 block-jackknifing windows per chromosome (-j 20). For further analysis we used a conservative significance threshold of block-jackknifing z>5.5 corresponding to a Bonferroni FWER<0.01. The _BBAA.txt file resulting from Dsuite Dtrios orders trios in a way that BBAA>ABAB, BBAA>BABA and ABBA>BABA (i.e., P1 and P2 share more derived allele with each other than with P3 and P2 shares more derived alleles with P3 than P1 shares with P3). Loading the file into python3, we selected for each inversion chromosome all trios for which (i) P1 and P2 were from the same subclade (*deep* or *shallow benthic*) and P3 was from the other subclade, and (ii) the inverted haplotype was present (i.e., had frequency > 0) in one of the P1 and P2 species but absent (frequency=0) in the other. We then grouped the resulting trios into two groups: (1) those for which the inversion state of P3 (presence/absence) was shared with P2; and (2) those for which the inversion state of P3 was shared with P1. For the former, f4 ratios are displayed on the positive x-axis in the histograms of Fig. 3C, while for the latter, f4 ratios are displayed on the negative x-axis in Fig. 3C (i.e., were multiplied with -1.0). Under the null hypothesis of inversion sharing across *deep* and *shallow benthic* clades being due to ILS (and thus independent of gene flow), we expect the resulting distribution of f4 ratios to be symmetric around zero. Conversely, if inversions were transferred by genetic introgression we expect a positive skew of these ABBA-BABA statistics towards positive values. Chromosome 9 was excluded from the analysis because of the very low number of non-inverted alleles among *deep benthics*.

#### Principal component analysis of *benthic* samples

To obtain further information on genetic relationships among benthics, we subsetted the VCF files for each chromosome to only include *benthic* samples and only SNPs outside of the inversion regions using bcftools view --keep <samples> --regions <outside-inversion-breakpoints>, then concatenated all VCFs using bcftools concat (*79*), and translated them into plink binary files using plink v1.90b6.21 (*132*) with the option --make-bed. We further used plink to prune SNPs in high LD using the option --indep-pairwise 500 10 0.1 and then --extract. On the resulting file we performed PC analysis with a minor allele frequency cutoff of 1% by running plink with the options --pca --maf 0.1 --chr-set 23 no-xy no-mt. The resulting first two PCs are shown in fig. S37.

#### Habitat depth and inversion state

Depth data for the benthic species included in this study was obtained, when available, from our sampling data collected during a trip to Lake Malawi between October and December 2023. Information on catch depth was obtained from fishermen (for artisanal fisheries catches) and from a Malawi Fisheries Department trawl survey on board the research vessel R.V. Ndunduma. During the trawl survey, a total of 116 stations were sampled by bottom trawling over 30 minute sessions. We first approximated the mean depth of each session as the mean value between the depth of the net at the start and the end of the session. Then, we estimated the mean depth per species as the average depth across all sessions, including the data from the trawl survey and from fishermen. This data is summarised in table S14. We intersected species with available depth data and those that were sequenced and featured in our 684 sample *benthic* phylogeny. We pruned the phylogeny to one representative per species (n=75) and converted the resulting tree to ultrametric. Finally, we plotted inversion frequency (using the joint PCA and PCR inversion genotyping; table S2) along with the mean depth per species. All tree operations including plotting were performed in python using the ETE toolkit (*94*).

#### Fixation index ($F_{ST}$) population branch statistic (PBS) in windows

$F_{ST}$ and the population branch statistic PBS, a variation of the commonly used $F_{ST}$ statistic that uses three populations and can detect evolutionary divergence specific to one population (*133*) was computed using ANGSD (v0.941) (*134*) for the same subset of 406 benthic individuals used in Fig. 2A. First, site allele frequency (SAF) likelihoods were precomputed separately for the three benthic subclades *deep benthic*, *shallow benthic* and *utaka* from phred-scaled genotype likelihoods (angsd -doSaf 1 -vcf-PL $vcf -nLines 500 -P 3 -anc $anc_fasta), supplying the ancestral state reference sequence in FASTA format. Next, pairwise 2D site frequency spectra (SFS) were calculated and indexed (realSFS, default settings). Finally, $F_{ST}$ between all three *benthic* groups and PBS were calculated and summarised in overlapping 1 Mbp windows (realSFS fst stats2 -win 1000000 -step 10000) and plotted along chromosomes.

#### MSMC analysis

We used MSMC2 (version 2.1.1) (*135*) to assess differences in the demographic patterns between the inverted and the non-inverted haplotypes. For the comparative analysis we selected *deep benthic* species fixed for the inversions on chromosomes 2, 10, 11 and 13 and *shallow benthic* species featuring no inversions on the same chromosomes. Chromosome 9 was excluded from the analysis as it lacked the homozygous non-inverted state in *benthics*. For each species we selected four samples from the same sampling location with at least 12 X sequencing coverage to create the MSMC2 input files (.multihetsep). We calculated the coalescence rate for the entire non-inverted region for each species using the .multihetsep files from all non-inverted chromosomes together (i.e, all chromosomes except 2, 9, 10, 11 and 13). To calculate the coalescence rate for the inverted regions we used the .multihetsep files from each inversion chromosome separately for both species with and without inversions. For all estimations we specified the time interval pattern (-p) to be 1*2+15*1+1*2 along with the '-s' option to skip over ambiguously phased sites. Next the combine script (combineCrossCoal.py) from the MSMC toolkit was used to calculate cross-coalescence between pairs of each *deep benthic* species with inversions and *shallow benthic* species without the inversions. Additionally, we used MSMC-IM (*136*) on the MSMC cross-coalescence estimations with default parameters and mutation rate '-mu' set to $3.5\times10^{-9}$ based on

(*31*). MSMC-IM incorporates an immigration isolation model for a more detailed estimation which helps to understand certain trends seen in the estimates of the original MSMC results.

#### Genome-wide association analysis

To investigate the extent to which the chromosomal inversions on chromosomes 2, 9, 10 and 13 contribute to genetic divergence between *shallow* and *deep benthic* clades, we conducted a genome-wide association (GWA) analysis for genetic clade adherence using GEMMA v0.98.3 (*137*). The analysis was performed by fitting a univariate linear mixed model including 543 benthic samples ($N_{shallow\ benthic} = 389$, $N_{deep\ benthic} = 154$), specifying the phenotype (genetic clade) as a binary trait. Sites with minor allele frequency (MAF) < 5% were excluded (default), after which 2,385,167 SNPs were analysed. A centered relatedness matrix (option '-gk 1') was used to account for population structure.

To visually compare the estimated effect sizes contributed by inverted and "non-inverted" regions of the genome (all chromosomes without inversions and inversion chromosomes excluding inversion regions), we calculated, per chromosome, the difference between the observed SNP effect sizes (sum of effect sizes divided by total effect size genome-wide) and the null expectation (chromosome size relative to genome size) (fig. S40).

#### SNP–inversion-correlation

For each combination of SNP and inversion region, we computed Pearson's correlation coefficients between SNP and inversion genotypes across all *benthic* samples using the python function scipy.stats.pearsonr. Both the correlation coefficient, r, and the score (-log10(p-value)) were used in further analyses. A negative sign was recorded for both r-values and scores if the derived allele was at higher frequency on the non-inverted haplotype. We note that p-values below a (r-dependent) threshold of $\sim 10^{-230}$ were recorded as 0.0 by python. The corresponding scores were thus set to the score largest in absolute value for that chromosome. SNPs with high absolute r values and scores are referred to as "inversion correlated SNPs" (ICS) in the text. The following ICS thresholds were used in different analyses: (1) For GO enrichment analysis, ICS with $|r| > 0.99$ and $p < 10^{-230}$ were identified and genes with transcription start sites within ± 25 kbp of these ICS were considered; (2) "Non-synonymous ICS" (nsICS) were considered for zebrafish expression analysis (see below) and literature search if they had a $p<10^{-200}$, which corresponded to $|r|>0.86$. Finally, we computed average SNP-inversion correlation scores in windows of 100 adjacent SNPs with 25 SNPs overlap (Fig. 4D) and considered the positive and negative 0.5% outlier windows in further analysis. We note that Pearson's r coefficients showed extremely high correlation with Kendall's τ (Spearman's r = 0.98, $p<10^{-16}$), which is robust to violations of normality.

#### Simulations of selection on inversion haplotypes

We used SLiM 4.3 (*138*) to perform forward-in-time simulations of the evolution of inversion haplotypes. Rather than exploring the full large parameter space of possible evolutionary models, we focused on plausible scenarios, investigating the effect of the strength of selection and of the number of selected sites on inversion maintenance and the dN/dS ratio. Full code and parameter combinations are available at (*139*). In short, we simulated the evolution of chromosomes in a scenario of two habitats (called p1 and p2) under the Wright-Fisher model with symmetric migration. The carrying capacity (population size) of habitat p2 was set to 5000 individuals, while the one for p2 was initially assumed to be between 1% and 20% of that of habitat p1, representing an ecological niche that is not yet fully explored.

Chromosomes were modeled such that there are coding regions separated by intergenic sites. Coding regions were composed of synonymous and non-synonymous sites. Mutations at intergenic and synonymous sites were always neutral, while mutations at non-synonymous sites were of two types: (1) Mutations which had an unconditionally deleterious fitness effect and (2) mutations which had a conditional fitness effect, being deleterious in p1 and adaptive in p2. The fitness of individual $i$ in population p was calculated as

$$f(i,\, p) \; = \; \frac{(1+S_D^p)^{Y(i)} \cdot (1+S_C^p)^{X(i)} \cdot (1+\epsilon^p)^{X(i)(X(i)-1)/2}}{\sum_{j=1}^{N_p} (1+S_D^p)^{Y(j)} \cdot (1+S_C^p)^{X(j)} \cdot (1+\epsilon^p)^{X(j)(X(j)-1)/2}},$$

where $X(i)$ and $Y(i)$ denote the number of copies of conditionally adaptive and deleterious mutations, respectively, and $S_C^p$ and $S_D^p$ their respective fitness effects. $N_p$ denotes the population size. The interaction (epistasis) between conditionally adaptive mutations was modeled with the term $\epsilon^p$.

After a burn-in of $4N$ generations during which deleterious mutations (genetic load) are expected to reach an equilibrium in both populations, we sampled a random haplotype from p2, added $X$ conditionally adaptive mutations and introduced an inversion that partially suppresses recombination with non-inverted haplotypes. We chose not to model the actual random accumulation of X conditionally adaptive mutation on a given haplotype and the probability of this haplotype being subject to an inversion explicitly, because the waiting time for these events to happen would be long and the goal of the simulation was to study the effect of different evolutionary regimes on inversion maintenance and dN/dS ratio, rather than the establishment of inversions in the first place. Once the inversion was introduced, the population size of p2 grew exponentially to match the population size of p1, representing an expanding ecological niche. After a sufficient number of generations to allow the inverted haplotype to become fixed or nearly fixed, we ended the simulation and estimated the dN/dS ratio as described in *Selection tests (NI, DoS, dN/dS)*.

We simulated data for two scenarios of high and low number of conditionally beneficial alleles, $X = 100$ and $X = 2$, while keeping their total selective advantage constant. We performed 250 replicates with different sets of parameters for each scenario. We note that despite the inverted haplotype being advantageous, it was sometimes lost due to drift or genetic load. In those cases, we reset the simulation to its state before the inversion and choose a different random number seed.

Bins of highly correlated SNPs often had a low number of polymorphic sites and suffered from high variance. We therefore calculated the probability of observing a ratio as extreme as or more extreme than the observed ratio given the number of SNPs in a bin. Specifically, we performed a one-tailed test based on the distribution of dN/dS ratios expected under strict neutrality. For each bin, we counted the number of polymorphic sites, $P$, and simulated distribution of N/S ratios by sampling from a binomial distribution, where $N \sim Binomial(P,\, p_n)$ and $S \; = \; P \, - \, N$. We set $p_n \; = \; 0.691$ to match the chosen value in the forward-in-time simulations. Additionally, we examined how the number of conditionally adaptive loci influenced the frequency with which the inversion was lost due to drift. To do so, we simulated data for increasing amounts of total selective advantage and measured how often the inversion was lost. We performed a total of 600 replicates for each scenario. Results are shown in fig. S45.

#### Gene ontology enrichment analysis

Gene ontology (GO) enrichment tests were performed on sets of genes in the vicinity of top-ranking inversion-correlated SNPs (ICS; correlation coefficient $|r| \geq 0.99$) independently per inversion, using the R package topGO v2.38.1 (*140*). The gene sets were tested against all genes in the SNP annotation (see *Short read sequencing, alignment, SNP detection, and statistical phasing*). For consistency across analyses, the test genes were extracted from the fAstCal1.2 Ensembl release version 99, matching the prebuilt database used by snpEff. To this end, a gene table was retrieved from BioMart using the Bioconductor R package biomaRt v2.60.0 and genes with transcription start site within 25 kbp distance from the target ICS were extracted. The number of test genes per inversion ranged between 46% and 53% of the total number of genes within the inversion, defined as the region between the breakpoints 'start_left' and 'end_right' (table S8).

Genes were mapped to GO terms using the zebrafish (*Danio rerio*) annotation package (org.Dr.eg.db) from Bioconductor. The nodeSize parameter was set to 5. Statistical overrepresentation of GO terms was calculated performing Fisher's exact test, using the weight algorithm (object class 'weightCount'). The significance values are reported without multiple testing correction.

#### Selection tests (NI, DoS, dN/dS)

To assess selective forces acting in the inversion regions we calculated the neutrality index (NI = dSpN/dNpS), the direction of selection statistic (DoS = dN/(dN + dS) − pN/(pN + pS)) (*141*) and the dN/dS ratio across inverted regions for different bins of SNP-inversion correlation coefficients r (see section *SNP–inversion-correlation*). While for dN, dS all SNPs annotated as (non)synonymous in the respective regions by SNPeff with an r value in the respective bin were used, SNPs with weak inversion correlation ($|r| < 0.04$) were used for pN and pS. Estimation of dN/dS ratio was based on the Jukes-Cantor substitution model; transitions were assumed to happen at five times the rate of transversions. Under these assumptions we expect 30.9% of synonymous and 69.1% of nonsynonymous substitutions to happen under purely neutral evolution of the standard genetic code (*142*). This ratio was used for normalisation of the observed amount of synonymous and nonsynonymous substitutions in r bins. When calculating neutrality statistics in r bins, we found that estimates for intermediate correlation bins were noisy due to a low density of SNPs with intermediate r values (most r values were either small or large in absolute value). To overcome this issue, we calculated neutrality statistics in widening bins of r values, i.e., {r ∈ [-1, 0.99], [-1,0.98], .., [-1,0], [0,1], .., [0.98,1],[0.99,1]}. We consider this approach as conservative, because it smoothens the signal coming from high r SNPs across bins.

#### Expression patterns in zebrafish homologs

To obtain a better understanding of target genes' functional categories, we used the Daniocell zebrafish single cell expression database (*143*). Out of the 441 genes with at least one nsICS, we selected those

which have orthologs in the zebrafish genome (315 genes). For these we gathered data on tissue-specific expression in zebrafish embryos at developmental stages between 3 and 120 hours post fertilisation. For each gene we recorded whether it is highly expressed in 19 initial tissues. For each gene one tissue with the highest mean expression level was recorded. Applying this approach we gathered tissue-specific expression patterns of 315 genes (table S19).
To check for significant tissue-specific enrichment patterns across the 315 target genes we additionally assessed 12,873 random zebrafish orthologs for comparison (table S20). Then we divided gene sets into groups according to inversion and haplotype. If a gene had more than one nsICS mutation, it was counted once per nonsynonymous nsICS mutation. In some analyses we grouped 19 tissues into 3 sub-categories: neurosensory, coloration and other. Neurosensory category included neural, eye, otic and taste tissues. Coloration included pigment cells, fin and epidermis. All the other tissues were grouped into "other". Finally, we compared the proportion of genes expressed in target and random gene sets of each tissue (or tissue group) independently (Fisher test, Benjamini-Hochberg false discovery rate correction). Results are presented in figs. S46, S47, S48.

#### Association between bower building and inversion state

Based on a literature search we categorised each *benthic* species with information on bower building behaviour as either 'pit-digging' or 'castle-building' (ignoring more complex architectures like 'pit-castles') (n=56, table S21). We intersected these with our sequenced species and pruned the *benthic* 684 sample tree (data S1) to a set of 48 individuals, each representing a pit-digging or castle-building species. We encoded the presence/absence of the five focal inversions as binary traits, considering an inversion present in a species if among all genotyped individuals (tables S7, S11) at least one was heterozygous. Along with bower type, we plotted these traits onto the pruned tree (fig. S50).
We then used BayesTraits (*144*) to test for associations between the presence/absence of each focal inversion and bower architecture, taking into account phylogenetic relationships as well as uncertainty. We used the same 100 kbp window trees that were computed earlier to construct the 684 sample *benthic* tree (data S1), and excluded all subtrees with with ≤ 80000 sites unmasked, the inversion chromosomes and the extremely repetitive chromosome 3. We then pruned all remaining 4,219 subtrees to the same set of 48 samples and combined them in a NEXUS file according to BayesTraits specifications. All tree operations were performed in python with the ETE toolkit (*94*). To test for correlation between bower type and inversion presence/absence, we used BayesTraits to compare the fit of two continuous-time Markov models for each inversion, one assuming independent character evolution ('Discrete: Independent' model) with bower type, the other assuming dependent evolution ('Discrete: Dependant' model). For each run we selected 'MCMC' as analysis method, setting all rate priors to an exponential with mean of 10 ('PriorAll exp 10'), running each chain for 5,000,000 iterations ('Iterations 5000000') and discarding the first 500,000 iterations as burnin ('BurnIn 500000'), and used the stepping stone sampler with 1000 stones of 100,000 iterations each ('Stones 1000 10000') to estimate the marginal likelihood. For each inversion, we then converted the marginal likelihoods of both models into a Log Bayes Factor (LBF) using the formula

$$LBF = 2\,(log\ marginal\ likelihood\ dependent\ model - log\ marginal\ likelihood\ independent\ model).$$

LBF were then interpreted to assess trait correlation.

#### Allele-specific expression analysis

To identify genes under allele specific expression (ASE) for inverted and non-inverted haplotypes among the 11 RNA sequenced male *C. chrysonotus*, we first calculated for each SNP in coding sequences an inversion correlation coefficient analogous to the SNP-inversion correlation analysis above, but restricted to 34 whole genome-sequenced *C. chrysonotus*. We then identified for each individual and for each gene within the chromosome 11 inversion region (including 500 kbp left and right of the breakpoints) the heterozygous SNP with the highest correlation coefficient and used the number of RNAseq reads aligned to the same region containing this SNP as a proxy for allele specific expression. Gene × individual combinations with $|r| < 0.2$ were set to missing. Significance of allele specific expression values was calculated for each gene across all included individuals based on allele counts. We used a Wilcoxon signed-rank test as implemented in scipy.stats.wilcoxon with nan_policy='omit' and zero_method="pratt" and then corrected for false discovery rate using the Benjamini–Hochberg procedure as implemented in statsmodels.stats.multitest.fdrcorrection (*145*). This analysis was performed both separately for each tissue (which did not yield any gene with FDR $< 0.05$) and for all tissues jointly (fig. S52, table S24).

#### RNAseq and differential gene expression

RNA sequencing reads were generated separately from 5 tissues (brain, gills, gonads, liver, muscle ) for 24 individuals (11 *C. chrysonotus* heterozygous for the chromosome 11 inversion, 11 *C. mloto* homozygous inverted chromosome 11 inversion) as follows (all procedures were conducted on ice, unless otherwise specified): samples stored in RNAlater were thawed from -80 °C and transferred to 500 µL of Trizol. For each sample, 100 mg of 0.1 mm zirconia/silica beads (Stratech) were added before homogenization using a TissueLyser II (Qiagen) for 120 seconds at 30 Hz. The samples were then topped up to 1 ml with chilled Trizol and allowed to rest for 5 minutes. Next, 200 µL of chloroform (ThermoFisher Scientific) was added and the samples were vigorously shaken for 15 seconds, briefly vortexed and incubated at room temperature for 15 minutes. Samples were then centrifuged at 300 x *g* for 20 minutes at 4 °C. The supernatant was carefully transferred to a fresh tube and further processed using the Direct-Zol RNA Purification Kit (Zymo) according to the manufacturer's instructions. Quality and quantity of the extracted total RNA were assessed using Qubit (RNA BR assay, Agilent) and Tapestation (Agilent). RNA library preparation (75 bp, paired-end) and total RNA sequencing (Illumina HiSeq 4000) were performed at the Wellcome Sanger Institute. The differential expression analysis was conducted applying the RASflow pipeline (*146*). In short: trimmed reads were aligned with HISAT2 (*147*) to the *A. calliptera* reference genome (fAstCal1.2), count matrices were obtained with featureCounts (*148*) and analysed in the DESeq2 R package (*149*). We used the following comparisons: gonads vs. all other tissues in *C. chrysonotus* with "individual" as confounding factor and *C. chrysonotus* vs *C. mloto* for each tissue separately. As significance thresholds we used absolute log2FC $> 1$ and FDR adjusted p-value $< 0.01$.

#### Gene haplotype analysis

To visualise haplotypic differences in genes of interest we extracted their coding sequence based on the fAstCal1.2 annotation (Ensemble genome database release version 99). Using bcftools consensus (*79*), and specifying fAstCal1.2 as the reference, we separately extracted the coding sequence for up to two samples per included species and additionally for the ancestral sample (see ancestral state inference) from the re-phased inversion region VCF files (see inversion haplotype rephasing). If a gene was annotated on the reverse strand we reverse complemented it. Next, we translated nucleotides into amino acid sequences using seqmagick convert (v0.8.6) (https://github.com/yulab-smu/seqmagick) with the option --translate dna2protein. After concatenating the amino acid sequences for all samples per gene, we created UPGMA trees with FastTree 2.1 (*150*) applying the -nosupport option. Finally, we rooted the tree to the inferred ancestral sequence using Newick utils (*151*) function nw_reroot and visualised haplotype trees in Haploviewer (http://www.cibiv.at/~greg/haploviewer.shtml). Results are shown in fig. S49.

### Supplementary text

#### Text S1: Comparative genomics and small inversions

We initially identified five chromosome-scale polymorphic inversions on the basis of the regionally restricted, but consistently aberrant patterns in genetic structure among the *benthic* group of Malawi cichlids, resulting from strong recombination suppression.

*Ancestral inversion state inference*

Our examination of introgression patterns and evolutionary history suggests that *A. calliptera*, which is also used as the mapping reference in our Malawi variant callset, carries none of the focal inversions. This is supported by PC analyses of the inversion regions (fig. S21), where *A. calliptera* consistently clusters with putatively non-inverted samples. To further confirm the absence of the respective inversions in *A. calliptera*, we aligned publicly available genome assemblies of two outgroup species (*Pundamilia nyererei*, a representative of the Lake Victoria sister radiation, and *Oreochromis niloticus*, the Nile tilapia) to our Hi-C-(re-)scaffolded *A. calliptera* assembly (materials and methods, fig. S21 and fig. S22). Despite multiple other, smaller rearranged segments, none of the focal inversions were present in these comparisons. We therefore conclude that *A. calliptera* possesses the ancestral configuration with respect to the chromosome-scale inversions identified here. We furthermore suggest that the majority of the numerous smaller rearrangements are likely misplaced or misoriented contigs in the two outgroup genomes, which did not use Hi-C technology and are more comparable to our astCal1.2 assembly in terms of structural integrity.

*Inversion prevalence*

To further examine the characteristics and prevalence of the focal inversions, we generated new Hi-C data and chromosome-level assemblies for one representative of the remaining four Lake Malawi cichlid clades *Rhamphochromis*, *Diplotaxodon*, *benthics* and *mbuna* (table S4). To examine structural rearrangements, we generated *A. calliptera*-referenced pairwise genome alignments and Hi-C maps, respectively (figs. S9 to S12, S23). We furthermore generated linked reads (haplotagging) for 23 representatives of 9 benthic species (fig. S14), which, based on our radiation-wide inversion genotyping were expected to carry inversions on chromosomes 2, 9, 11 and 13 (table S2). In addition, we designed fluorescence in situ hybridization (FISH) probes to physically confirm inversions on chromosomes 9 and 11 between *A. calliptera* and *Au. stuartgranti* (*deep benthic*), both species were also used in our interspecific cross (Fig. 2C, fig. S8).

While *Rhamphochromis* and *mbuna* featured none of the five inversions, the *benthic* representative carried the full chromosome 9 and 11 inversions, which was apparent from whole genome alignments, Hi-C maps, linked reads and FISH. These comparative analyses furthermore revealed that the inversion on chromosome 11 is a compound inversion of two approximately equally sized inverted segments. Since both segments behave identical in all analyses we conducted, we treat the entire region as a single inversion and suggest that both respective inversion events occurred together or over a short evolutionary period. Consistent with local PC analyses (fig. S21), the *Diplotaxodon* assembly also carried the inversions on chromosomes 9 and 11, which was reflected in the respective whole genome alignment and Hi-C map.

We furthermore confirmed the presence of the focal inversions on chromosomes 2 and 13 using haplotagging linked reads. Chromosomes 2 and 13 inversions were present in 1 and 4 samples, respectively, and matched our expectations from PCA- and PCR-based inversion genotyping for the species (table S2). We were unable to access high molecular weight samples for putative carrier species of the chromosome 10 inversion, which is restricted to *deep benthics*.

Based on the pairwise genome alignments between our curated Hi-C based assemblies we identified several additional smaller inversions. Among them, the three largest ones are located on chromosomes 2 and 20.

##### *A nested inversion segregates on chromosome 2*

The small inversion on chromosome 2 is only present in the *deep benthic* assembly, and located within the larger pericentric chromosome 2 inversion region, directly adjacent to the centromere (fAstCal1.2 coordinates: 12,919,679 to 16,823,069 bp). This region coincides with a region on chromosome 2 where the inversion genotypes segregate less distinctly in windowed PC analyses compared to the rest of the chromosome 2 inversion (fig. S15), and compared to the other four large inversions (Fig. 2A). Furthermore, recombination is locally reduced in the corresponding region on chromosome 2 in our interspecific cross (Fig. 2E, upper panel). Taken together, this suggests that this region is structurally complex and contains one or possibly several smaller inversions that segregate independently among subsets of *benthic* samples. Unfortunately, due to the strong heterogeneity in local genetic structure we were unable to determine inversion genotypes of individual samples with sufficient certainty.

Notably, chromosome 2 has also been implicated in an earlier study that investigated the genetic basis of bower building in *benthic* Malawi cichlids, and it is conceivable that genetic variants on one or several nested inversions could impact bower building behaviour. Remarkably, the entire region is absent from the included Lake Victoria genome assembly (fig. S22), while it appears to be present in *Oreochromis niloticus* (fig. S23), an outgroup to both radiations. While Victoria cichlids are not known to build elaborate bowers (*152*, *153*), *Oreochromis* species do (*154*), including *Oreochromis karongae* which is native to Lake Malawi. Possibly, structural rearrangements in this region originate in the pelagic subradiation and introgressed into *benthic* clades. While we think this structurally diverse region is very interesting, we see only limited room for further analyses using short-read based variant calls due to the structural heterogeneity in that part of the genome. We suggest that future studies with a focus on the genetic basis of bower building behaviour should consider this region on chromosome 2 specifically. This would require additional ecological or behavioural data on bower building, as well as more chromosome-scale assemblies to characterize the presence/absence and locations of rearrangements in the respective region.

##### *Two adjacent inversions on chromosome 20 could have played a role in the evolution of pelagic clades*

We identified two adjacent inversions on chromosome 20, at astCal1.2 coordinates 4,788,189 to 8,315,641 bp and at 8,331,131 to 12,943,513 bp. The first segment is inverted in *Rhamphochromis*, *Diplotaxodon* and the *deep benthic* individual relative to *A. calliptera*, while the second segment is not inverted in *Rhamphochromis*, but in *Diplotaxodon* and in the *deep benthic* representative (fig. S9, fig. S10, fig. S11, fig. S25). Both outgroup species lack these inversions (fig. S22, fig. S23). Since we did not detect these rearrangements as aberrant regions in other analyses (e.g. Fig. 2A) but observed strong recombination suppression in our interspecific cross (Fig. 2E, upper panel), we suggest that they are entirely or largely fixed for one or the other orientation within extant Malawi cichlid clades.

To further explore the possibility of an inversion polymorphism that went undetected in our previous analyses we inspected PC1 to PC4 of all *benthic* individuals for the respective compound inversion region and compared the segregation patterns to that of the combined flanking regions on the same chromosome (fig. S16A-D). On PC4, we identified a pattern among *shallow rocky Aulonocara* (part of the *deep benthic* clade) which could be consistent with a polymorphic inversion (fig. S16D). For increased spatial resolution of this focal group we re-plotted PC3 and PC4 for the 27 *shallow rocky Aulonocara* individuals while omitting all other samples (fig. S16E,F). We furthermore conducted a windowed PCA of chromosome 20 for the same 27 individuals and visualized PC1 and PC2 (fig. S16G and fig. S16H). We conclude from this extended analysis that we could not find evidence supporting an inversion polymorphism in *benthic* clades and that the observed aberrant pattern among *shallow rocky Aulonocara* most likely reflects normal variation in genetic structure along the genome (fig. S16E-H). While long read-based genome assemblies of multiple *shallow rocky Aulonocara* individuals could be generated for absolute certainty, we believe that either outcome would not change the main conclusions from this study, since only a small subset of *deep benthic* taxa is affected.

Based on available information and our revised model for the Lake Malawi radiation (Fig. 3B), we hypothesise that the first segment on chromosome 20 inverted and got fixed in the ancestor of pelagics (since it is shared between *Rhamphochromis* and *Diplotaxodon*), while the second segment inverted and got fixed in the ancestor of *Diplotaxodon*, after the split from *Rhamphochromis*. Both segments were then passed on from *Diplotaxodon* to *benthics*, likely as part of the hybridization event at the base of *benthics* which also explains the presence of the chromosome 2 and 11 inversions in *benthics*.

*Centromere localisation and structural differences between inversion haplotypes*

Querying our *Diplotaxodon* high quality PacBio HiFi-based genome assemblies for centromeric repeats allowed us to locate and then project centromeric regions to the reference genome, revealing that only the large inversion on chromosome 2 is pericentric (Fig. 2A, fig. S17, table S6). Analysing the large insertion sequences between *A. calliptera* and *Au. stuartgranti* assemblies (differentially fixed for chromosome 9 and 11 inversion states) for occurrence of transposable elements, we found a slight trend that inversion regions tended to harbour fewer recently inserted and deleted DNA sequences compared to the rest of the genome (fig. S18) but this trend was not consistent across superfamilies (LTR, DNA, LINE or SINE) and inversions (fig. S19). Given the sex-linkage of the chromosome 9 and 11 inversions (Fig. 5), we also checked whether any of the indel sequences in inversion regions correspond to any of 124 genes that have previously been previously implicated in sex-determination, which would support the translocation of sex determination genes. However, we found no significant matches.

#### Text S2: Evolutionary history of inversions

To better understand the evolutionary histories in inversion regions, we estimated genetic divergence times both between *benthic* lineages and from the other major Malawi clades for the five inversion regions (both the inverted and non-inverted haplotypes) as well as for the remaining non-inverted regions of the genome (Fig. 3A, fig. S26). We also rephased heterozygous SNPs (see materials and methods) in inversion heterozygote individuals using SNP allele frequencies in the different inversion genotypes and built consensus phylogenies for all haplotypes in all inversion regions (fig. S29 to fig. S33). Together, the results of these analyses imply that the evolutionary histories of all five inversion regions were shaped by multiple introgression events (Fig. 3B, fig. S34).

Genome-wide phylogenetic trees for Malawi cichlids (e.g. ref. (*31*), Fig. 1, data S2) suggest that *benthics* are a sister clade to *A. calliptera* and *mbuna*. However, in the top row of Fig. 3A we see that both *shallow* and *deep benthics* are closer to *Diplotaxodon* than to *Rhamphochromis*, and closer to *A. calliptera* than to *mbuna*. These results are inconsistent with the genome-wide trees. It was already established that there is evidence for gene flow from *Diplotaxodon* into the *benthics* (*31*). However, here we suggest that the most likely resolution of these observations is that the *benthics* arose through admixture between the *Diplotaxodon* and *A. calliptera* lineages after their splits from *Rhamphochromis* and *mbuna* respectively, as shown in Fig. 3B. This hybrid *benthic* origin model changes the relative order of divergence of *benthics* and *mbuna* from the *A. calliptera* stem, which is a natural consequence of the inability of the binary tree phylogeny to account for admixture. In the context of this revised model of the origin of the *benthics*, the sharing of the chromosome 9 and 11 inversions (as well as the chromosome 20 compound inversion, see fig. S9 to fig. S12) between all *Diplotaxodon* and some *benthics* is most parsimoniously explained by them having arisen in an ancestor of *Diplotaxodon* and passing into the *benthics* in their founding admixture event, as indicated by Ⓐ in Fig. 3B.

Considering subsequent events, the chromosome 9 region is fixed inverted in most *benthic* species except for the *eukambuzi* and a few others where it is heterozygous (Fig. 1). However, it is striking that the non-inverted state found in these few shallow *benthics* is much more divergent from the rest of the Malawi radiation than any other inversion haplotype and the rest of the *benthic* genome (Fig. 3A, row 2). This suggests that this haplotype in the non-inverted orientation re-introgressed into shallow *benthics* from a lineage that is phylogenetically outside the entire Malawi radiation. To search for the origin of this haplotype, we analysed a second SNP callset containing representatives of all major Malawi clades as well as a wide variety of related riverine and lacustrine cichlid species ("haplochromines") (*32*) (fig. S28, table S12, materials and methods). Computing ABBA-BABA tests (table S13) revealed excess allele sharing of the non-inverted *benthic* chromosome 9 haplotype with a number of distantly related outgroup species, with the strongest signal being to *Pseudocrenilabrus philander*, one of the few outgroup species present in the catchment of Lake Malawi (D = 0.45, block-jackknifing z-score 6.5; FWER corrected p = $4\times10^{-9}$). We conclude that the chromosome 9 non-inverted *benthic* haplotype is not closely related to other Malawi haplotypes (whether inverted or not), but instead arrived in the ancestor of *eukambuzi* through admixture with a distantly related *Pseudocrenilabrus*-like cichlid lineage (Ⓑ in Fig. 3B) and spread to the few other species in which it is found by subsequent introgression.

Turning next to the chromosome 11 inversion, this is fixed inverted in the *deep benthics* and *shallow Lethrinops* and variably present in other shallow species, being mostly non-inverted in the *eukambuzi* (Fig. 1). Unlike in all the other inversion regions, the non-inverted haplotype is closest to *A. calliptera* (Fig. 3A, row 3), suggesting that it re-introgressed into shallow benthics from an *A. calliptera* ancestor or an extinct close relative, as indicated by Ⓒ in Fig. 3B. Although we cannot exclude that the non-inverted haplotype was retained since the admixture event at the base of benthics and subsequently independently lost in deep benthics and most non-eukambuzi shallow benthic lineages, this seems less parsimonious.

Furthermore, the lower heterozygosity (and nucleotide diversity) compared to the inverted haplotype (fig. S27) is consistent with a genetic bottleneck upon re-introgression. Signatures of gene flow from *A. calliptera* into the *shallow benthics* also exist outside of chromosome 11 where *shallow benthics* are closer to *A. calliptera* than *deep benthics* are (Fig. 3A, row 1). Furthermore the non-inverted haplotypes of the chromosome 2 and 13 inversions are closer to *A. calliptera* than the inverted haplotypes, which are present predominantly in the *deep benthics* and protected from introgression because of their orientation (Fig. 3A). The f-branch analysis in Fig. 3 of our earlier study (fig. S32) further supports this genome-wide gene flow, showing significant excess allele sharing of the ancestral branch of shallow *benthics* with *A. calliptera* relative to *deep benthics*. Finally, some *utaka* also received the *A. calliptera*-like non-inverted haplotype in the chromosome 11 region. Overall, consensus phylogenies and genetic divergence measures suggest that *utaka* are a complex (and variable) mixture of *shallow benthic* and other contributions, which also explains their non-monophyletic clustering in the genome-wide phylogeny (Fig. 1, figs. S36 and S37).

The remaining inversions on chromosomes 2, 10 and 13 are all common among *deep benthics*, and rare or absent among *shallow benthics* and *utaka*, suggesting that they rose to high frequency early in the deep *benthic* lineage (Fig. 3B). This is also consistent with the phylogenetic relationships within the inversion regions (fig. S33, fig. S31 and fig. S32), which suggest that the few cases of *deep benthics* with non-inverted states and *shallow benthics* with inverted states are due to a limited number of later gene flow events (fig. S34). Most of these events transmitted more than one inversion haplotype between *deep* and *shallow benthic* lineages and also genetic material outside the inversion regions, as seen in ABBA-BABA tests of excess allele sharing outside inversion regions (Fig. 3C) (*41*). This underlines that the sharing of inverted haplotypes among *deep* and *shallow benthic* species is generally not a result of random segregation through incomplete lineage sorting but due to gene flow. The most consequential of these transmission events are indicated in Fig. 3B. Specifically, inverted haplotypes introgressed from *deep benthics* into the *shallow Lethrinops*, which show traits similar to deepwater *Lethrinops* consistent with their taxonomic assignment to the same genus despite their distinct molecular phylogeny (Ⓓ in Fig. 3B). Furthermore, non-inverted chromosome 10 haplotypes introgressed from a shallow source into a broader *deep benthic* group of species (indicated by Ⓔ in Fig. 3B). Finally, the non-inverted haplotypes for chromosomes 2, 10, and 13 introgressed from some *shallow benthic* source into *shallow rocky Aulonocara* (Ⓕ in Fig. 3B), where they now appear to be fixed. Further events, shown in fig. S34, are qualitatively similar to these in that they mostly correspond to inversion transmissions between *deep* and *shallow benthic* species that were collected at a similar depth (fig. S35). This suggests that non-inverted (respectively inverted) haplotypes tend to carry alleles favourable to shallow (respectively deep) environments, which we address further in the main text.

#### Text S3: Inversion-correlated SNPs (ICS) and selection

To describe selection forces that affected early inversion evolution, we focussed on SNPs with high inversion correlation coefficients (ICS) as those are likely to have been present early on inversion haplotypes potentially affecting their evolution.

##### *Impact of different evolutionary forces on selection statistics*

First, we explore how the observed patterns of selection statistics such as dN/dS, neutrality index (NI) and direction of selection (DoS) (*141*) are affected by different evolutionary forces. It is important to note that all these statistics are affected by both positive and negative (purifying) selection, but also by genetic drift. Genetic drift can be caused by different processes, most prominently demographic changes (small effective population size) but also by linked selection (i.e., the effect of positive or negative selection on linked (neutral) sites).

The effects of selection and drift are most straightforward for dN/dS. In the absence of selection, the expectation for (normalised) dN/dS is 1.0, because synonymous and non-synonymous mutations are equivalent. Positive selection increases dN/dS, negative selection reduces dN/dS. Drift reduces the efficacy of either type of selection, and will therefore drag the value of dN/dS closer to 1.0, whether it is >1.0 (excess of positive selection) or < 1.0 (excess of negative selection). It is important to note that while dN/dS < 1.0 is possible in the presence of positive selection (if negative selection is dominant), dN/dS > 1.0 is not expected in the absence of positive selection (*43*, *155*). Of course, depending on the number of N and S mutations, dN/dS > 1.0 is likely to be the result of several positively selected mutations, as a single mutation would generally not shift the balance relative to deleterious mutations.

McDonald-Kreitman-type statistics such as NI and DoS use both divergence (dN, dS) as well as polymorphism (pN, pS) measures. Although these statistics are widely used as evidence of positive selection (when NI < 1 or DoS > 0), it is important to realise that increased drift (reducing the efficacy of purifying selection) can also lead to NI < 1 or DoS > 0 (*42*). Since, as we discuss in the following, there are several reasons why inversions are expected to experience more drift than other genomic regions, we do not base any of our conclusions on the NI and DoS, but rather on raw dN/dS for which, as explained above, dN/dS > 1.0 is only consistent with adaptive evolution. We merely display NI and DoS for completeness (fig. S43) since these are commonly used statistics.

##### *The impact of drift on inversions*

Inversions are expected to be subject to drift (and thus reduced efficacy of purifying selection) for several reasons. First, young inverted haplotypes are expected to exist at low population frequencies for some time after their birth, thus being present mainly in heterozygous state with suppressed recombination. These circumstances prevent effective purifying selection and increase the genetic load. In a broad range of evolutionary scenarios, this process prevents inversion from reaching high frequency or fixation, and either leads to their maintenance at intermediate frequencies through heterozygote advantage (if deleterious mutations are recessive), or to their loss (*62*). Second, if inversions carry positively selected variants and therefore rapidly raise in frequency, linked selection will drag to high frequency all variants present on the initial inversion haplotype. This will affect both variants that were rare and variants that were common in the original population, but since rare variants are more likely to have deleterious fitness effects it will lead to an accumulation of genetic load relative to the original population. Since these variants are "fixed" in the inverted population, purifying selection will only ever be able to remove them if recombination happens with a non-inverted haplotype (if we ignore back mutations). Third, if the inversion haplotype grows rapidly, purifying selection against new deleterious variants arising during this growth phase will be less efficient. This will increase the relative amount of deleterious variants on the inverted haplotype. Such deleterious variants appearing post-inversion will generally not fix in the inverted population (the chance loss of the alternative allele is low in a growing population). Therefore, once the haplotype is at high frequency and effectively recombines within the inverted population, purifying selection can remove these variants, thereby reducing genetic load.

For the inversions discussed in our paper, their phylogenetic distribution (Fig. 1) makes it clear that they rose to high frequency (and probably fixation) rapidly in ancestral lineages of present day clades. Given the general fitness penalty of inversions due to genetic load (*62*), this alone is a strong argument that inversion haplotypes experienced strong positive selection. Furthermore, rather than segregating at low frequency for a long time and thereby accumulating genetic load through suppressed recombination, inversion haplotypes likely were (nearly) fixed in the populations/species where they occurred for much of their existence. Therefore, we believe that the rapid frequency increase of the inversion haplotype (points two and three in the last paragraph) will be the main factor contributing to drift (and thus relaxed purifying selection) in the evolution of the Malawi cichlid inversions rather than suppressed recombination (point one in the last paragraph).

*In-silico expanded haplotypes neither show dN/dS > 1 nor tissue-specific expression of nsICS*

To test our expectations for selection statistics against an extreme scenario of expansion of an initially rare haplotype, we performed the following in-silico experiment. We picked a random haplotype from our actual SNP data of 656 benthic species with coverage > 10 and copied it 656 times to create a hypothetical inverted population (designating all individuals present in the actual SNP callset “non-inverted"). We then calculated inversion correlation for the resulting SNP genotype just as we did for real inversions, performed selection tests and identified ICS (referred to as artificial or aICS in the following).

Consistent with our expectations, we found that in such a scenario, while NI and DoS attained values below 1.0 and above 0.0, respectively, that dN/dS did not become > 1.0 (fig. S44). For all eight randomly selected benthic species for which we picked a haplotype to expand, dN/dS stayed below 0.6 for positive inversion correlation coefficients. For highly negative correlation coefficients (corresponding to rare ancestral alleles on the selected haplotype) dN/dS values had more variance due to low SNP numbers in these bins, but dN/dS still remained < 1.0. This leads us to conclude that dN/dS > 1.0 is indeed not expected to result from the strong expansion of a haplotype alone, without the presence of positive selection on several non-synonymous variants.

We further applied our single-cell tissue expression analysis (applied to ICS in Fig. 4C, figs. S46 to S48) to aICS. We did this to gain insight into whether a strong haplotype expansion (as expected for real ICS) could cause the observed overrepresentation of expression in specific tissues even in the absence of selection (e.g., through a bias in the variants present on the the initial, rare inversion haplotype, or through biases in gene length). Testing this for all chromosomes and in each of the eight randomly selected benthic individuals used to calculate aICS (see fig. S44), we found that only a very small fraction of the tests (mean 0.9%) yielded significant tissue enrichment (tables S30 and S31). We take this to conclude that our approach to calculate tissue enrichment for nsICS is not intrinsically biased by the variant composition following the rapid expansion of a rare haplotype or by gene length.

*Observed dN/dS ratios are consistent with positive selection on multiple sites in time-forward simulations*

To further investigate the effect of the strength of selection and of the number of selected sites on inversion maintenance and dN/dS ratio, we used SLiM 4.3 (*138*) to perform forward-in-time simulations of the evolution of inversion haplotypes carrying beneficial and deleterious mutations (see materials and methods). The point of these simulations was not to explore the full large parameter space of possible evolutionary models or to precisely match the (unknown) evolutionary history of the Malawi cichlid inversions, but to qualitatively explore the effect of the number of sites under adaptive evolution on dN/dS ratio. We found that for a relatively large number of positively selected mutations (X=100), relative dN/dS increased significantly above 1.0 for high SNP-inversion correlation bins (ICS). In contrast, for a low number (X=2) of sites under adaptive evolution (the same total selection coefficient), dN/dS was not significantly elevated above 1.0 (fig. S45). The simulations confirmed that generally there is a high chance that inversions get lost due to drift or genetic load and that the chance of inversions rising to high frequency increases with the strength of positive selection (fig. S45C). Overall, these simulations support our conclusion that the observed dN/dS ratios are likely the result of positive selection at multiple sites in each inversion haplotype.

#### Text S4: Signals of adaptation in inversions

*Widespread signatures of adaptation on inversion haplotypes*

To identify SNPs and genes involved in early inversion haplotype evolution, we computed correlation coefficients and significance scores ($-\log_{10}$ p-value) between SNP and inversion genotypes. We expected the most highly inversion correlated SNPs ("ICS", Fig. 4A) to correspond to those present on the early inversion haplotypes that might have driven the early evolution of these haplotypes.

Positive ICS values correspond to derived alleles being at high frequency on the inverted haplotype, and negative ICS values to ancestral alleles being on the inverted haplotype (Fig. 4A). Consistent with the distinct evolutionary histories of the five inversions, the distribution of derived alleles between inverted and non-inverted haplotypes differed among chromosomes, with chromosomes 9 and 11 showing an excess of derived SNP alleles on the non-inverted, "re-introgressed" haplotype (negative ICS), while chromosomes 2, 10, and 13 showed an excess of derived SNP alleles on the inverted, "*deep-benthic*" haplotype (fig. S42).

Adaptive evolution can occur through selection on protein sequence (coding mutations) or through heritable changes in gene expression, the latter often accomplished through cis-regulatory mutations in the vicinity of genes. To identify biological functions associated with both processes we first performed functional enrichment analyses on genes near ICS (see materials and methods). This should capture both protein coding and regulatory adaptation. In a second step, we further investigated genes with high nonsynonymous ICS ("nsICS genes", Fig. 4A) and their expression patterns in zebrafish development. Finally, to understand whether nsICS genes fall into regions of overall increased haplotype divergence we inspected regions of high average ICS (top 0.5% quantile of mean ICS scores across windows of 100 SNPs, "windowed ICS outliers"; Fig. 4D).

In analysis of nsICS genes we relied on the raw number of nsICS per gene and "density" of nsICS, which is the number of nsICS normalized by gene length (measured in nsICS per nucleotide) (table S25). We also tested dependency of nsICS number from gene length and found no significant connection for inversions at chromosomes 9, 10, 13. For inversions at chromosomes 2 and 11 there was significant dependency (F-statistic for linear regression, $p = 4\times 10^{-4}$ and $p = 3\times10^{-11}$). However, for the chromosome 2 inversion, the dependency becomes insignificant if an extremely long gene (*ahnak*) is removed from the sample.

##### *Inversions contribute to sensory adaptation and behaviour*

Given that all five inverted haplotypes are found on average at higher frequencies in deepwater-living species (fig. S35), we hypothesised that they could have contributed to niche divergence along a depth gradient. Specifically, we expected relevant sensory and physiological adaptations related to changes in light, oxygen, and hydrostatic pressure as observed in many organisms (*45–47*), including cichlids (*31*, *48*, *49*). Consistent with this hypothesis, gene ontology (GO) enrichment analyses on zebrafish homologs of cichlid genes near ICS (see materials and methods) showed sensory system-related categories to be enriched in each of the five inversions. Enrichments included, for example, "otic vesicle morphogenesis" (chromosome 9); "closure of optic fissure" (chromosome 11); "sensory perception of smell" (chromosome 10); "inner ear auditory receptor cell differentiation" (chromosome 2); "otic placode formation" and "embryonic camera-type eye morphogenesis" (chromosome 13) (table S23). Some of the significant genes in these GO categories are involved in early developmental pathways during embryogenesis and play key roles in the formation of the eye (*sfrp5*, *sox4b*) (*156–158*), and the inner ear and lateral line system (*pax2a*, *eya1*) (*159*, *160*) in zebrafish (Fig. 4C).

While the GO-based approach, which included potentially regulatory non-coding ICS, identified predominantly developmental processes, we next focused on the 330 genes with at least one amino acid changing mutation with highly significant inversion correlation ("nsICS"; $|r|>0.86$, $p<10^{-200}$, Fig. 4A). nsICS genes are strong candidates for adaptive evolution. To obtain insights into potential functional roles of these genes we investigated the expression levels of their homologs across different tissues in single cell RNA sequencing data of developing zebrafish (Daniocell database (*44*)), assuming that tissue-specific expression patterns in zebrafish are indicative of the genes' functional roles in cichlids. We found that nsICS genes are significantly more likely to be highly expressed in a group of neural and sensory tissues compared to randomly selected genes (fig. S48, table S22). This pattern is significant for both inverted and ancestral haplotypes (FDR corrected $p=6x10^{4}$ and $p=0.02$), and visible for separate inversions (fig. S47, table S21). Specifically, genes with nsICS are significantly highly expressed in neural, eye and otic tissues (FDR corrected $p=6x10^{4}$, $p=0.02$ and $p=0.04$, Fig. 4C and D, fig. S46, table S18). Interestingly, enrichment in neural, eye and otic tissues is significant for genes with nsICS derived on inverted haplotype (FDR corrected $p=1x10^{3}$, $p=0.01$, $p=0.04$, fig. S47, table S21), but not on ancestral haplotype.

Among the 83 genes with two or more nsICS were a cone arrestin (*arr3a*), a guanylyl cyclase (*gucy2d*) and a gamma-aminobutyric acid (GABA) A receptor (*gabra6a*), all three expressed in the retina (*161*, *162*). Both *arr3a* and *gucy2d* are involved in signal transduction in photoreceptor cells and have previously been shown to share signatures of deepwater adaptation between *Diplotaxodon* and *deep benthics* (*31*). Additional strong candidates for depth adaptation among the top nsICS genes are tectorin-alpha (*tecta*), involved in otolith tethering (*163*) and myosin 7ab (*myo7ab*), orthologous to the human *myo7a*, implicated in hearing loss (*164*). Together, our results show that all five inversion chromosomes harbour (ns)ICS genes relevant for sensory perception and/or the development of sensory structures (Fig. 4C-D). Sensory systems mediate sound perception, mechanoreception and vision, essential for navigation, communication (territoriality, courtship) and the detection of prey in cichlid fishes (*63*), making them important targets of selection (*64*).

Following up on the overrepresentation of nsICS genes expressed in neural tissues on the inverted haplotype (fig. S41), we investigated the potential functional involvement of inversion genes in reproductive behaviour. We expect that exploration of deepwater environments with reduced light availability requires a wide range of behavioural adaptations and may affect general social signalling patterns as well as courtship behaviour and the building of mating platforms ("bowers") by males. We tested our set of nsICS genes for overrepresentation of neuroreceptors that have been previously associated with social (glutamate) and affiliative (oxytocin and arginine vasopressin/vasotocin, opioid receptors, dopamine, serotonin) behaviour, using a list of known genes (table S19) (*50*). Against a background of 14,436 total zebrafish orthologs, we found a significant overrepresentation of candidate neuroreceptor genes in our set of 330 nsICS genes (6 out of the 46 candidate genes, Fisher's exact test, p= 0.0011): three glutamate receptors (*gria3a* and *gria4b* on chromosome 10, *grik3*, on chromosome 11), one opioid receptor (*oprd1b* on chromosome 11), one dopamine receptor (*drd2l* on chromosome 11), and one serotonin receptor (*htr7a* on chromosome 13). Remarkably, genes and GO categories associated with neurotransmitter regulation are not only associated with fish social behaviour in general, but were specifically connected to bower building behaviour in Malawi cichlids (*51*). Bower building is a highly variable, sexually selected male extended phenotype in benthic Malawi cichlids (*52*), which was hypothesised earlier to be functionally connected to the existence of supergenes (*50*). Consistent with this, a previous divergence scan had identified conspicuous outlier SNP peaks for bower architecture within our chromosome 2 and 11 inversion regions. To investigate this further, we tested for correlation between bower architecture ('pit' vs. 'castle') and the presence and absence of each identified inversion in a likelihood framework, which revealed that inversion-bower-correlation is dominated by phylogenetic signal (fig. S50). That said, overall, our results suggest that adaptive alleles on inversion haplotypes affect sensory and behavioural pathways relevant in sexual selection and assortative mating.

##### *Inversions contribute to adaptation of the vascular system*

In addition to light-related selective pressures, a distinctive feature of the deepwater environment is its low oxygen availability. In Lake Malawi, dissolved oxygen drops below 4-5 ppm at ca. 120 m, a level which is generally considered to be stressful for fish without specific adaptations (*165–167*) – while *deep benthics* and *Diplotaxodon* are found down to the anoxic layer at about 200 m. Consistent with this, the most significant GO enrichment for the chromosome 13 inversion region concerns the regulation of angiogenesis (formation of blood vessels, functionally linked to responses to hypoxia stress (*168*)), while significant GO terms on other inversions include "primitive erythrocyte differentiation" (chromosome 11) or "regulation of heart morphogenesis" (chromosome 2) (table S23). Furthermore, several nsICS genes are expressed in vascular tissues in the zebrafish embryo (Fig. 4D). Angiotensin (*agt*), a master regulator of vasoconstriction, is among the five genes with two or more nsICS on chromosome 13. Interestingly, two of four angiotensin receptor genes annotated in the reference genome are also located inside inversion regions, *agtr1b* on chromosome 9 and and *agtr2* on chromosome 2. This implies that inversion haplotypes could have contributed to adaptation of the angiotensin-renin system to deepwater conditions. Finally, we note that one of the two cichlid orthologs of *epas1*, a gene widely implicated in high-altitude-related hypoxia adaptation in several organisms (*133*, *169*, *170*) is localised within the chromosome 13 inversion, consistent with a role in depth adaptation (despite not being represented among the most differentiated candidate genes). Together, pervasive signals of adaptation related to sensory and vascular systems across all five inversion haplotypes support a scenario in which multiple genes on several of these haplotypes contribute to adaptation to the deepwater environment (Fig. 4).

##### *Adaptations related to sex and reproduction*

Consistent with roles for inversions in sex determination (Fig. 5) we found many of the strongest candidate genes for adaptive evolution to be related to sex and reproduction (Fig. 4D). Specifically, there were candidates for sex determination or sex-related function among the 83 genes with two or more nsICS across all five inversions (table S23).

For example, the only two genes on chromosome 2 with five or more nsICS are *ahnak*, involved in sex determination in the cichlid *Oreochromis mossambicus (171)*, and *atrx* – X-linked and sex-reversing in humans (*172*). Furthermore, *smchd1* – a chromatin repression regulator responsible for ovarian differentiation in mammals (*173*) – is among the eight genes with two or more nsICS on chromosome 9. On chromosome 11, the only gene with five or more nsICS is *plecb* (fig. S49), whose ortholog mediates sex differentiation in mouse gonads (*174*). On chromosome 10, the only gene with four nsICS is *pof1b* (fig. S49) linked to premature ovarian failure in mammals (*175*). On chromosome 13, *gfra1*, which is a part of the Gdnf-Gfra1 pathway, is involved in spermatogonial stem cell renewal in mammals and teleost fish (*176*). In addition to harbouring multiple nsICS each, all but *pof1b* also fell into ICS outlier windows consistent with divergent selection having acted on an extended genomic region (table S24).

Given that the evolution of sex-determining regions often involves changes in gene expression between male and female haplotypes, we obtained transcriptomic data of five tissues (muscle, liver, brain, gills, gonads) for eleven males of *C. chrysonotus*, the *utaka* species in which all males were heterozygous for the chromosome 11 inversion while females were fixed for the inverted state, and investigated allele specific expression (ASE). Furthermore, lacking access to appropriate female samples, we obtained equivalent data for eleven male *C. mloto*, a congeneric species fixed for the derived chromosome 11 inversion state, to perform differential gene expression (DE) analysis between the two inversion states.

Consistent with the young age of the Malawi radiation, expression of most genes was similar across haplotypes in male *C. chrysonotus*, with a moderate bias towards lower expression of the Y-like non-inverted haplotype (of ancestral orientation) among genes with significant ASE (fig. S52), a pattern seen in many organisms (*57*). Several of the significant ASE and DE genes (FDR < 0.05) were implicated in sex determination, sex specific expression or gonad function in other (fish) species (see table S24). Of those, several overlapped with candidate ICS genes (e.g., *trim35-28* among top windowed ICS, *dscaml1* with two nsICS; Fig. 4D). Reciprocally, and in support of this, we found ICS to be significantly overrepresented among genes with significant allele specific expression (Fisher's exact test $p=1.6*10^{-7}$).

Interestingly, investigating ICS, ASE, and DE outliers also revealed genes related to meiosis, DNA repair, and gamete morphogenesis in the chromosome 11 inversion, with "negative regulation of meiotic nuclear division" also being the most significantly enriched GO category on chromosome 11. We suggest that this pattern could be related to the direct impact of inversion heterozygosity on chromosomal crossover and meiosis, with the development of gametes with aberrant karyotypes likely incurring a considerable fitness cost (*10*). Overall, our analyses support the presence of adaptive alleles contributing to reproduction, sex determination, or sex-specific function on all inversion haplotypes (Fig. 4D).

### Figs. S1 to S54

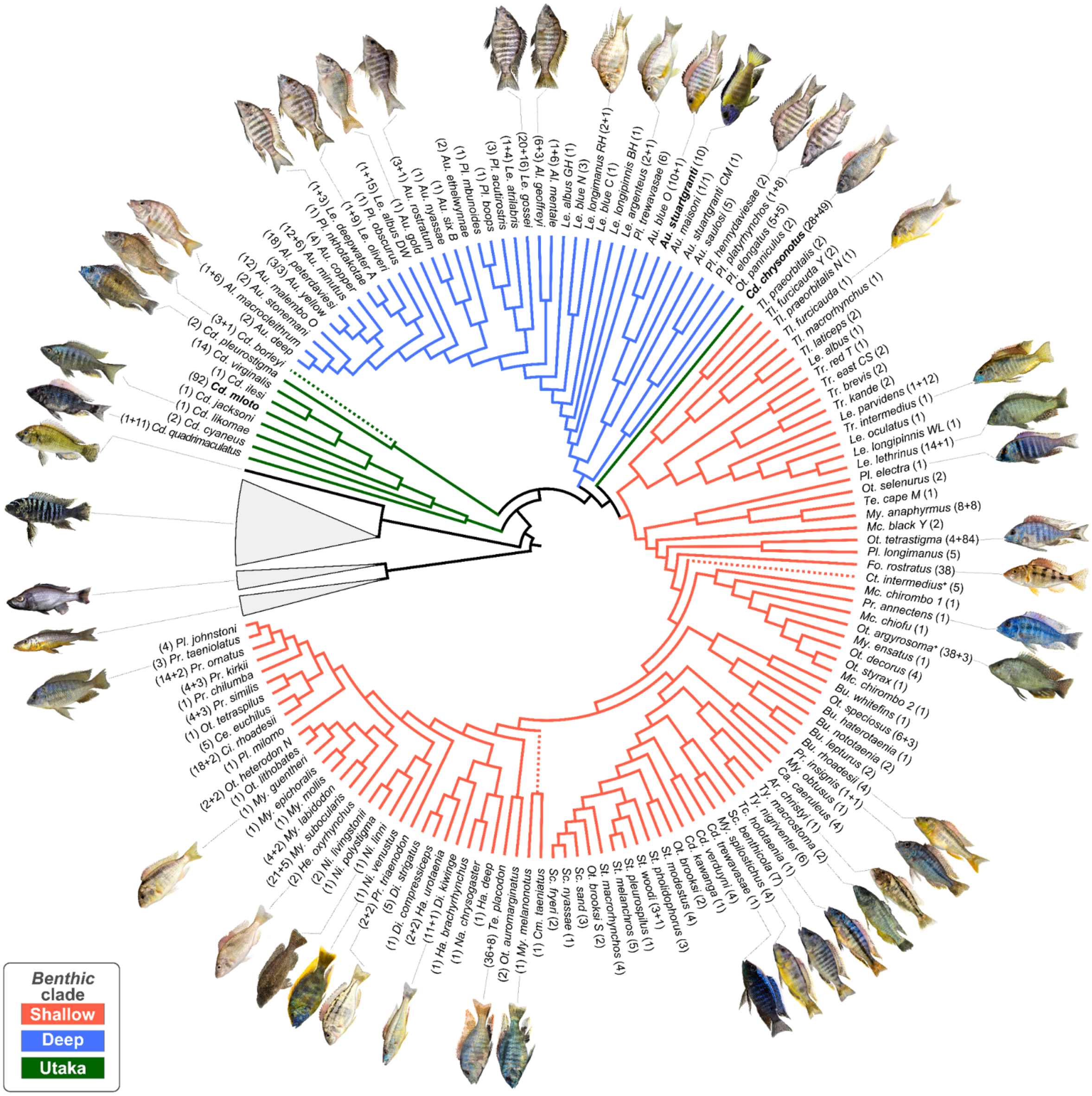

#### Fig. S1: Representative photos used in Fig. 1.

Simplified version of Fig. 1, with all information other than the phylogeny removed. The same photos of representatives of taxonomic groups are displayed, but arranged so that it is clear which species or clade they belong to. *Benthic* subclades are colour-coded in the tree (see legend). All photos were recorded by us, but the representative individuals shown in pictures are not necessarily the ones sequenced in this study. Species name abbreviations (tip labels) are decoded in table S3. We note that the positions of the three dashed branches (*Cd. borleyi*, *Ct. pictus* + *Ct. intermedius*, *My. melanonotus* + *Cm. taeniatus*) were unstable between runs with minor changes in filtration procedures due to short branches or nodes lacking statistical support. Their placement should be considered arbitrary. Our single sample of *Cyrtocara moori* grouped within *Ot. argyrosoma* clade, denoted by a plus sign.

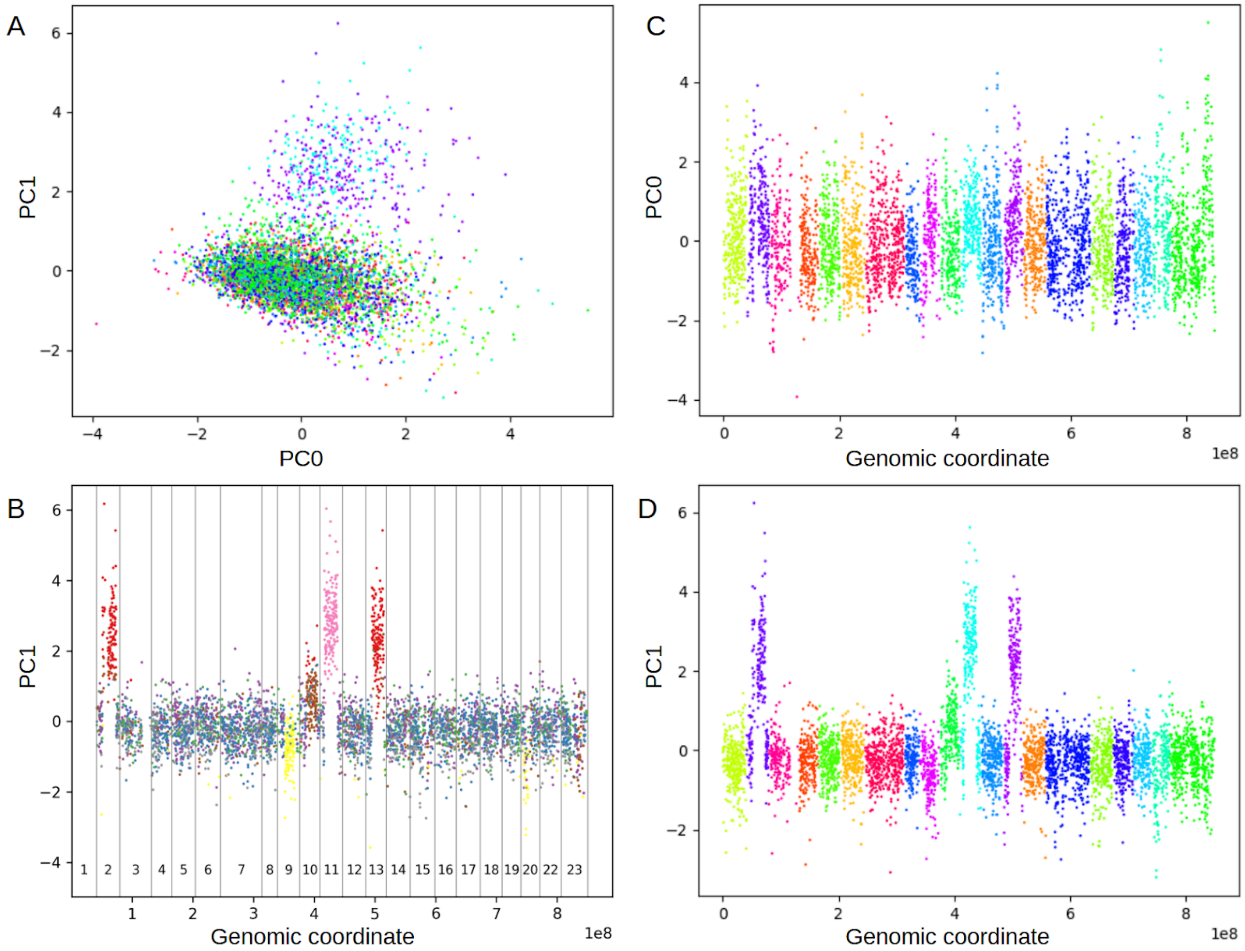

**Fig. S2: Clustering of distance matrices.**

(**A**)-(**E**) PCA analysis on flattened distance matrices. Each dot represents a 100 kbp genomic window. (**A**) First two principal components, colors by chromosome number. (**B**)-(**C**) Principal components vs linearized genomic coordinates, colors by chromosome number. (**B**) PC0. (**C**) PC. (**D**) The same as (**C**), but colors by cluster number for K-mean clustering with K=8, numbers above the "X" axis are chromosome numbers.

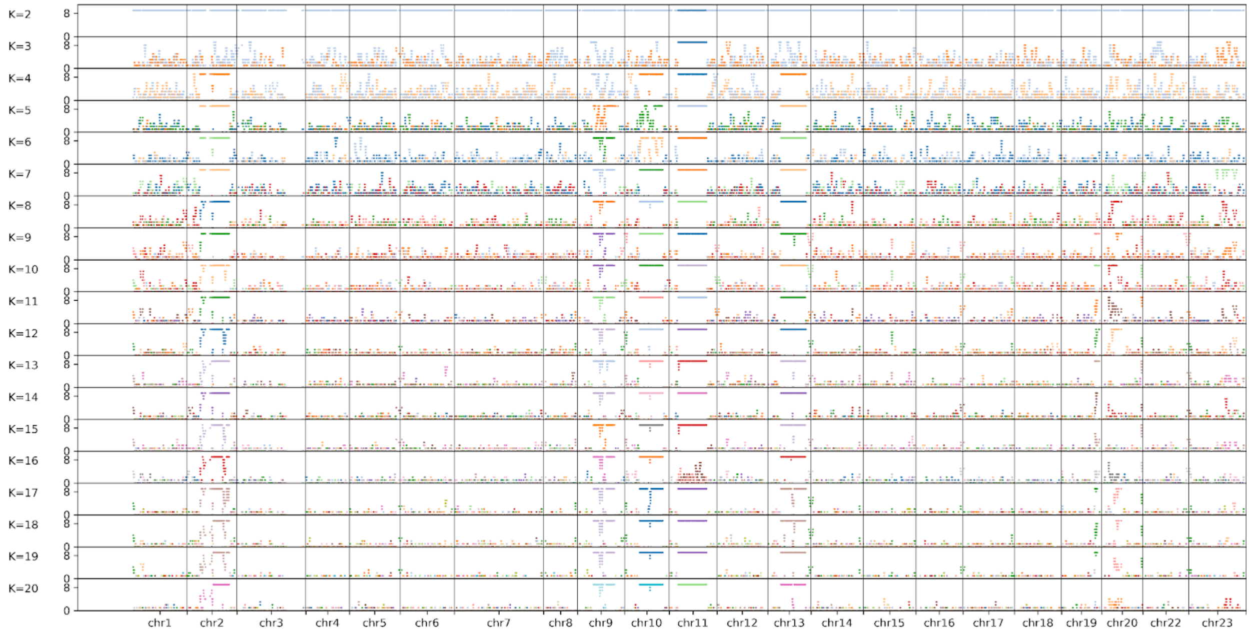

**Fig. S3: Contiguous runs of cluster identity.**

Spans of 100 kbp genomic windows which have the same cluster ID for apriori cluster count (K) from 2 to 20. X axis - linearized genomic coordinate, Y axis - greater of two values: count of 10 left or right neighbours with the same cluster ID as window under consideration. Color by cluster ID.

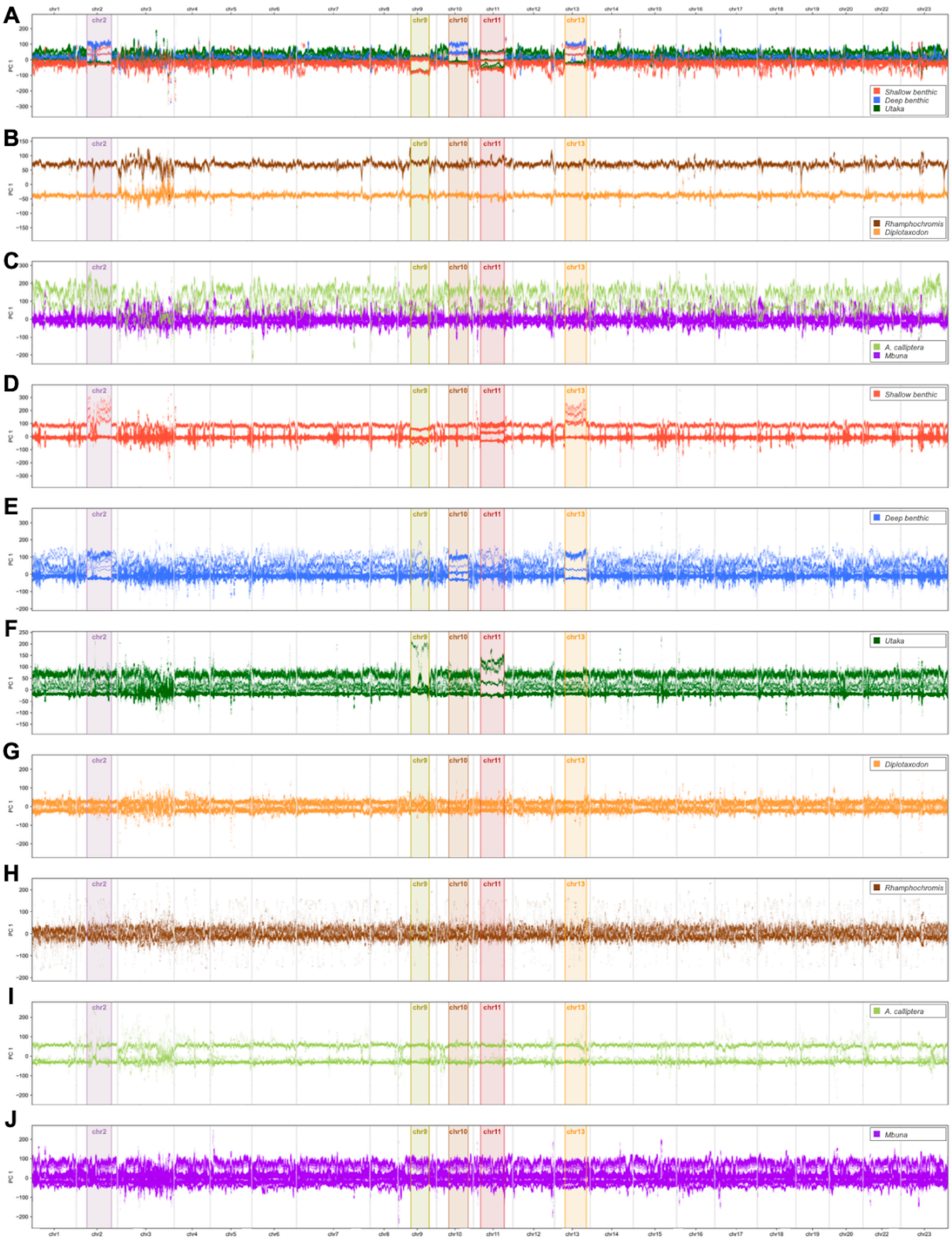

#### Fig. S4: Clade-wise windowed PC analysis.

First genetic principal component in overlapping 1 Mbp windows along chromosomes (see materials and methods) for combinations of clades (**A-C**) and for each clade separately (**D-J**). **A** is similar to the bottom panel of Fig. 2A, but uses all available *benthic* samples and does not omit outlier windows. Inversion regions are highlighted and colour-labelled using the same colours as the main text figures.

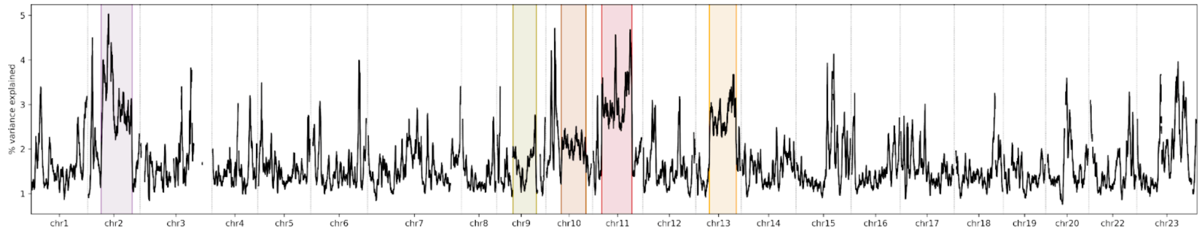

**Fig. S5: Variance explained by PC1.**

Variance explained by principal component 1 (PC 1) corresponding to the PC values shown in the bottom panel of Fig. 2A, and the same filters were applied for plotting (see materials and methods). Inversion regions are highlighted and colour-coded as in the main text figures.

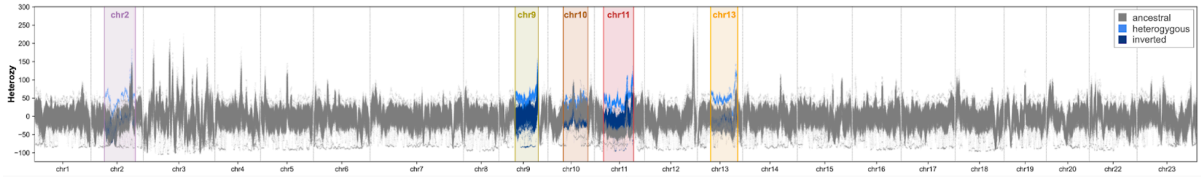

**Fig. S6: Windowed SNP heterozygosity.**

Number of heterozygous sites per sample in 1 Mbp windows along chromosomes (step size: 10 kbp) for species with ≥ 10 sequenced individuals. Y values are counts of heterozygous sites per window divided by 100,000 and normalised by the chromosome-wide (inversion-wide) average. Inversion regions are annotated and within inversions, samples are coloured by inversion state (see legend).

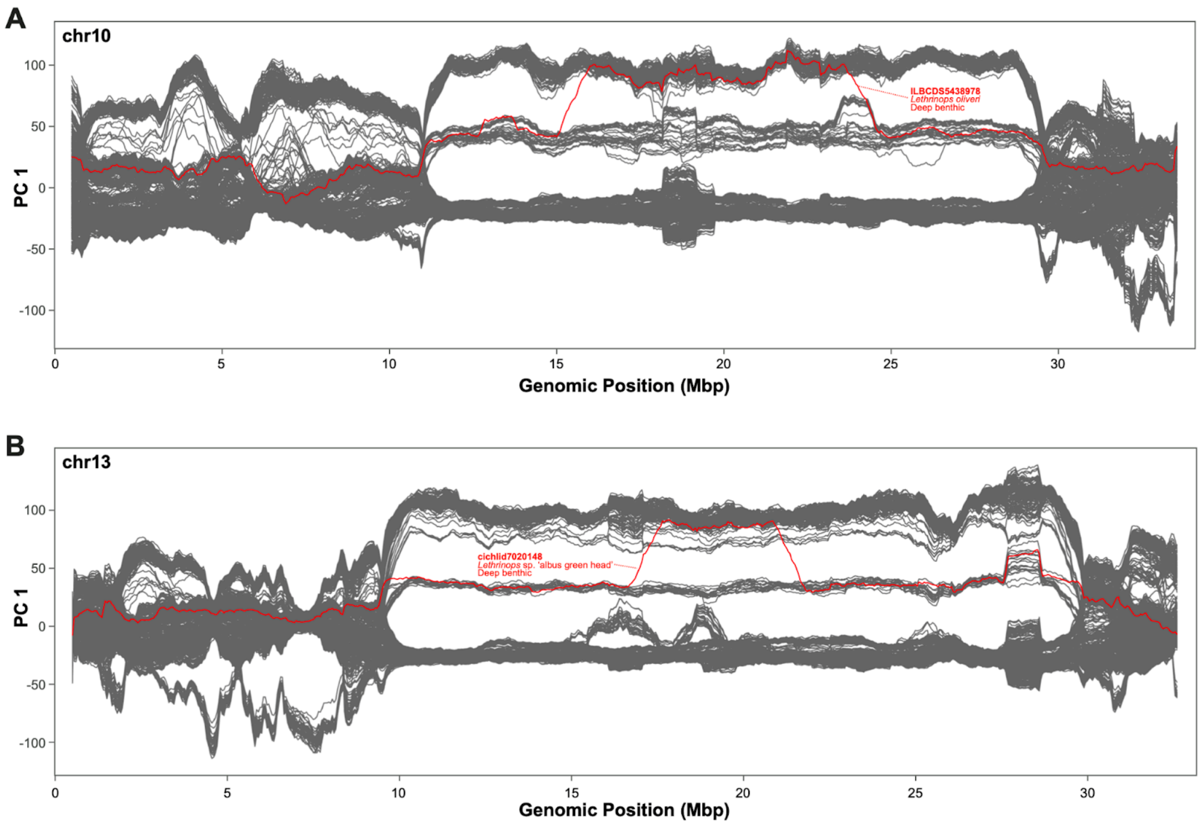

#### Fig. S7: Double recombination events in wild samples.

Across the entire dataset of wild samples, two double recombination events were observed in two different inversion regions. The respective samples are highlighted and annotated in red and plotted against the background of windowed PCA results of the remaining *benthic* samples (grey) for orientation. This figure re-plots data from Fig. 2A at higher resolution and using a different colour code to highlight double recombinants. Two *deep benthic* individuals from different *Lethrinops* species had double crossover events inside inversions: ILBCDS5438978 (*L. oliveri*) on chromosome 10 (**A**) and cichlid7020148 (*L.* sp. 'albus green head') on chromosome 13 (**B**).

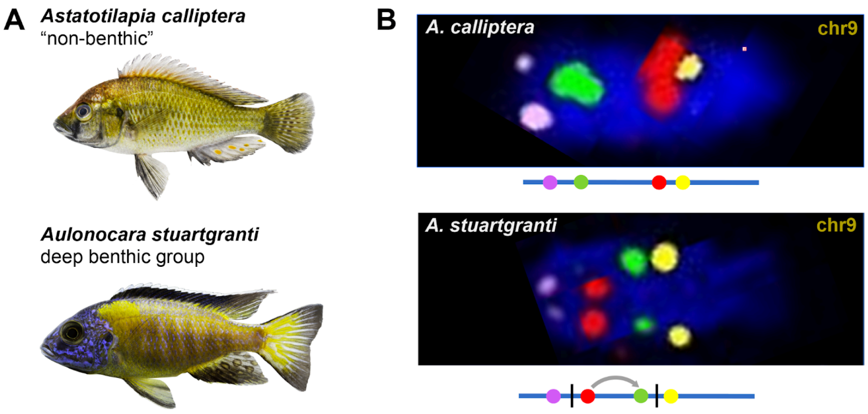

**Fig. S8: Fluorescence in situ hybridization results for chromosome 9.**

(**A**) Representative photographs of the species used. (**B**) Fluorescence in situ hybridisation (FISH) of markers on chromosome 9 left and right of the putative inversion breakpoints show the expected non-inverted orientation (upper panel) in *A. calliptera* while in *Au. stuartgranti* we see an inversion (lower panel).

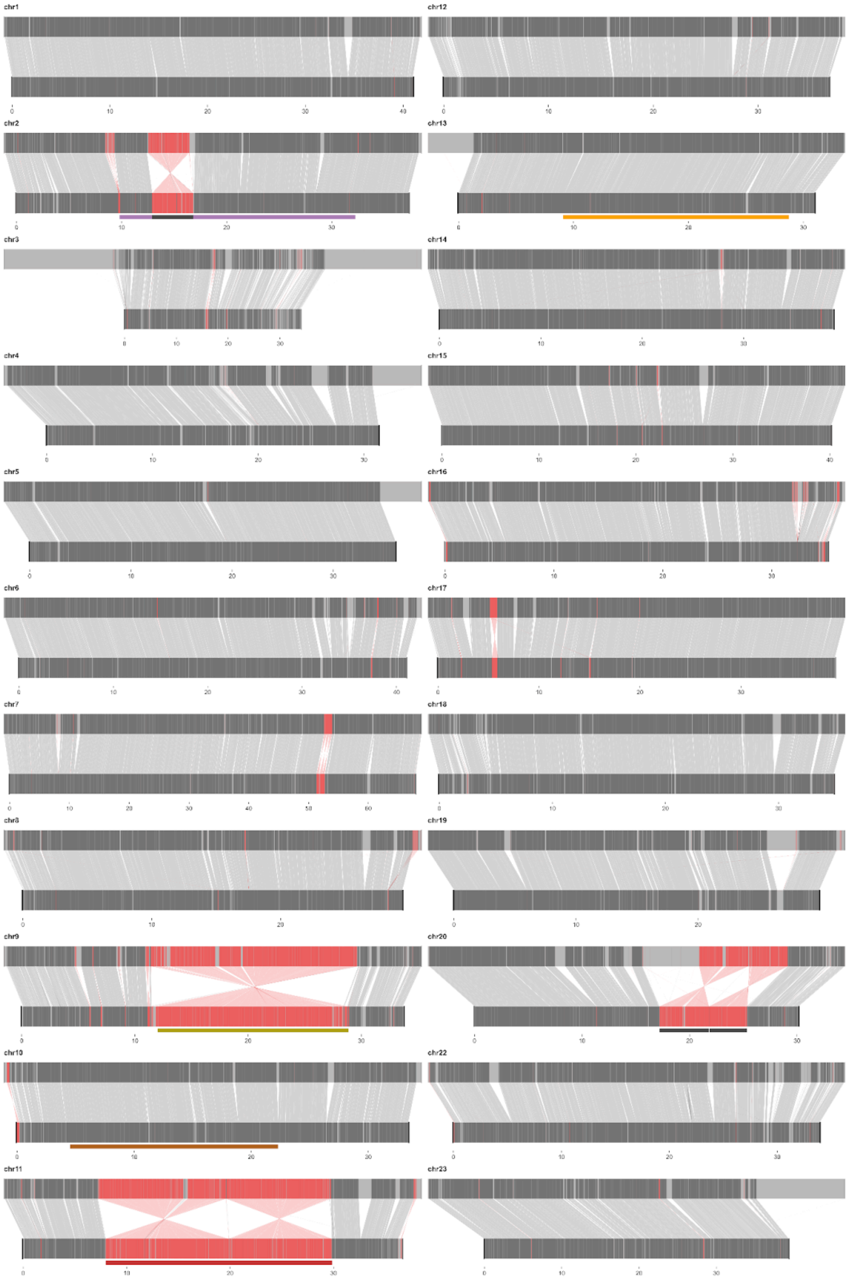

**Fig. S9: Structural rearrangements between *Astatotilapia calliptera* and *Aulonocara stuartgranti*.**

Whole genome alignment of *Au. stuartgranti* (top) to *A. calliptera* (bottom). Collinear alignment blocks are displayed in dark grey and grey lines connect them. Inverted blocks between reference and query are displayed in red, with red connector lines. Unaligned blocks (including ones that align to different chromosomes/scaffolds) are shown in light grey. The five focal inversion regions are shown as rectangles and colour-coded like in the main text figures. Locations of the three small inversions on chromosomes 2 and 20 (see text S1) indicated analogously as black rectangles.

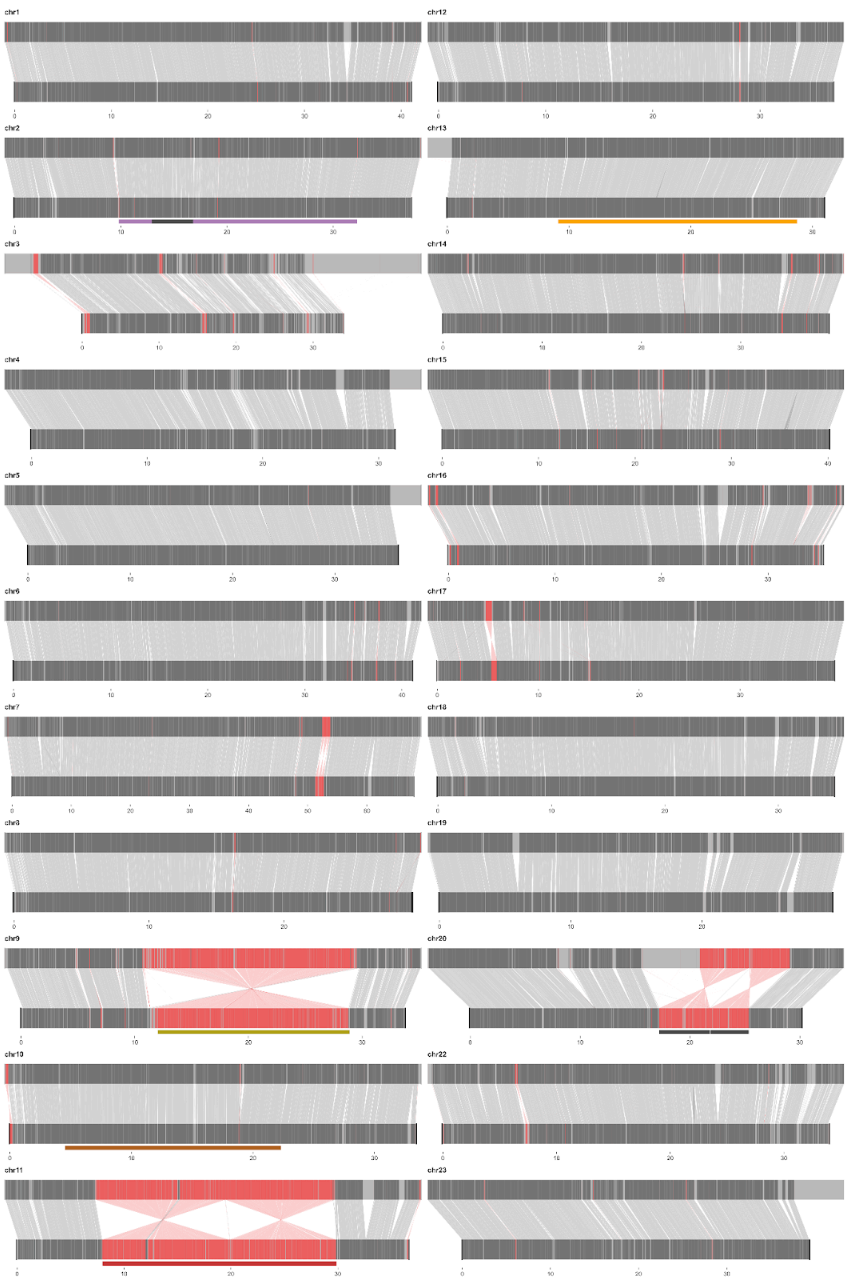

**Fig. S10: Structural rearrangements between *Astatotilapia calliptera* and *Diplotaxodon limnothrissa*.**

Whole genome alignment of *D. limnothrissa* (top) to *A. calliptera* (bottom). Collinear alignment blocks are displayed in dark grey and grey lines connect them. Inverted blocks between reference and query are displayed in red, with red connector lines. Unaligned blocks (including ones that align to different chromosomes/scaffolds) are shown in light grey. The five focal inversion regions are shown as rectangles and colour-coded like in the main text figures. Locations of the three small inversions on chromosomes 2 and 20 (text S1) indicated analogously as black rectangles.

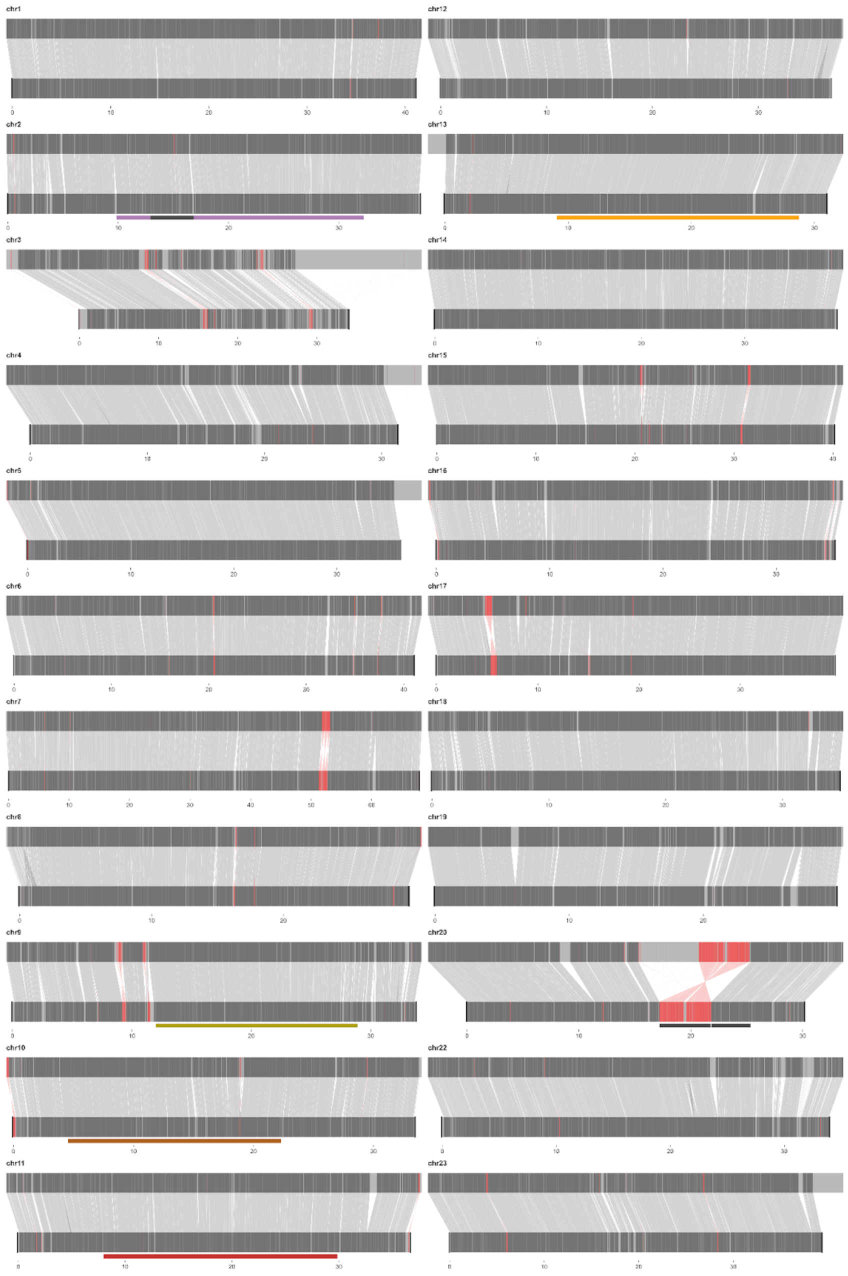

**Fig. S11: Structural rearrangements between *Astatotilapia calliptera* and *Rhamphochromis* sp. 'Chilingali'.**

Whole genome alignment of *R.* sp. 'Chilingali' (top) to *A. calliptera* (bottom). Collinear alignment blocks are displayed in dark grey and grey lines connect them. Inverted blocks between reference and query are displayed in red, with red connector lines. Unaligned blocks (including ones that align to different chromosomes/scaffolds) are shown in light grey. The five focal inversion regions are shown as rectangles and colour-coded like in the main text figures. Locations of the three small inversions on chromosomes 2 and 20 (text S1) indicated analogously as black rectangles.

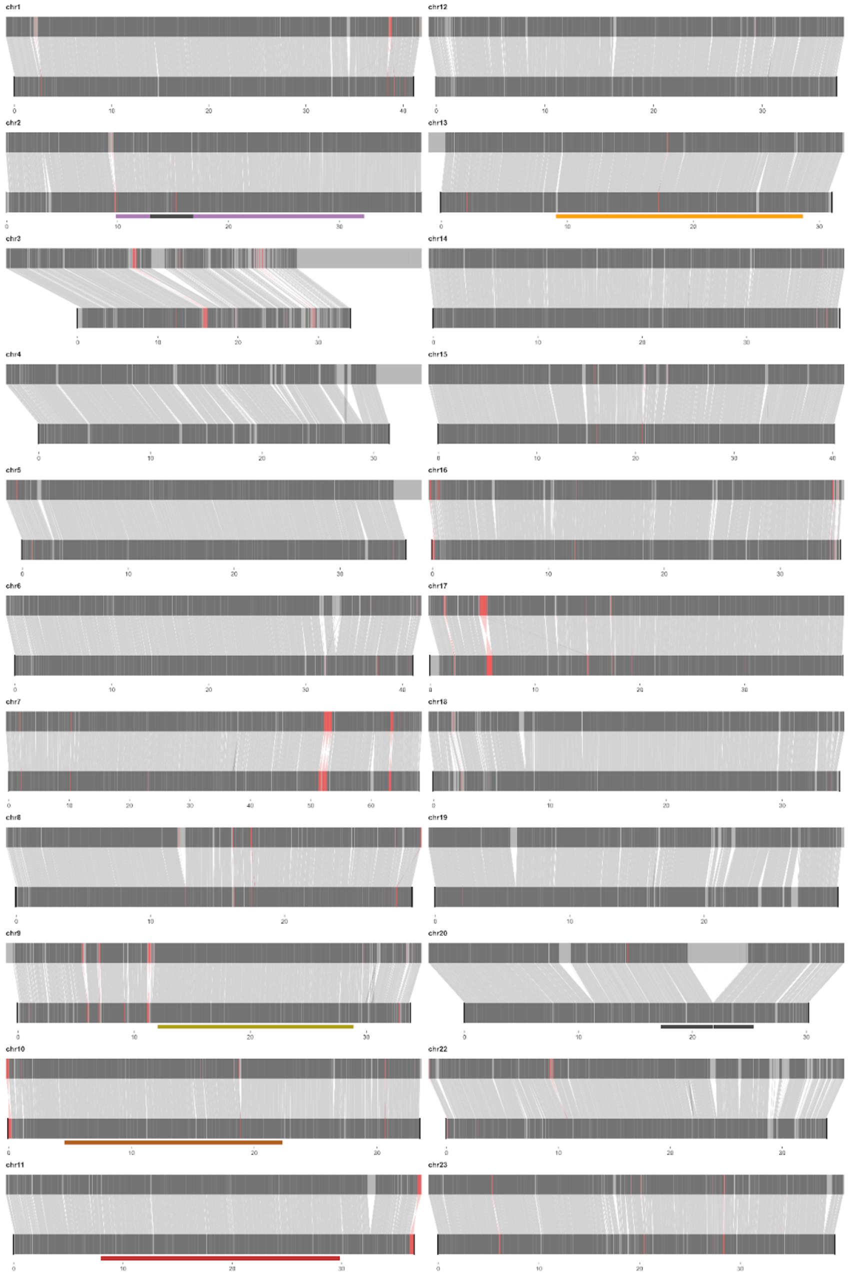

**Fig. S12: Structural rearrangements between *Astatotilapia calliptera* and *Tropheops* sp. ‘mauve’.**

Whole genome alignment of *T.* sp. ’mauve’ (top) to *A. calliptera* (bottom). Collinear alignment blocks are displayed in dark grey and grey lines connect them. Inverted blocks between reference and query are displayed in red, with red connector lines. Unaligned blocks (including ones that align to different chromosomes/scaffolds) are shown in light grey. The five focal inversion regions are shown as rectangles and colour-coded like in the main text figures. Locations of the three small inversions on chromosomes 2 and 20 (text S1) indicated analogously as black rectangles.

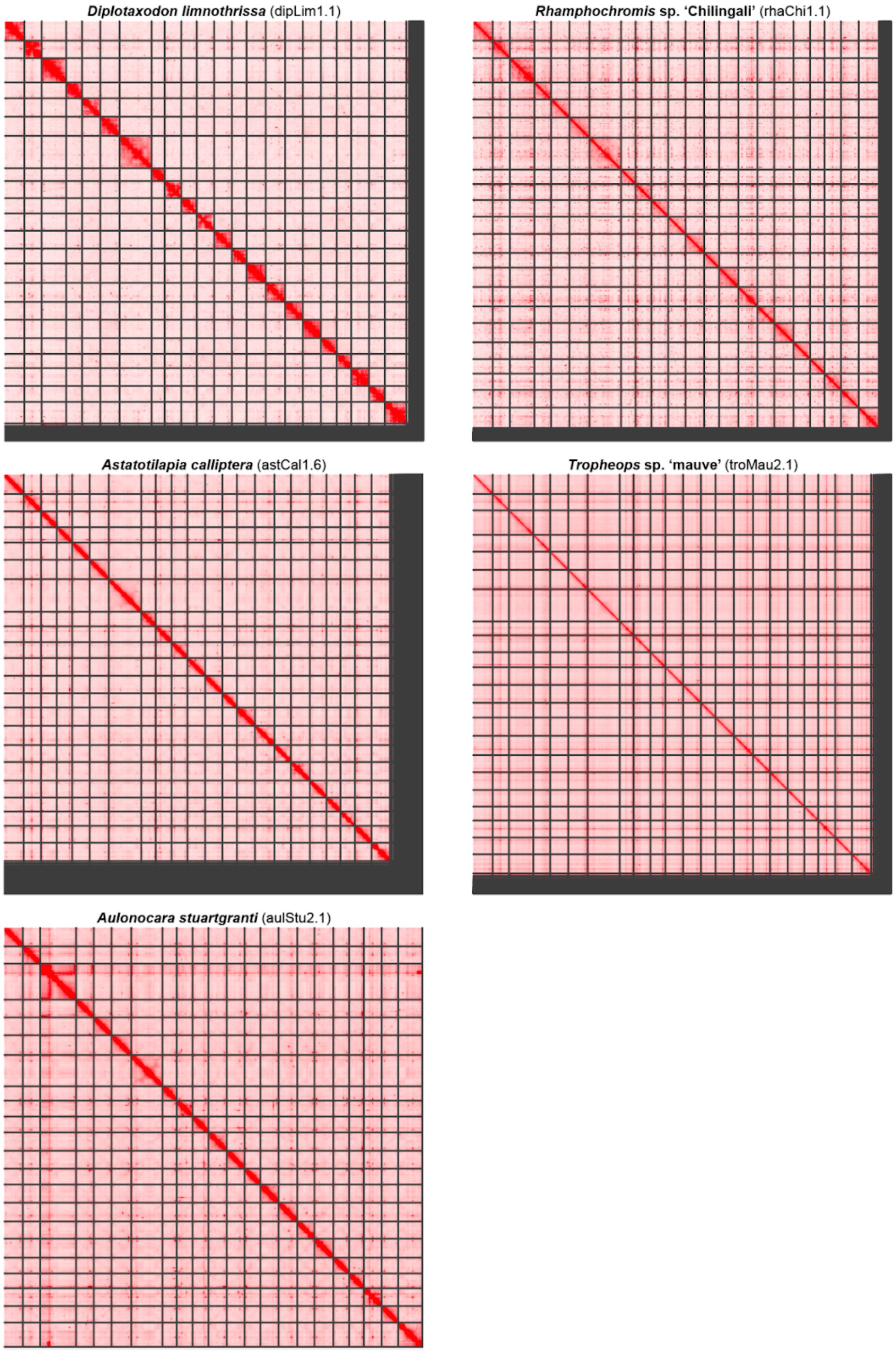

**Fig. S13: Hi-C maps of the chromosome-level genome assemblies.**

Hi-C maps of the five curated chromosome-level genome assemblies of Malawi cichlid clade representatives: *Diplotaxodon limnothrissa* (*Diplotaxodon*), *Rhamphochromis* sp. 'Chilingali' (*Rhamphochromis*), *Astatotilapia calliptera* (*A. calliptera*), *Tropheops* sp. 'mauve' (*mbuna*) and *Aulonocara stuartgranti* (*benthic*). See materials and methods and table S4 for more details on each assembly.

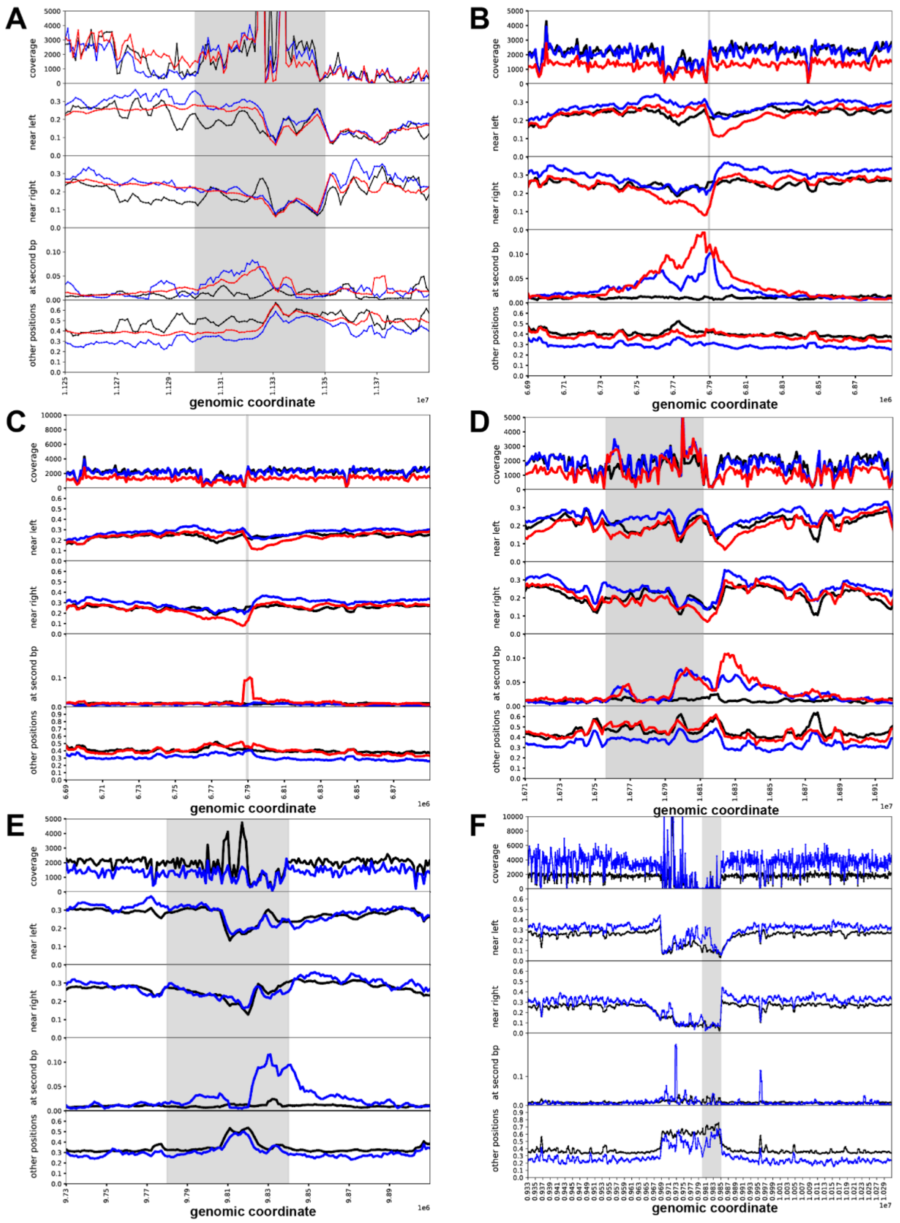

#### Fig. S14: Proportion of tags shared between breakpoints

The share of all tags in 1 kbp windows, which have at least one read with the same tag mapping 'near left'/'near right' (in the 100 kbp region next to the window under consideration, upstream and downstream respectively), 'at second bp' (in the 1 Mbp region centred around putative second inversion breakpoint) and 'other positions' (more than 100 kbp apart from the window under consideration and not in 'at the second bp' category). (**A**) chromosome 9. (**B**) chromosome 11, breakpoints 1 and 2. (**C**) chromosome 11, breakpoints 1 and 3; (**D**) chromosome 11, breakpoints 2 and 3. (**E**) chromosome 13. (**F**) chromosome 2. Rolling window smoothing of 5 data points. All metrics averaged over individuals with the same inversion state (black: ancestral, blue heterozygous, red: both haplotypes inverted). Grey background: putative breakpoint position derived from SNP PCA.

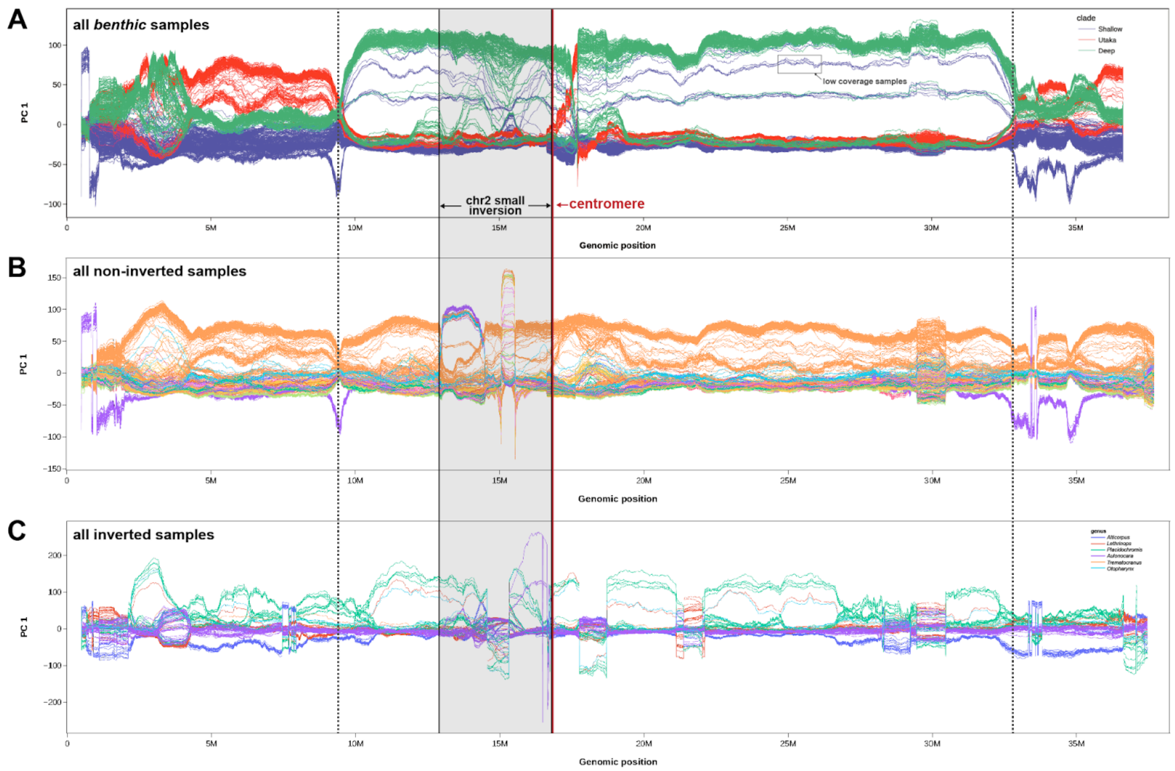

#### Fig. S15: Nested inversion on chromosome 2.

Windowed genetic PC analysis of chromosome 2 for three sets of samples. Located within the large inversion on chromosome 2 is a second smaller inversion, which was identified in pairwise whole genome alignments (text S1). In the panels (**A**)-(**C**), breakpoint locations of the large chromosome 2 inversion region are shown as dashed vertical lines. The centromere is shown as a red vertical line and the location of the small nested inversion (which is directly adjacent to the centromere) is highlighted in grey, while its breakpoint locations are shown as solid black vertical lines. (**A**) Windowed PC analysis of chromosome 2 featuring all *benthic* samples. Within the nested inversion region the usual separation of inversion genotypes into three clusters (top: homozygous inverted, centre; heterozygous, bottom: homozygous non-inverted) is less clear-cut compared to the remainder of the inversion, and compared to the other inversions (Fig. 2A, bottom panel). [Note that a small set of samples homozygous for the inversion has notably lower PC 1 values compared to the remaining samples of the same genotype, which is due to low sequencing coverage and the consequential increased rate of SNP genotyping errors]. (**B**) Windowed PC analysis for the subset of homozygous non-inverted samples from (**A**), coloured by genus to aid visual interpretation (names not shown due to the large number of genera). Inside the nested inversion region, a set of samples groups into distinct clusters in the first third of the inversion region. (**C**) Windowed PC analysis for the subset of homozygous inverted samples from (**A**), coloured by genus. A different set of samples forms distinct clusters in the last third of the inversion region. These observed instances of aberrant clusters in subsections of the small nested inversion suggest strong genetic structure in that region, possibly distributed across several principal components and therefore not always represented to the same extent in the displayed PC 1. It is conceivable that this region (and possibly another region of similar size to the right of the centromere) is structurally highly variable among *benthic* species. Genome sequences from additional samples that span this region are necessary to investigate this hypothesis more comprehensively.

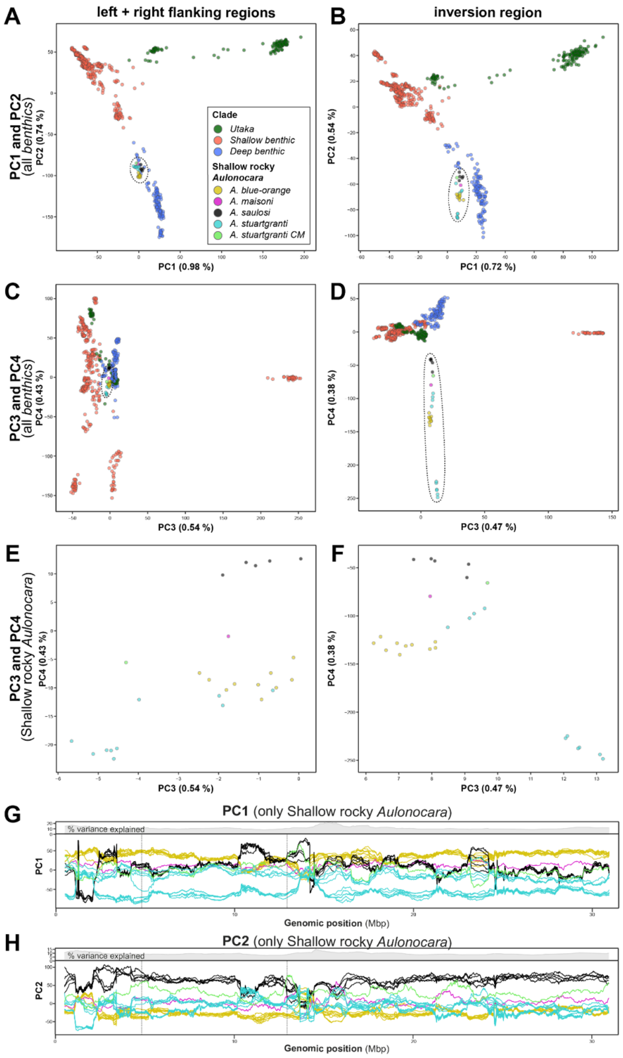

#### Fig. S16: PCA of chromosome 20.

We identified a double inversion between Diplotaxodon and benthic clades at chr20:4788189-12943513 through whole genome alignments (see materials and methods). However, the region went undetected in our scan for polymorphism inversion patterns which relies on local changes in genetic structure. To rule out the possibility of a yet undetected inversion polymorphism reflected in higher order principal components, we examined PCs 1 to 4 for the chromosome 20 inversion region separately. Panels are colour-coded by clade with members of shallow rocky Aulonocara (who belong to the deep benthic clade) highlighted separately. (A) PC1 and PC2 of benthic samples in the non-inverted flanking regions. (B) PC1 and PC2 of benthic samples in the inversion region. (C) PC3 and PC4 of benthic samples in the non-inverted flanking regions. (D) PC3 and PC4 of benthic samples in the inversion region. (E) shallow rocky Aulonocara samples from (C) plotted separately for increased resolution. (F) shallow rocky Aulonocara samples from (D) plotted separately for increased resolution. (G) PC1 of a windowed PCA of shallow rocky Aulonocara along chromosome 20. The left and right inversion breakpoints are indicated with dotted vertical lines. (H) PC2 of a windowed PCA of shallow rocky Aulonocara along chromosome 20. The left and right inversion breakpoints are indicated with dotted vertical lines.

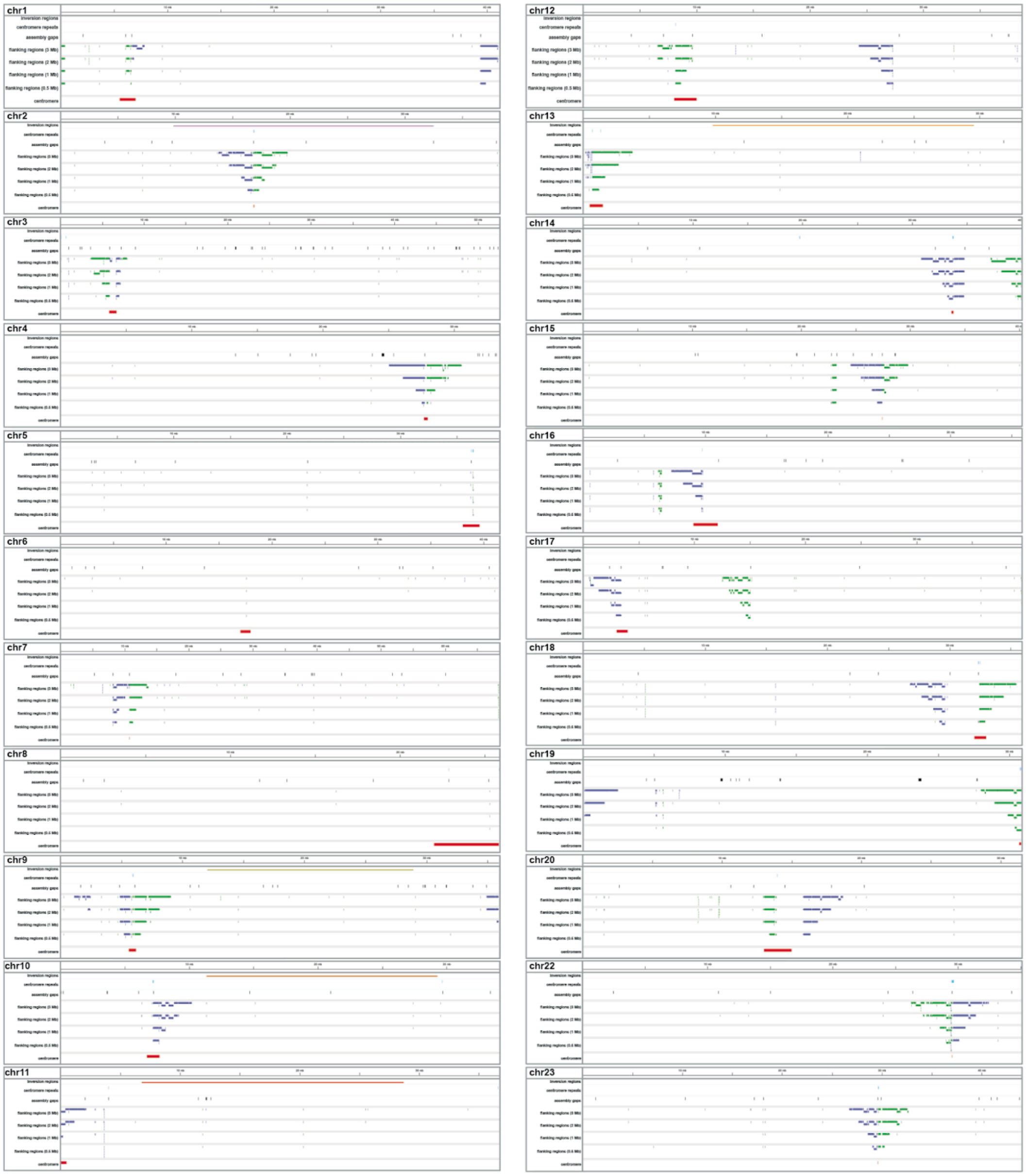

**Fig. S17: Centromere identification in the reference genome.**

Since centromeres are mostly unassembled in the reference fAstCal1.2 assembly we instead mapped centromeres in the higher quality *D. limnothrissa* assembly and then mapped the flanking regions back to fAstCal1.2 (see materials and methods). For each chromosome, different types of evidence that were used to locate centromeres are displayed as IGV (*123*) tracks: (a) assembled centromeric repeats (if present), (b) assembly gaps in the fAstCal1.2 assembly, (c) the aligned *Diplotaxodon* centromeric flanking regions (500 kb, 1 Mb, 2 Mb, 3 Mb). Additionally, the focal inversion regions are displayed and the final approximated centromere locations are shown. Note that this analysis used a slightly different version of the *D. limnothrissa* assembly (pre curation) than the one made available.

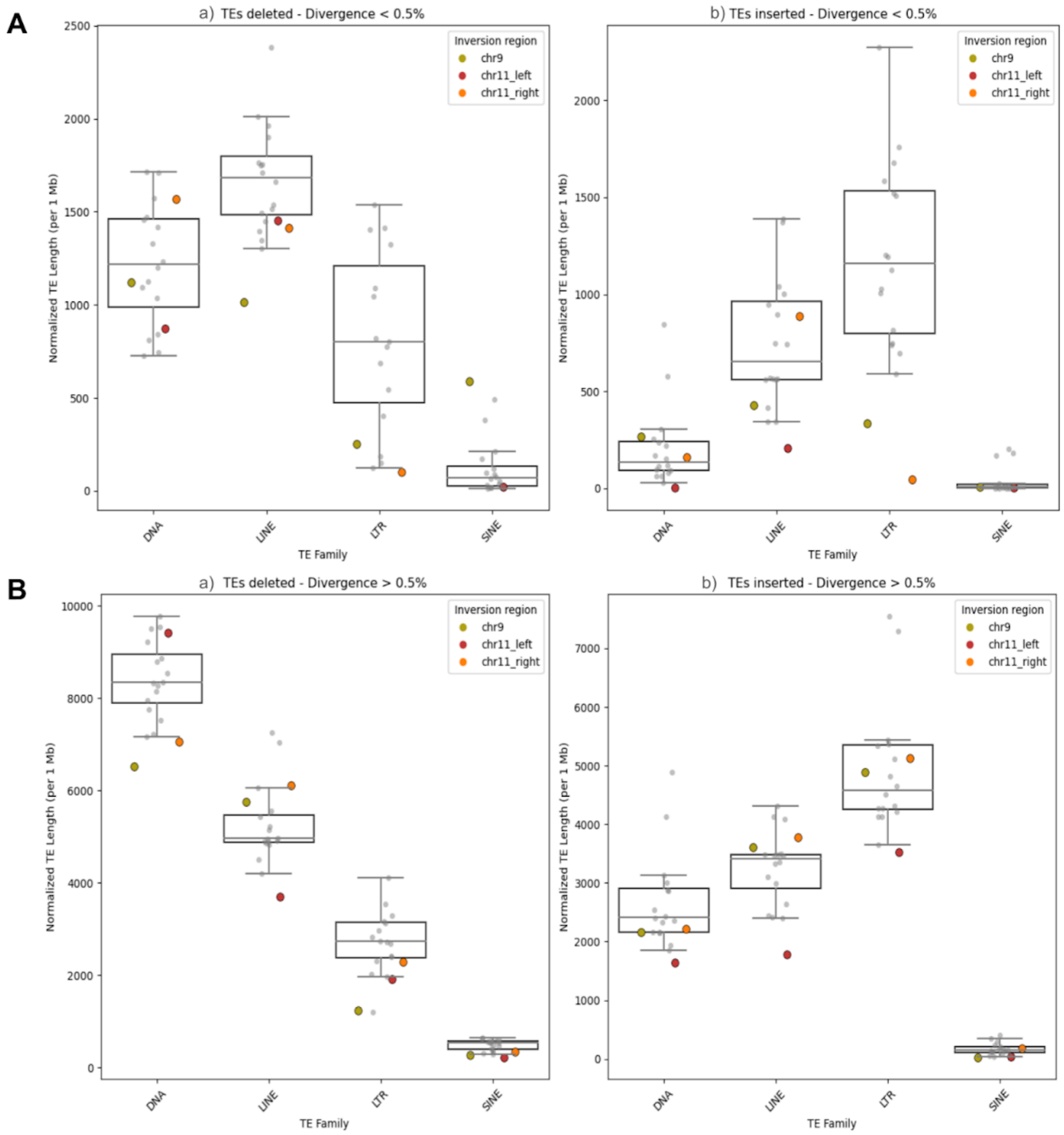

**Fig. S18: Differences in TE length between inverted and non-inverted haplotypes.**

We compared the length of transposable element (TE) insertions in a whole genome alignment of *Au. stuartgranti* to *A. calliptera*. Insertions correspond to *Au. stuartgranti*-specific TEs, while deletions correspond to *A. calliptera*-specific TEs. Box plots show the amount of sequence classified as transposable elements (per Mbp) in the genome. Each grey marker represents the mean TE length for one of the non-inverted chromosomes, while coloured markers correspond to inversion regions. (**A**) TEs with divergence below 0.5% with deletions (a) and insertions (b) shown separately. (**B**) TEs with divergence above 0.5% with deletions (a) and insertions (b) shown separately. See materials and methods on details on calculation and choice of the divergence threshold.

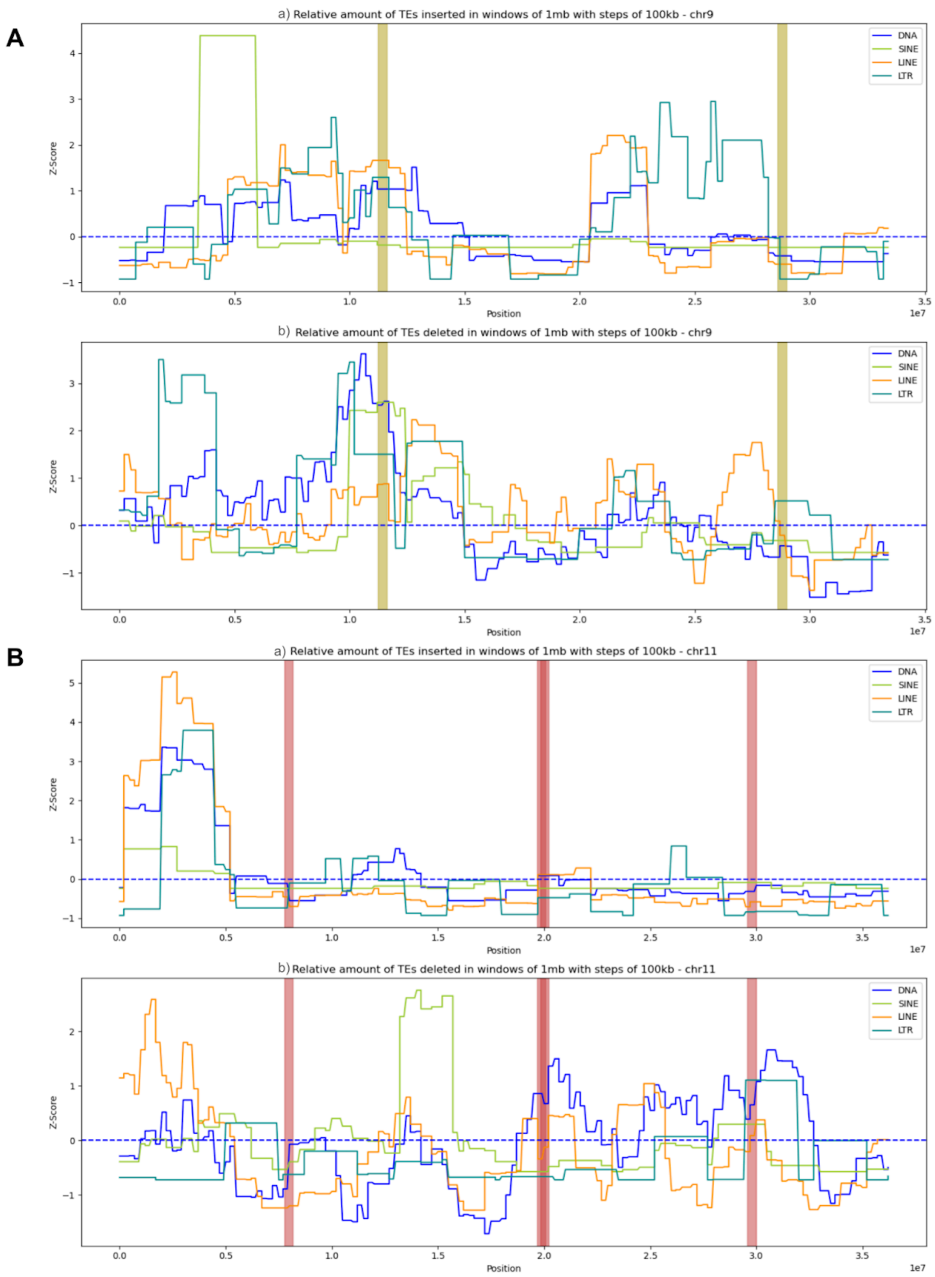

**Fig. S19: Transposable element content along inversion chromosomes.**

We compared the length of transposable element (TE) insertions in a whole genome alignment of *Au. stuartgranti* to *A. calliptera*. Insertions correspond to *Au. stuartgranti*-specific TEs, while deletions correspond to *A. calliptera*-specific TEs. Z-scores were calculated based on genome wide mean (only for non-inverted chromosomes) of TE lengths for inverted chromosomes in windows of 1 Mbp with a step size of 100 kbp for (**A**) chromosome 9 (a) inserted sequences (b) deleted sequences and (**B**) chromosome 11 (a) inserted sequences and (b) deleted sequences for *Aulonocara stuartgranti*. The highlighted regions show the +-200 kbp region around inversion breakpoints  (ochre: chromosome 9 inversion, red: chromosome 11 inversion).

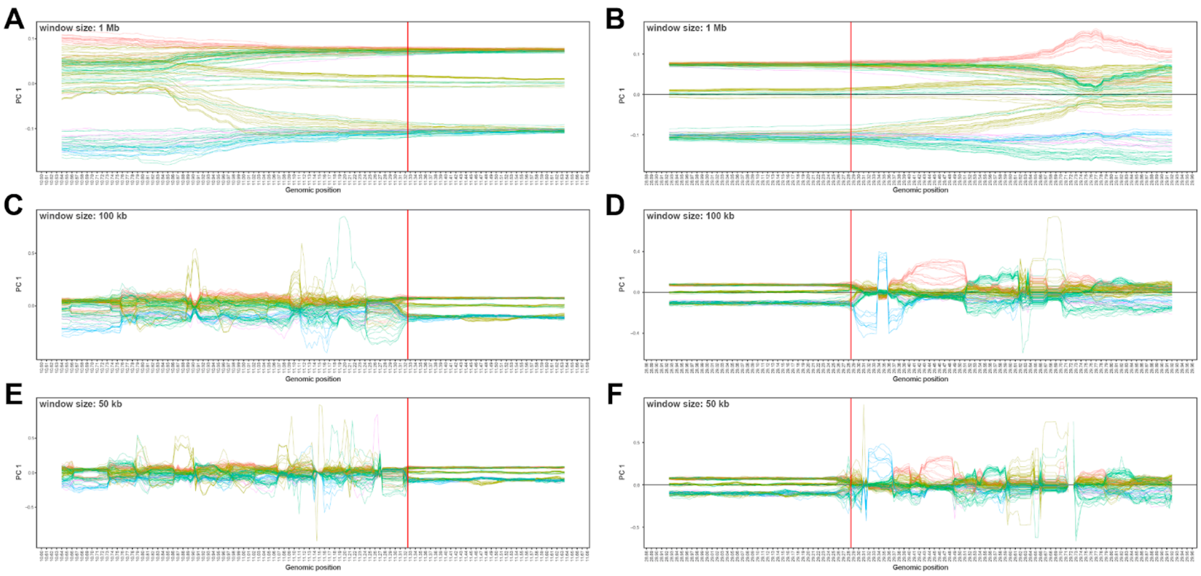

**Fig. S20 Inversion breakpoint estimation.**

Inversion breakpoint locations in the reference genome were approximated by conducting local high resolution windowed PCAs (step size: 5 kbp, window sizes 1 Mbp, 100 kbp, 50 kbp, see materials and methods). The above figure panels illustrate this approach for one inversion chromosome; the left breakpoint in panels (**A**), (**C**) and (**E**) and for the right breakpoint in panels (**B**), (**D**) and (**F**). The final breakpoint estimates are indicated as red vertical lines.

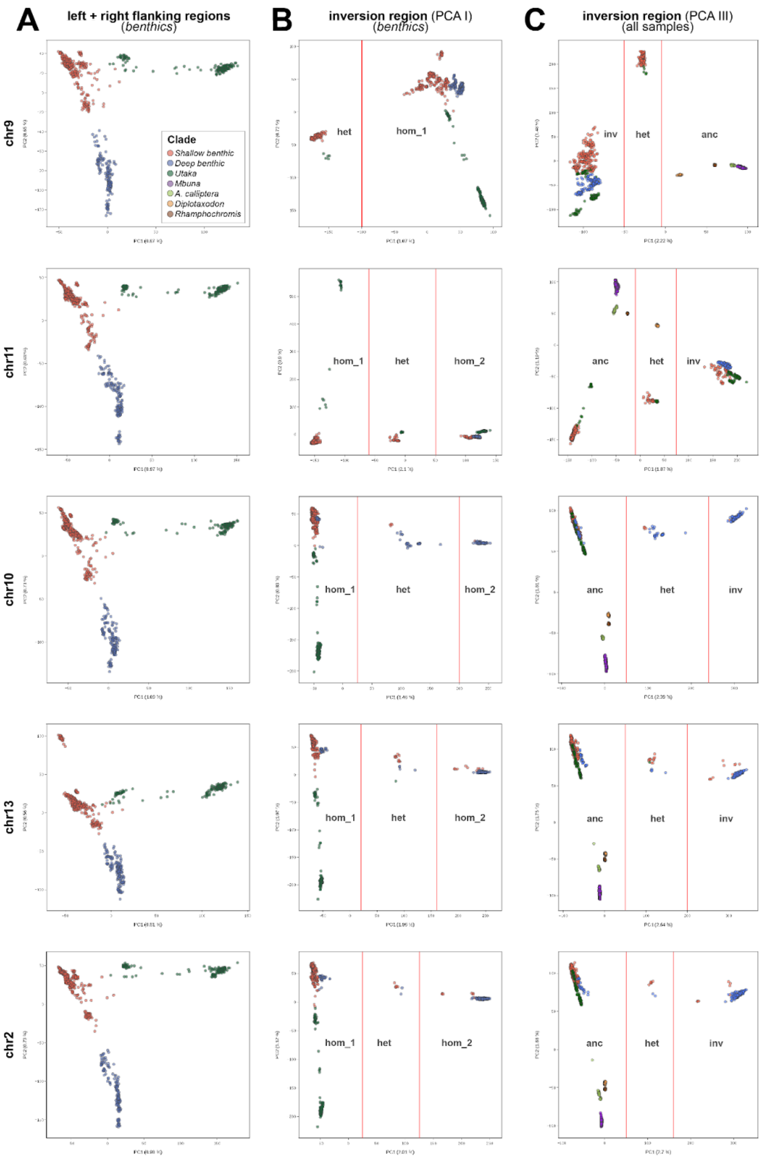

#### Fig. S21: PCA-based genotyping of inversion state.

A multi-step PCA-based approach was used to genotype all 1,375 sequenced samples' inversion states (see materials and methods). Shown for each inversion chromosome is: (**A**) the first two principal components of *benthic* samples outside the inversion region (flanking regions). (**B**) the first two principal components of *benthic* samples inside the inversion region (referred to as 'PCA I' in the methods). (**C**) the first two principal components of all 1,375 included samples inside the inversion region, but using only SNPs that were variable in a balanced set of homozygous samples from PCA I (referred to as 'PCA III' in the methods). In (**B**) (PCA I) and (**C**) (PCA III), PC1 values used as inversion genotype assignment thresholds are shown as vertical red lines and genotypes are annotated. Final genotypes (ancestral, heterozygous, inverted) were inferred from the location of the (non-benthic) outgroup samples on PC1 of PCA III. The distant genotype cluster relative to the outgroup samples was assumed to represent the homozygous inverted state, which was later confirmed in genome alignments with outgroup species (see materials and methods).

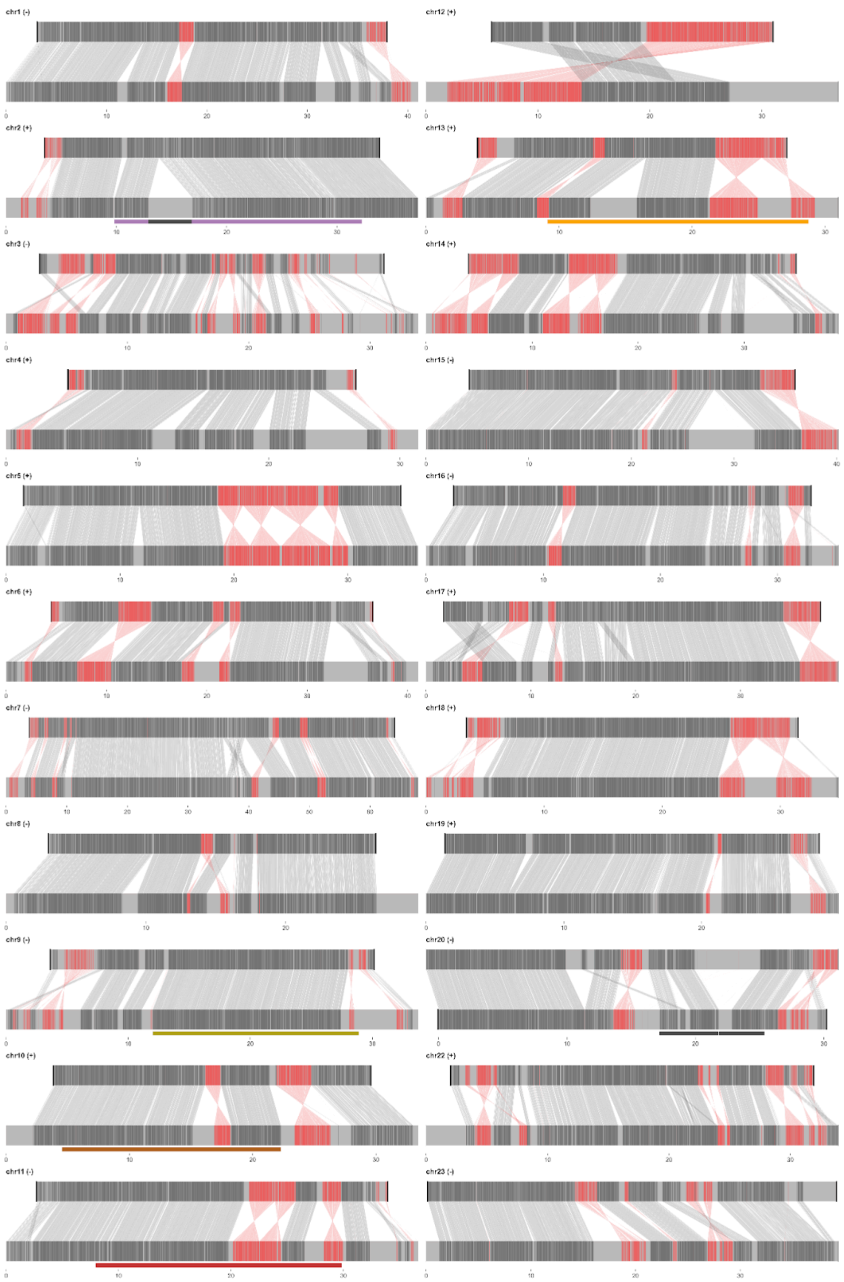

**Fig. S22: Structural rearrangements between *Astatotilapia calliptera* and *Pundamilia nyererei*.**

Whole genome alignment of *P. nyererei* (top) to *A. calliptera* (bottom). Collinear alignment blocks are displayed in dark grey and grey lines connect them. Inverted blocks between reference and query are displayed in red, with red connector lines. Unaligned blocks (including ones that align to different chromosomes/scaffolds) are shown in light grey. The five focal inversion regions are shown as rectangles and colour-coded like in the main text figures. Locations of the three small inversions on chromosomes 2 and 20 (text S1) indicated analogously as black rectangles.

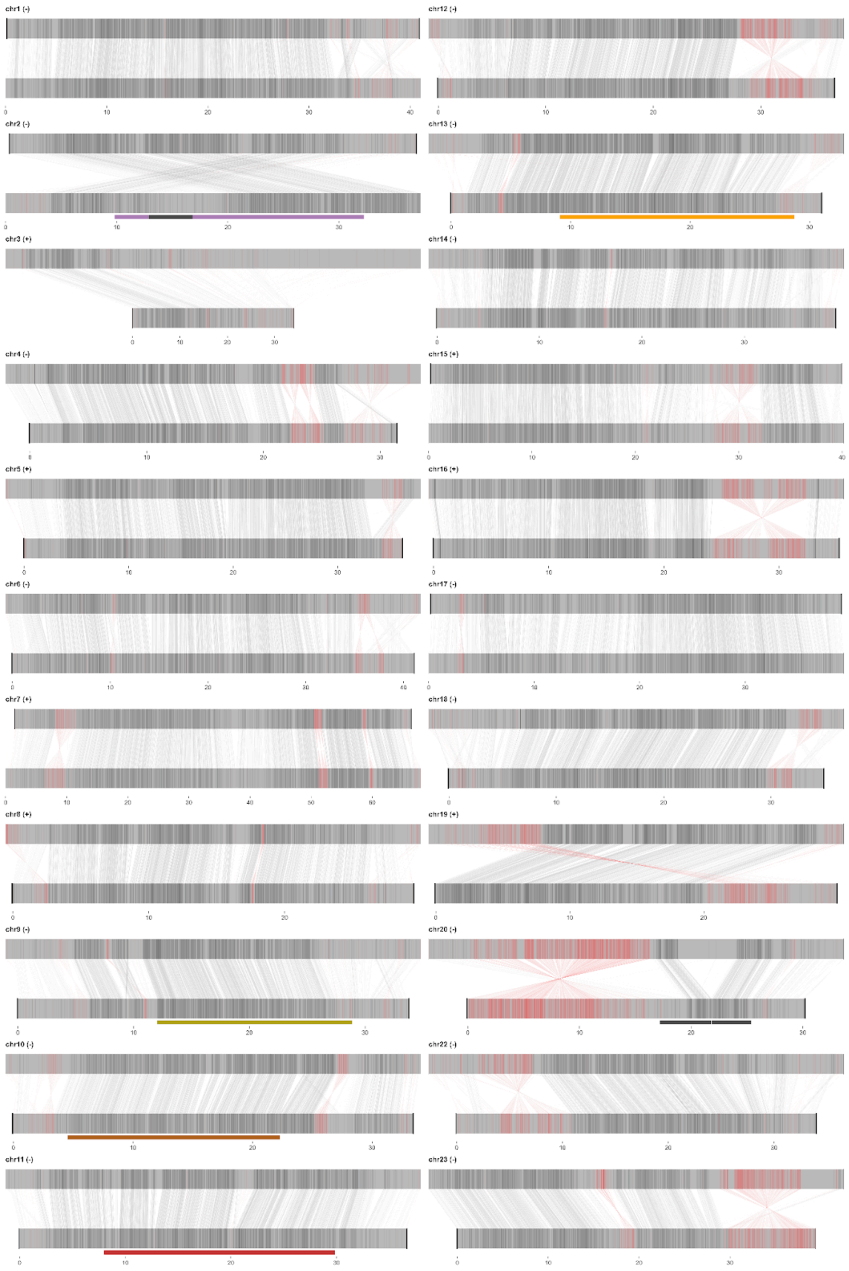

**Fig. S23: Structural rearrangements between *Astatotilapia calliptera* and *Oreochromis niloticus*.**

Whole genome alignment of *O. niloticus* (top) to *A. calliptera* (bottom). Collinear alignment blocks are displayed in dark grey and grey lines connect them. Inverted blocks between reference and query are displayed in red, with red connector lines. Unaligned blocks (including ones that align to different chromosomes/scaffolds) are shown in light grey. The five focal inversion regions are shown as rectangles and colour-coded like in the main text figures. Locations of the three small inversions on chromosomes 2 and 20 (text S1) indicated analogously as black rectangles.

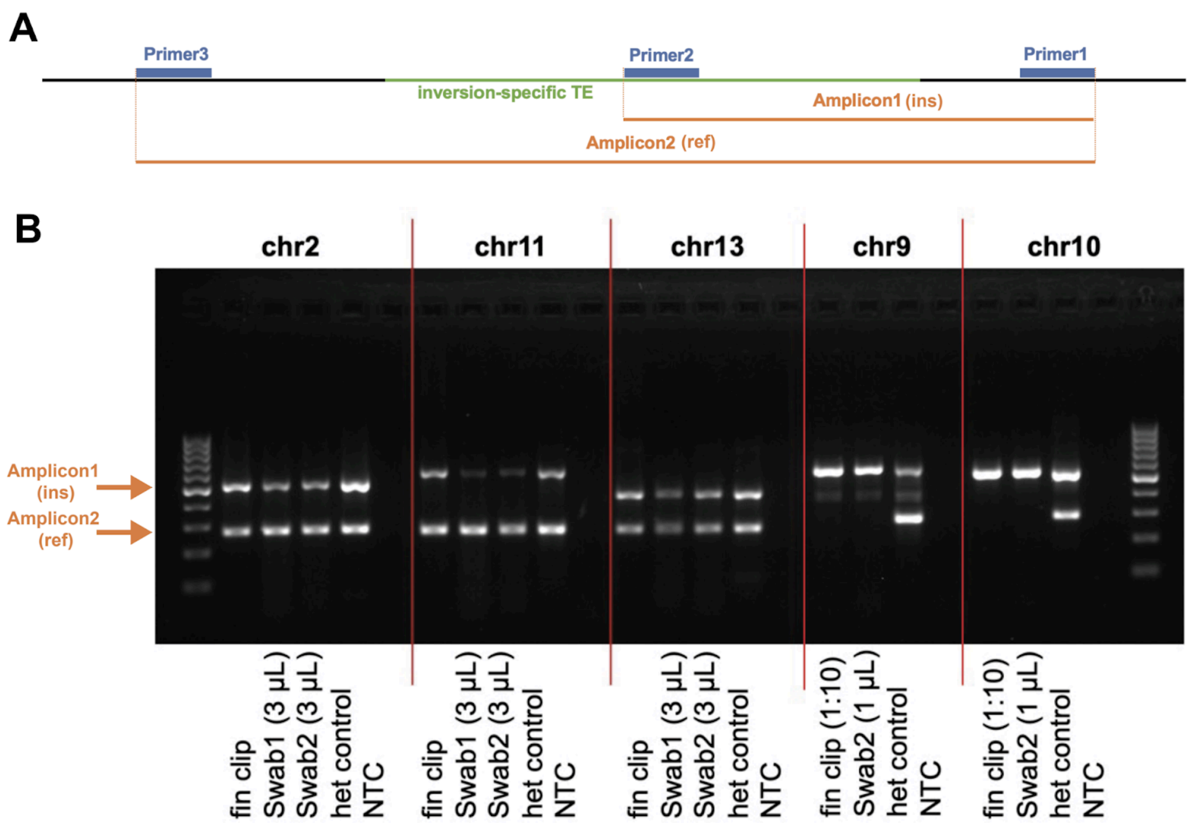

#### Fig. S24: PCR-based inversion genotyping.

(**A**) Inversion-correlated transposable element (TE) insertions were used to design PCR assays for inversion detection. A pair of primers 1 and 2 produce short Amplicon 1 (~250bp), specific for inverted state, a pair of primers 1 and 3 produce long Amplicon 2 (~500bp), specific for non-inverted state. Specific primers were designed for detection of inversions on chromosomes 2, 9, 10, 11,13. (**B**) Example gel electrophoresis illustrating PCR-typing method. Amplified fragments demonstrate clear bands at 250bp and 500bp for all 5 PCR assays (NTC: negative control). DNA samples extracted from fin clips and non-lethal skin swabs produced equally detectable bands.

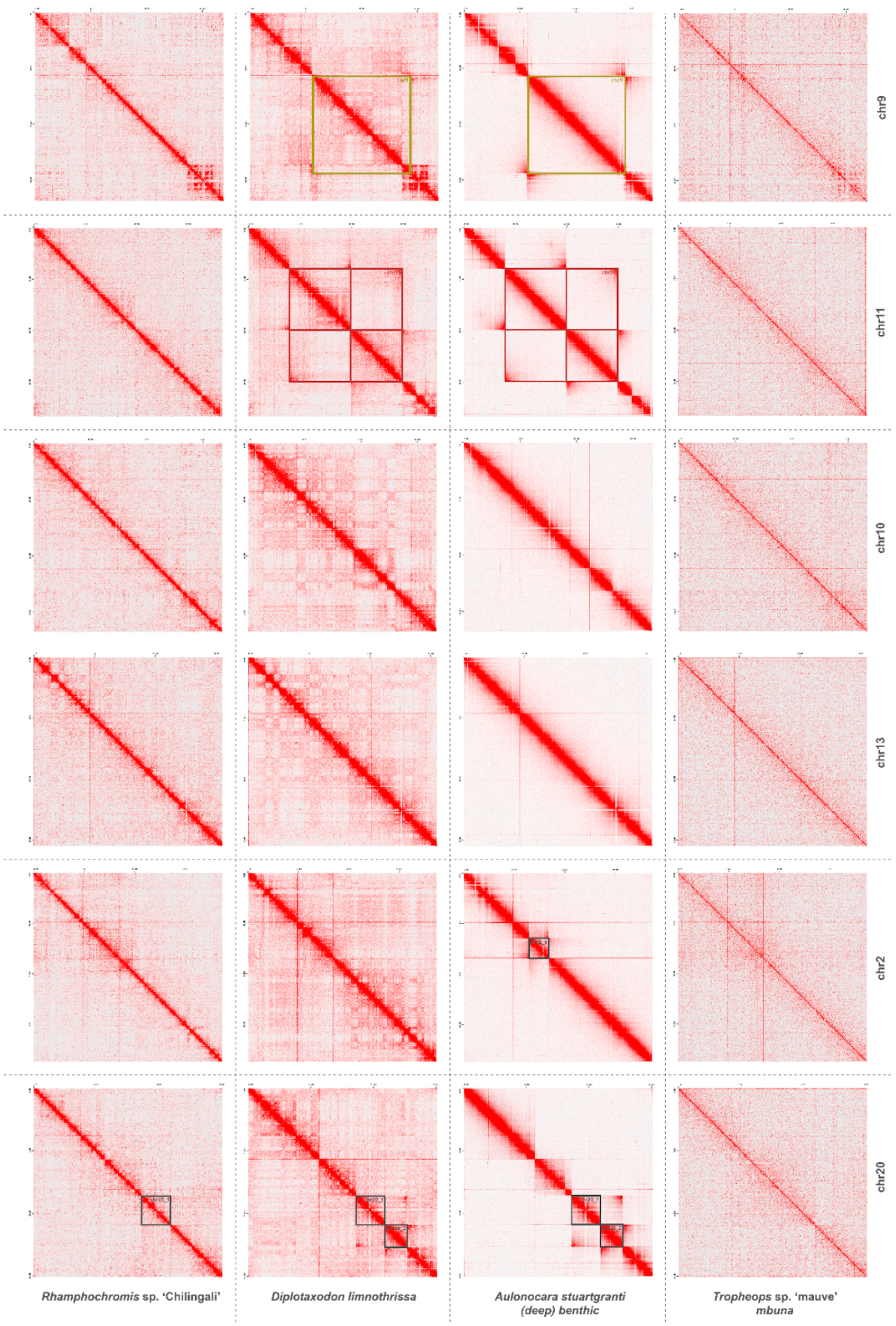

#### Fig. S25: Inversions are reflected in Hi-C datasets.

Hi-C reads of a *Rhamphochromis*, a *Diplotaxodon*, a (*deep*) *benthic* and a *mbuna* representative mapped against the *A. callipera* (fAstCal1.6) assembly. Locations of focal inversions on chromosomes 9 and 11 (colour-coded as in Fig. 1) are shown, as well as three additionally identified smaller inversions on chromosomes 2 and 20 (dark grey).

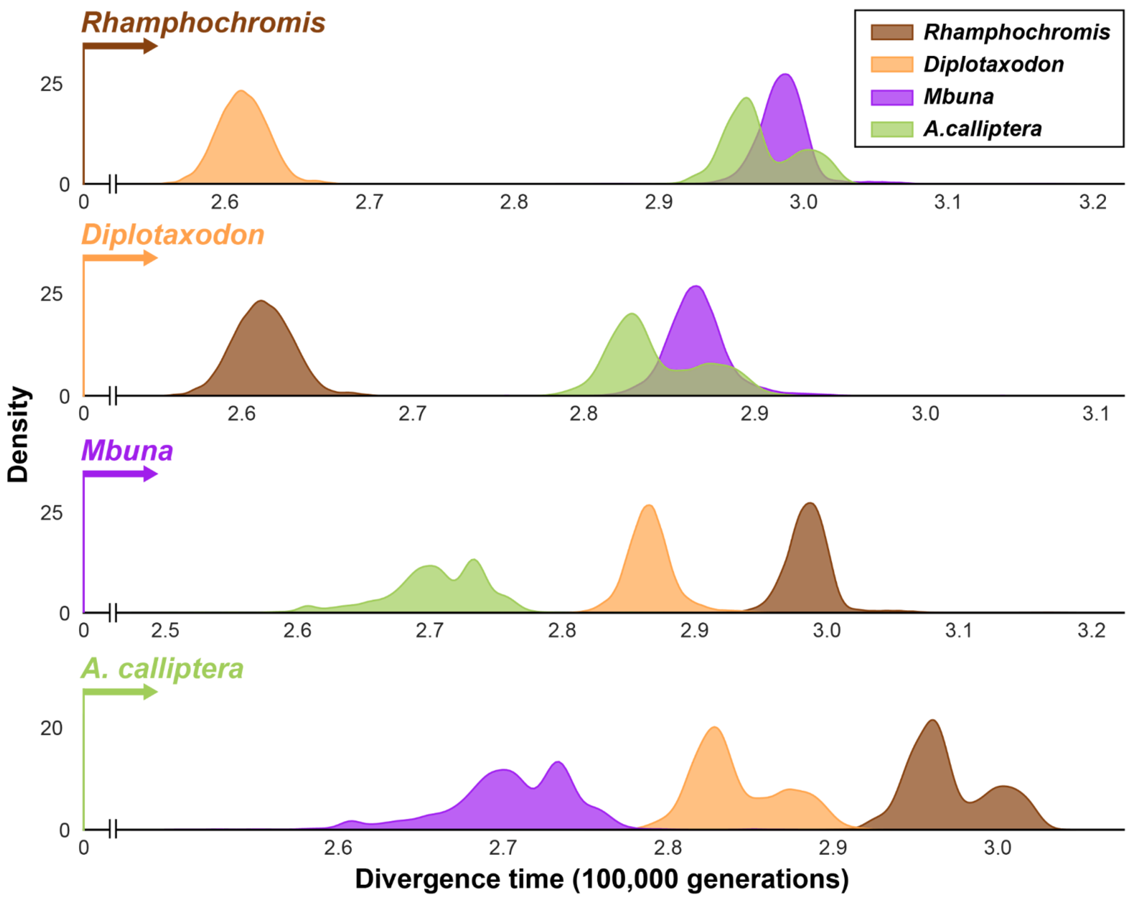

2581

2582 **Fig. S26: Divergence times between non-benthic clades.**

2583 Density plots of pairwise divergence times analogous to Fig. 3A in non-inverted parts of the
2584 genome but between non-benthic clades. The title of each sub-panel gives the focal species
2585 and the density plots show the divergence of this group from other non-benthic groups.

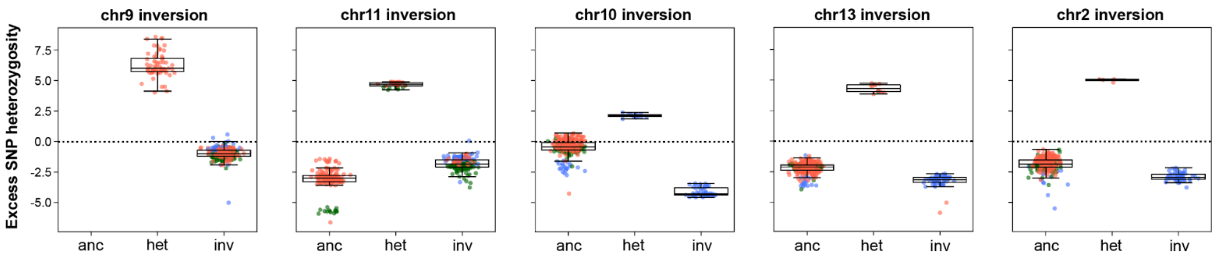

**Fig. S27: SNP heterozygosity by inversion genotype.**

Difference in heterozygosity between inversion regions and rest of the genome, with samples split by their inversion genotype on chromosomes 9, 11, 10, 13, 2. Only samples from species with at least 10 sequenced individuals are included (see materials and methods).

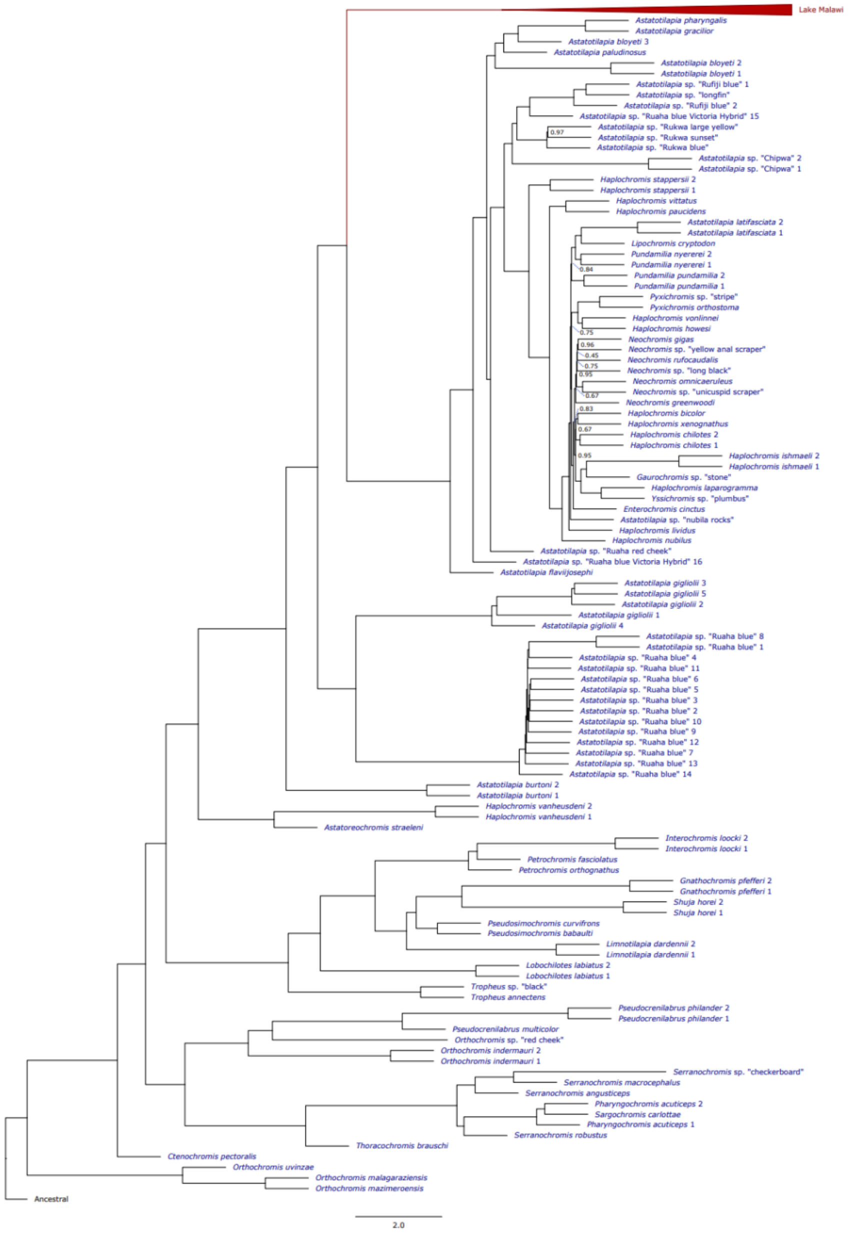

#### Fig. S28: Whole-genome phylogeny of riverine and lacustrine haplochromine cichlids

Genome-wide maximum likelihood (ML) phylogenomic tree (excluding inversion regions) for 612 Malawi (red) and non-Malawi haplochromine cichlid (blue) samples, rooted to a reconstructed haplochromine ancestral sequence (materials and methods). The same individuals were used in the ABBA-BABA analysis to identify the origin of the *benthic* non-inverted chromosome 9 haplotype. Branch support values are the ASTRAL local posterior probability (LPP), which gives the probability the branch is the true branch given the quartet score of gene trees. Only LPP values lower than 1 are shown. The scale bar gives coalescent units and is a measure of the amount of discordance in gene trees.

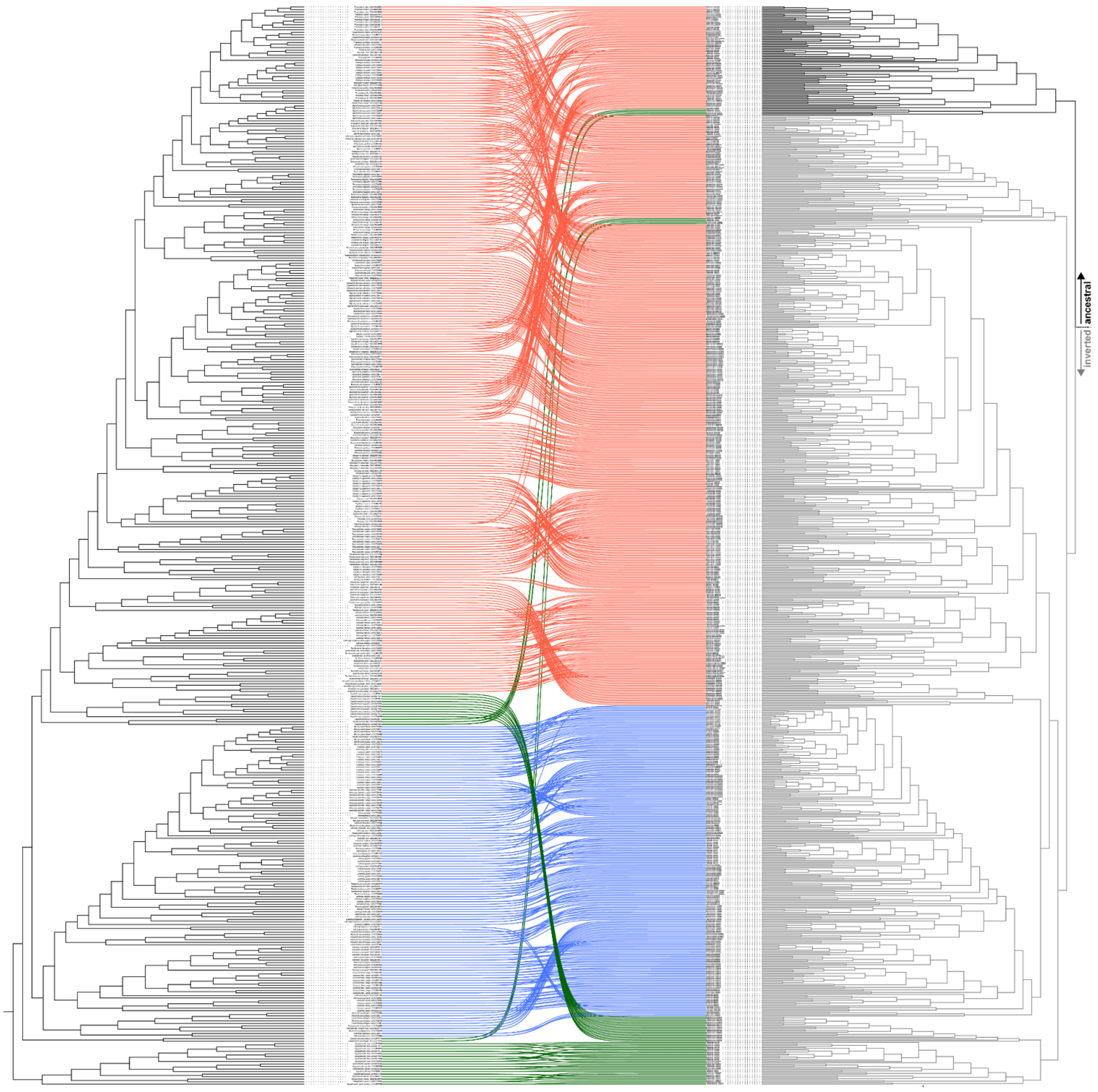

**Fig. S29: Phylogenetic relationships within the chromosome 9 inversion.**

Co-phylogenetic plot of *benthic* samples for the whole genome outside the inversion regions (and excluding chromosome 3) on the left, and for the re-phased chromosome 9 inversion region on the right (see materials and methods). Coloured lines between the left and right tree connect the same individuals in both trees and colour reflects *benthic* clade adherence (red: *shallow benthic*, blue: *deep benthic*, green: *utaka*). A high resolution version of this plot is available as data S4.

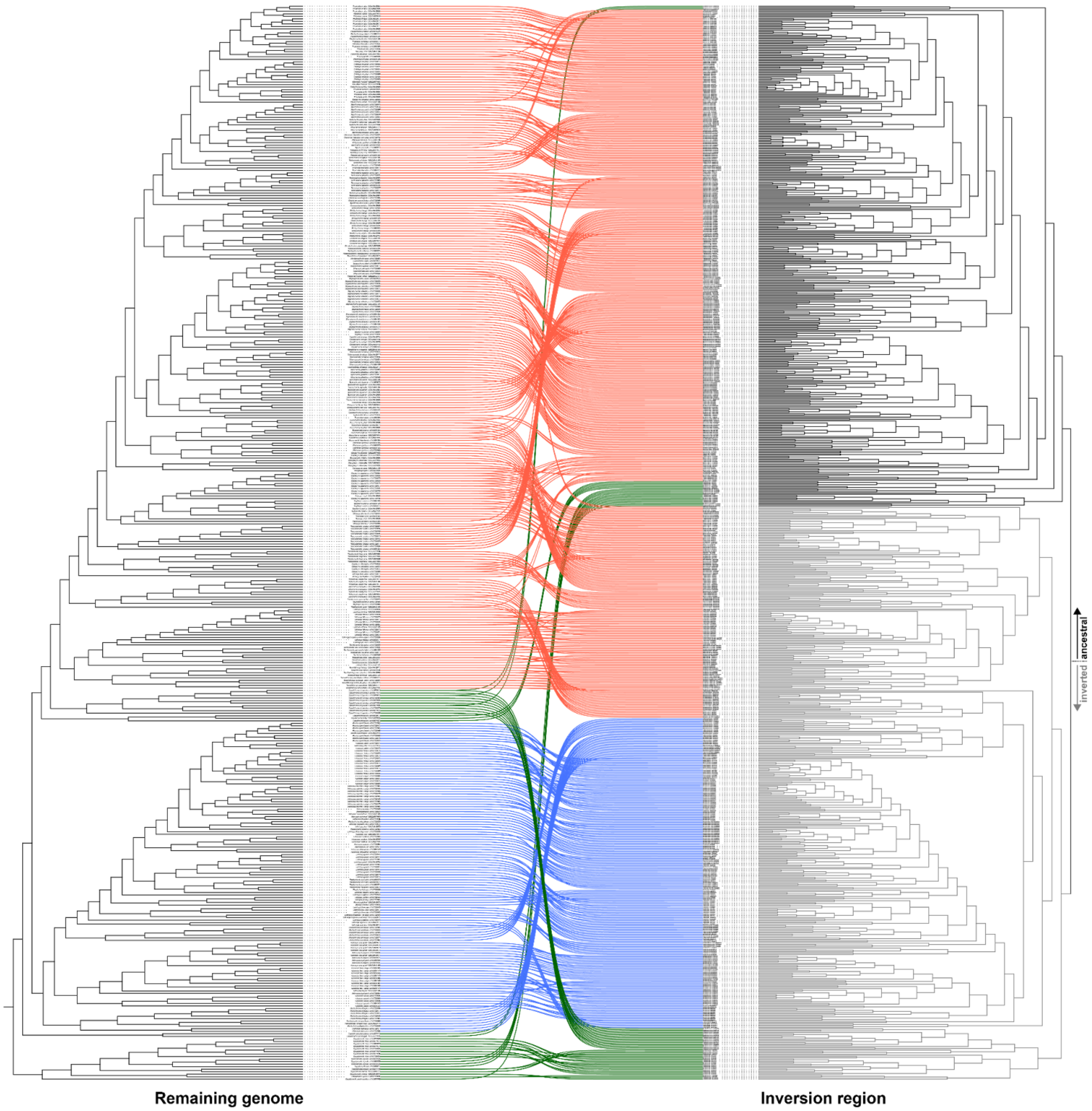

**Fig. S30: Phylogenetic relationships within the chromosome 11 inversion.**

Co-phylogenetic plot of *benthic* samples for the whole genome outside the inversion regions (and excluding chromosome 3) on the left, and for the re-phased chromosome 11 inversion region on the right (see materials and methods). Coloured lines between the left and right tree connect the same individuals in both trees and colour reflects *benthic* clade adherence (red: *shallow benthic*, blue: *deep benthic*, green: *utaka*). A high resolution version of this plot is available as data S5.

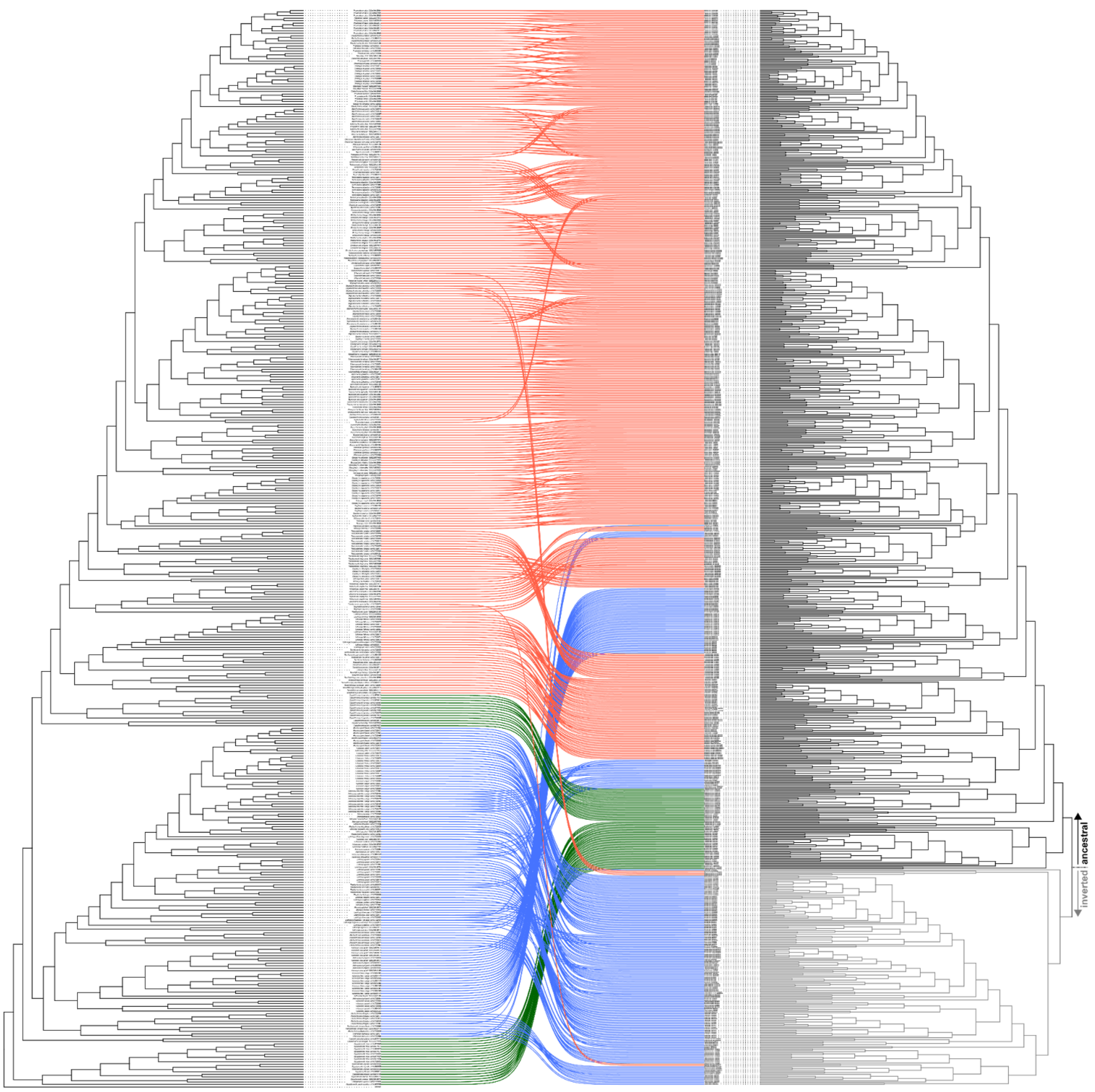

2619

#### 2620 Fig. S31: Phylogenetic relationships within the chromosome 10 inversion.

2621 Co-phylogenetic plot of *benthic* samples for the whole genome outside the inversion regions
2622 (and excluding chromosome 3) on the left, and for the re-phased chromosome 10 inversion
2623 region on the right (see materials and methods). Coloured lines between the left and right tree
2624 connect the same individuals in both trees and colour reflects *benthic* clade adherence (*shallow*
2625 *benthic*, blue: *deep benthic*, green: *utaka*). A high resolution version of this plot is available as
2626 data S6.

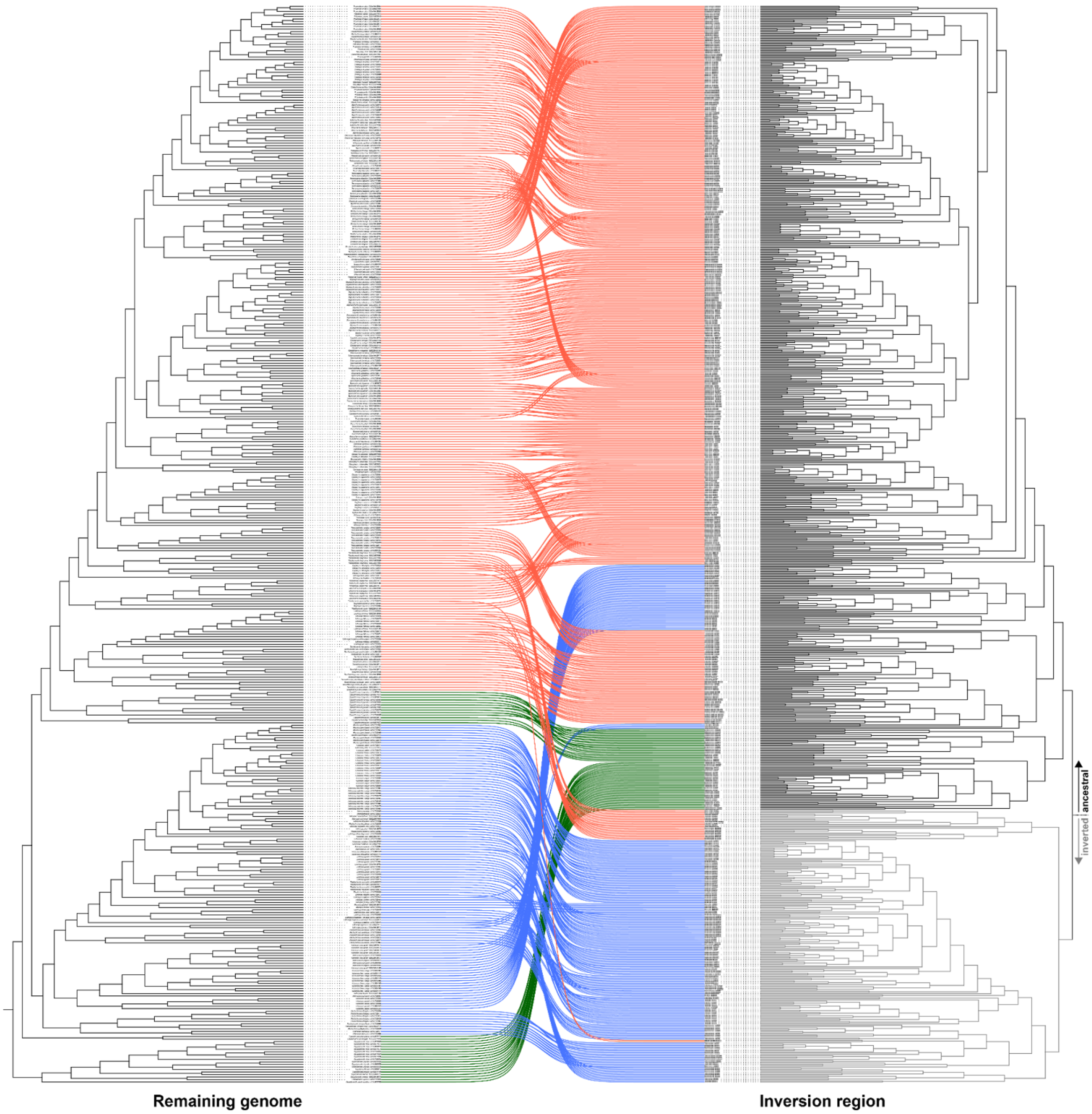

#### Fig. S32: Phylogenetic relationships within the chromosome 13 inversion.

Co-phylogenetic plot of *benthic* samples for the whole genome outside the inversion regions (and excluding chromosome 3) on the left, and for the re-phased chromosome 13 inversion region on the right (see materials and methods). Coloured lines between the left and right tree connect the same individuals in both trees and colour reflects *benthic* clade adherence (*shallow benthic*, blue: *deep benthic*, green: *utaka*). A high resolution version of this plot is available as data S7.

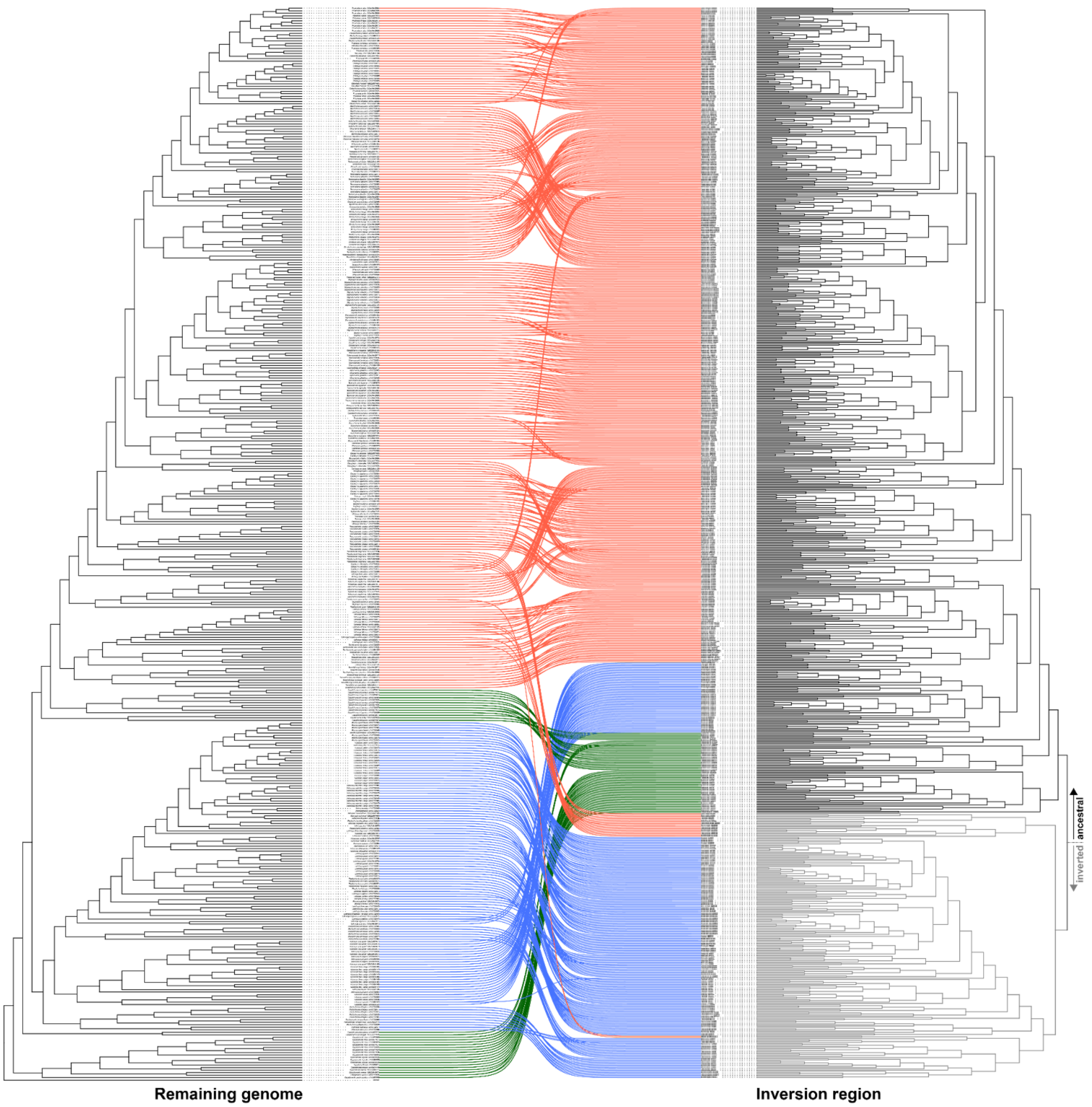

**Fig. S33: Phylogenetic relationships within the chromosome 2 inversion.**

Co-phylogenetic plot of *benthic* samples for the whole genome outside the inversion regions (and excluding chromosome 3) on the left, and for the re-phased chromosome 2 inversion region on the right (see materials and methods). Coloured lines between the left and right tree connect the same individuals in both trees and colour reflects *benthic* clade adherence (*shallow benthic*, blue: *deep benthic*, green: *utaka*). A high resolution version of this plot is available as data S8.

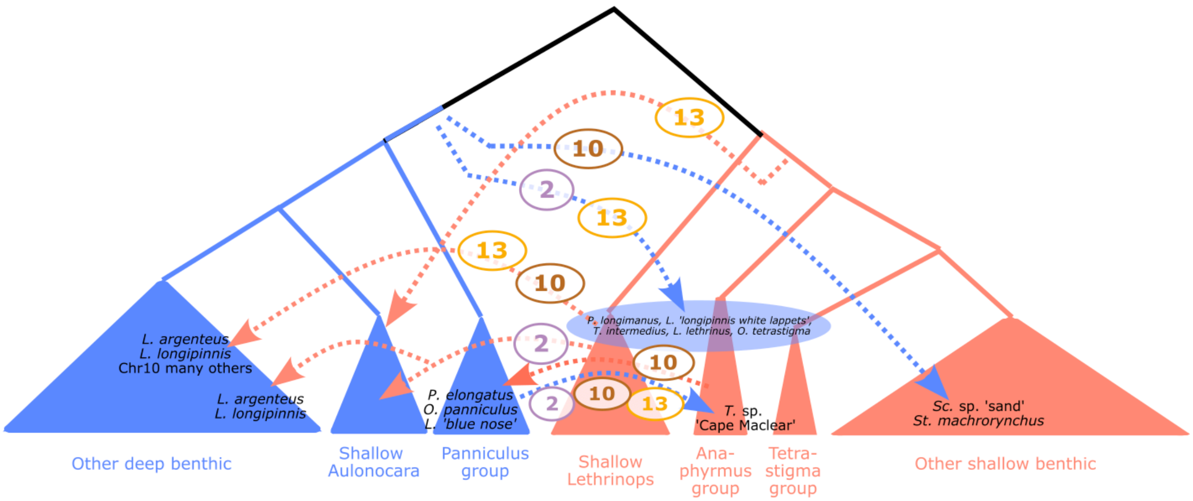

#### Fig. S34: Inversion introgression summary.

Qualitative summary of inversion haplotype transmission events for chromosomes 2, 10, 13 based on investigation of inversion region haplotype phylogenies (figs. S29 to S33). Species are summarised in groups based on the whole genome phylogeny (Fig. 1).

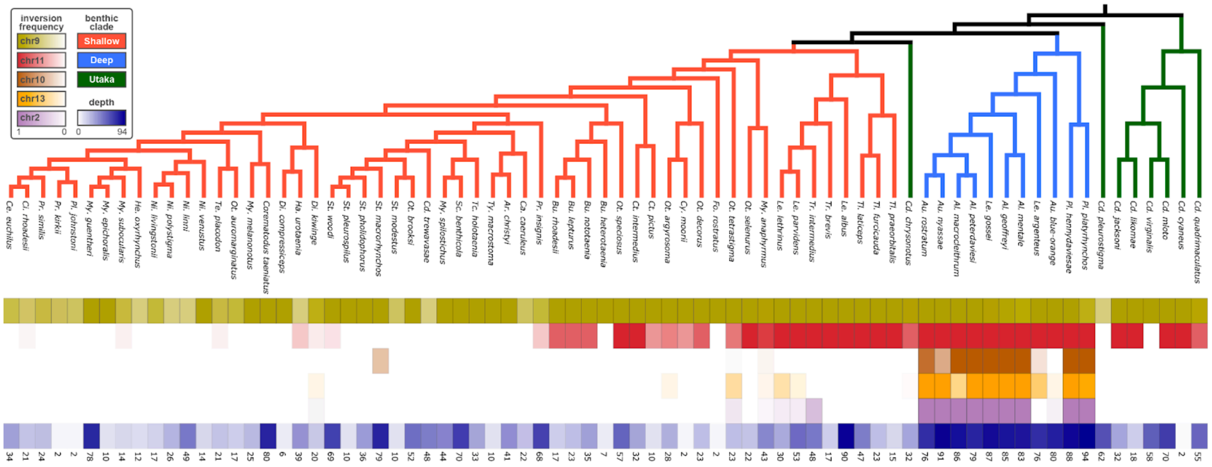

**Fig. S35: Mean habitat depth and inversion state.**

To be able to visually examine the correspondence between species' inversion states and their depth ranges, we aggregated per species depth information (using available literature and our own records) and plotted mean occurrence depth along with inversion frequency below relevant subset of our *benthic* phylogeny. See materials and methods for details on depth data aggregation and plotting.

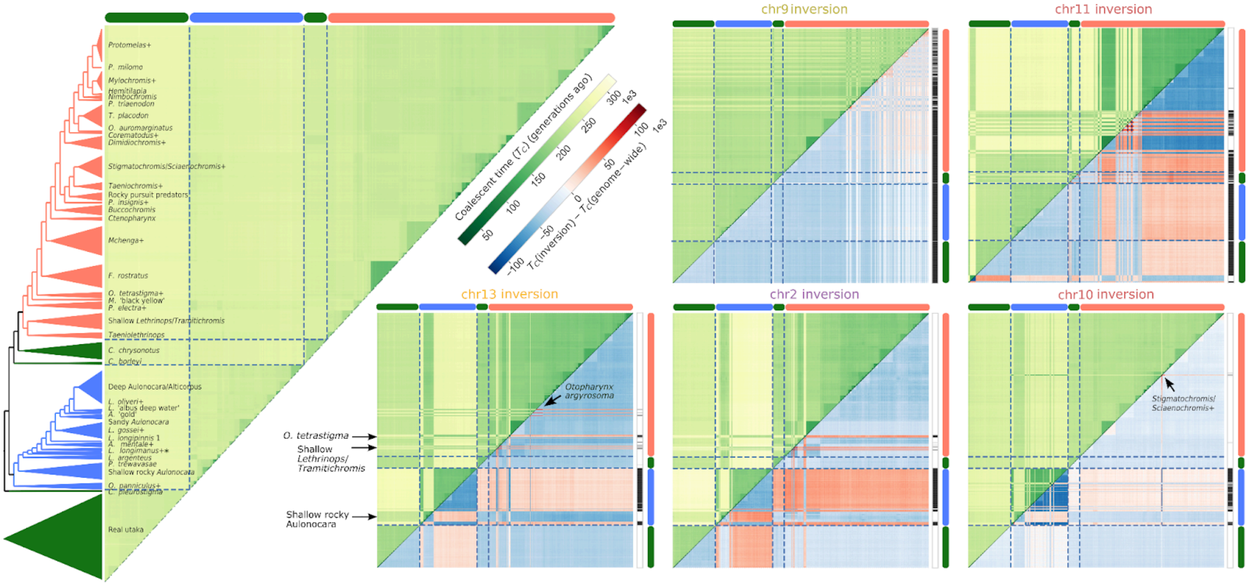

**Fig. S36: Pairwise coalescent times in inverted regions and the rest of the genome.**

Comparison of pairwise haplotype coalescence times in inverted regions and the rest of the genome. Estimates are based on SNP divergences (dxy and π) (see materials and methods). The left larger triangle shows coalescent time estimates for the genome outside of the inversions. Specimens are ordered according to the phylogeny in Fig. 1, which is displayed on the left of the triangle with species grouped for clarity (For group assignments see table S1). Smaller matrices show equivalent coalescent time estimates for inversion regions (green, above diagonal) and the difference between estimates in inversion regions and the rest of the genome (blue/red, below diagonal). The order of specimens is identical across all subpanels. Some groups showing atypical inversion states compared to sister taxa are annotated. Green, blue and orange bars next to matrices annotate adherence to *utaka*, *deep* and *shallow benthics*, respectively. Black/white vertical bars right of matrices annotate inversion states (white: non-inverted; black:inverted).

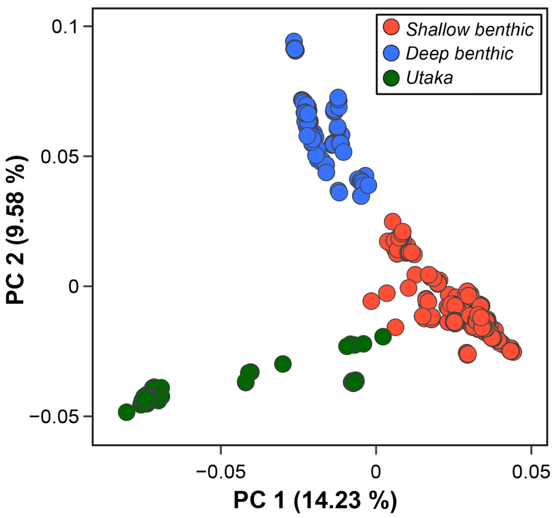

**Fig. S37: Principal component analysis of benthic samples.**

Genome-wide principal component analysis of benthic clades. Each included individual is represented as a single marker and colour-coded by clade-adherence.

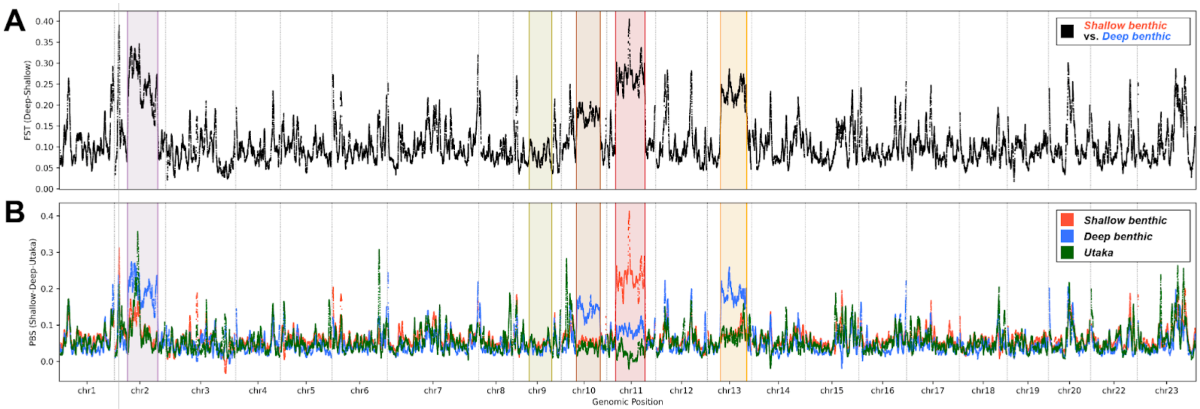

**Fig. S38: Genome-wide divergence patterns in windows.**

(**A**) Fixation index ($F_{ST}$) between most (species with at least 10 sequenced individuals) included *deep* and *shallow benthic* samples in 1 Mbp windows with 10 kbp step size. Inversion regions are highlighted in light red. (**B**) Population branch statistic (PBS), a measure of genetic distance between three populations that captures genetic divergence specific to one clade, among most (species with at least 10 sequenced individuals) *benthic* samples in 1 Mbp windows with 10 kbp step size. Inversion regions are highlighted and colour-coded as in the main text figures.

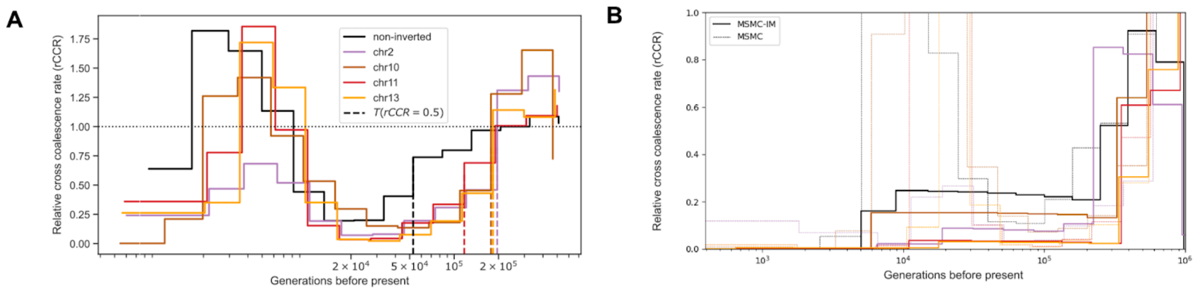

#### Fig. S39: Relative cross-coalescence using MSMC.

(**A**) Cross coalescence computed with MSMC between the *deep benthic* species *Alticorpus peterdaviesi* and the *shallow benthic* species *Trematochranus placodon* (see materials and methods). While *A. peterdaviesi* is fixed for the inverted orientation of all five focal inversions *T. placodon* has no inversions. Cross coalescence was computed separately in inversion regions and for the genome outside of the inversions ("non-inverted"). For each region, the time when cross coalescence first falls below 0.5 – often used as a proxy for speciation – is indicated by dashed vertical lines. We note an increase in cross-coalescence rate for times more recent than $10^4$ generations which we attribute to statistical phasing errors. (**B**) Using MSMC-IM (see materials and methods), the recent increase in cross coalescence disappeared, while the pattern of inversion regions showing earlier reduction in cross coalescence compared to the rest of the genome persisted.

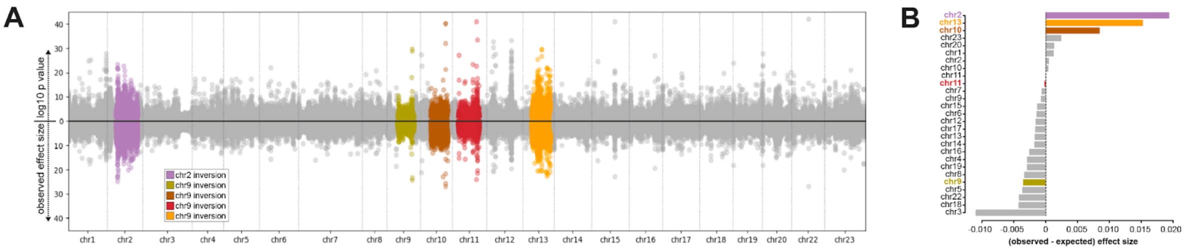

#### Fig. S40: GWAS on clade adherence.

Genome-wide association study on clade adherence between *deep* (n=389) and *shallow* (n=154) *benthics*. (**A**) Manhattan plots showing the significance of SNP genome-wide associations with genetic clade (likelihood ratio test) on the positive axis and estimated effect sizes on the negative axis. (**B**) Proportional effect size contributed per chromosome compared to the null expectation (proportion of the genome spanned by the chromosome). Inversion chromosomes are split into inverted (colour-coded) and non-inverted (grey) regions. Positive values are interpreted as an ‘excess’ of observed effect size, i.e. higher contribution to deep-shallow associated genetic variation than expected.

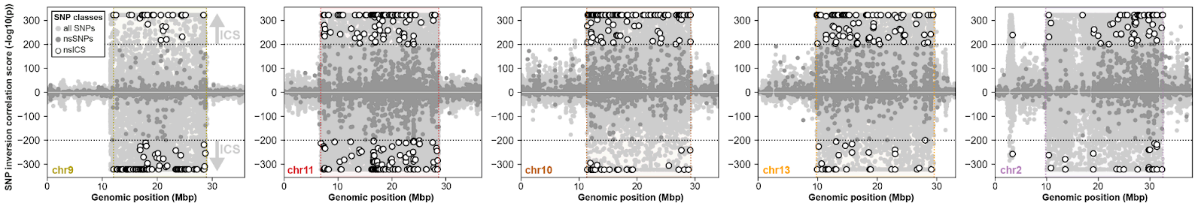

**Fig. S41: Inversion(-state)-correlated SNPs (ICS).**

Visualisation of the different sets of SNPs referred to in the text. Positive correlation scores signify that the derived allele is correlated with the inverted haplotype, while negative correlation scores signify that the derived allele is correlated with the non-inverted haplotype. All SNPs are shown in light grey. SNPs that cause non-synonymous mutations (nsSNPs) are displayed in dark grey. ICS – SNPs that are (highly) correlated with inversion state – have an absolute correlation score of ≥ 200 (indicated by grey arrows). The subset of ICS causing non-synonymous (amino acid-changing) mutations (nsICS) is displayed as white markers with a dark outline.

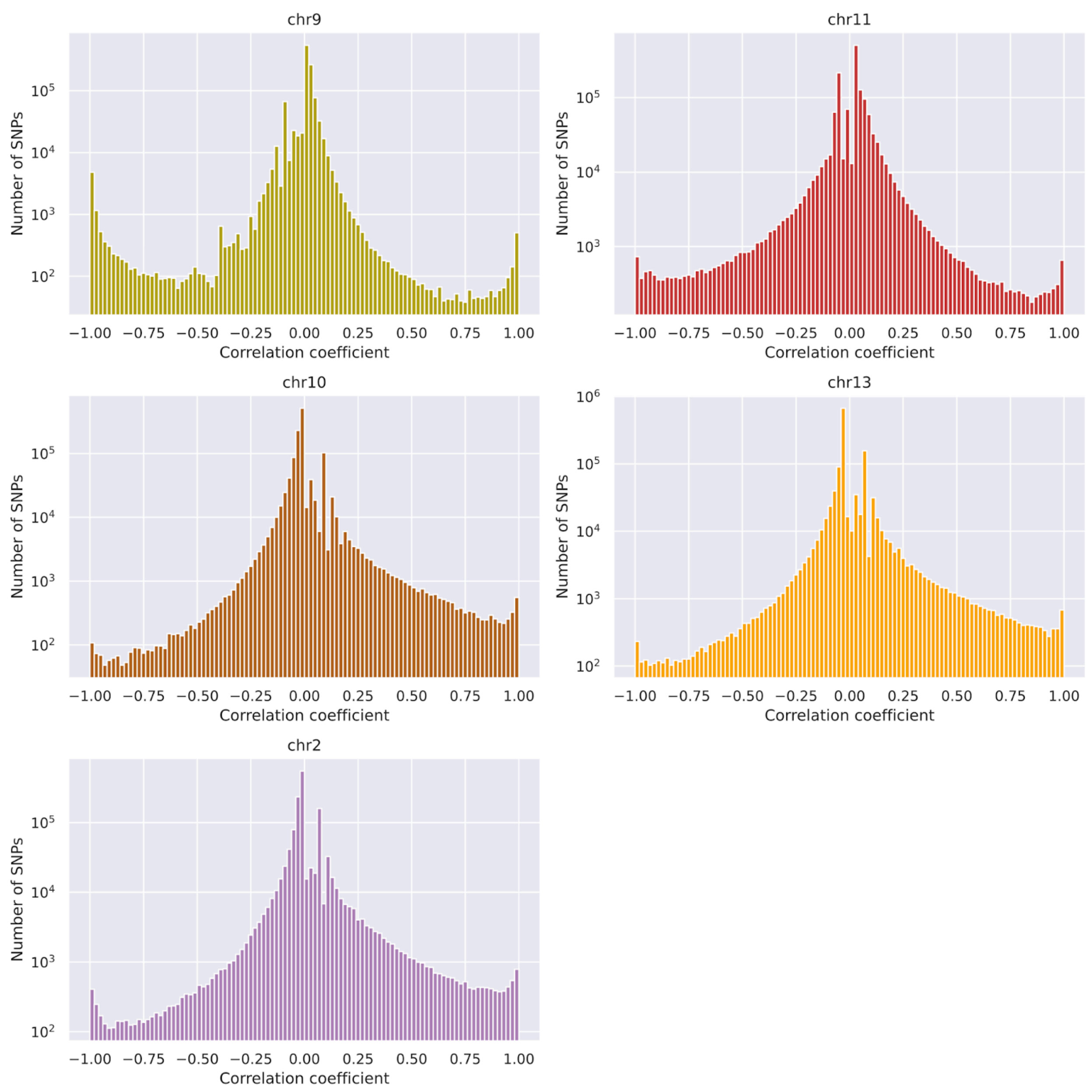

**Fig. S42: Histograms of SNP-inversion correlation coefficients in inversion regions.**

Histograms of SNP-inversion correlation coefficients in inversion regions. Note that the y-axis is on a log scale. While the three inversions common among *deep benthics* (chromosomes 10, 13, 2) show an excess of derived alleles being common on the inverted haplotypes, chromosomes 9 and 11 show an excess of derived alleles on the re-introgressed non-inverted haplotype. This pattern is particularly strong for chromosome 9, consistent with the non-Malawi origin of the non-inverted haplotype.

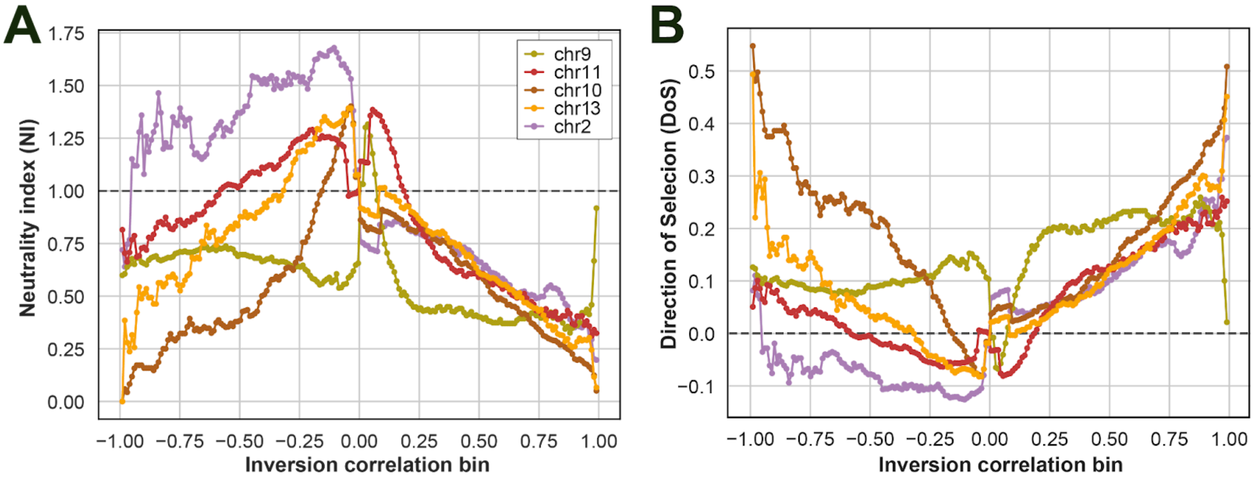

**Fig. S43: Neutrality index and Direction of Selection (DoS) tests.**

Neutrality index and Direction of Selection (DoS) statistics calculated in widening bins of SNPs across each of the five inversion regions (see materials and methods). (**A**) Neutrality index. (**B**) Direction of Selection statistics.

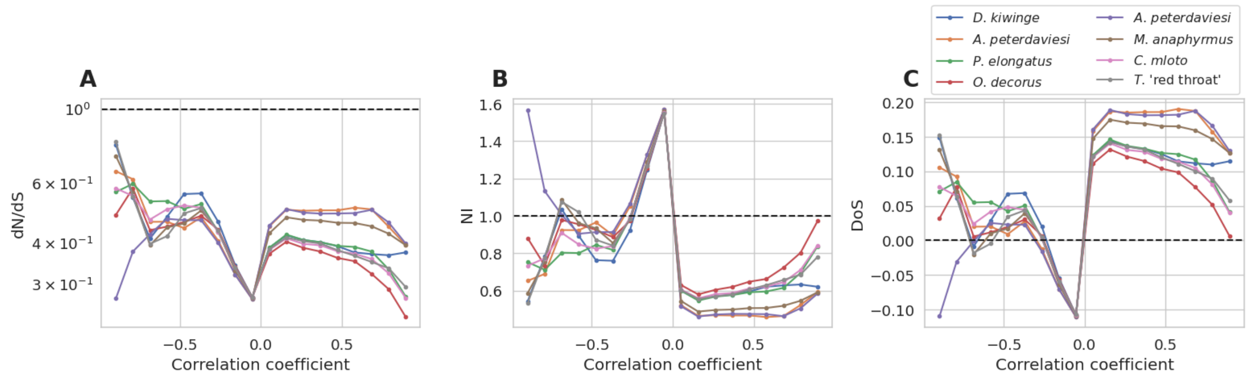

**Fig. S44: Selection statistics on artificial inversion correlated SNPs.**

This figure shows the same selection statistics as displayed for real inversion correlated SNPs in Fig. 4B (dN/dS) and fig. S43 but for artificial ICS produced by an in-silico expansion of randomly picking haplotypes and designating them as "inverted population" (see text S3) with the rationale of mimicking an extreme scenario of rapid growth of a haplotype under positive selection. We see that (**A**) dN/dS does not become > 1.0 in such a scenario, while (**B**) neutrality index, and (**C**) direction of selection statistics attain values below 1.0 and above 0.0, respectively, which could be wrongly interpreted as evidence of positive selection contributing to non-synonymous divergence. The randomly chosen benthic individual from which the haplotype was picked is given in the legend.

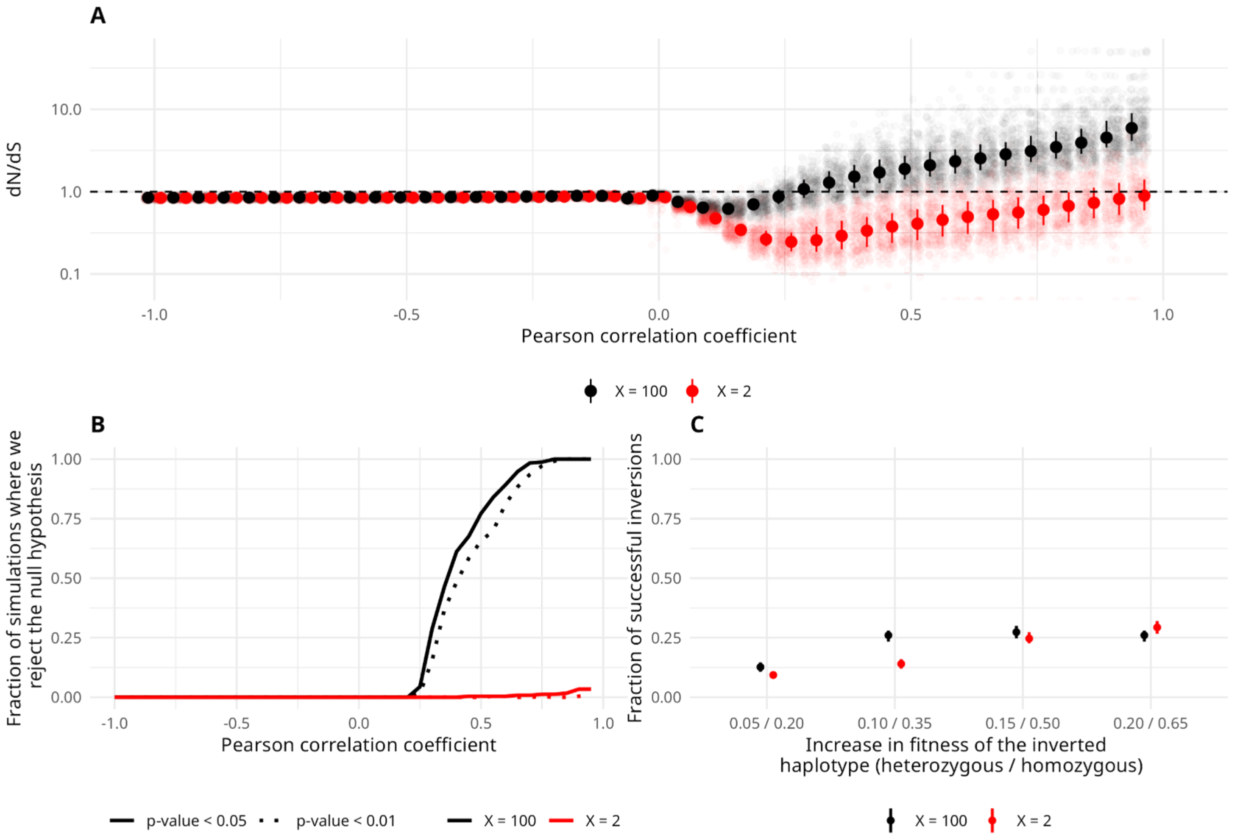

**Fig. S45: Simulations of selection on inversion haplotypes.**

This figure shows results for simulated scenarios of inversion evolution. (see materials and methods and text S3). (**A**) shows the distribution of simulated dN/dS for SNPs as a function of inversion genotype correlation analogous to Fig. 4B. For every bin, each translucid point corresponds to a different replicate. The solid dot shows the median, whereas the vertical lines correspond to the interquartile range. (**B**) For every bin and replicate, we performed a one-tailed test to assess whether the dN/dS was significantly greater than the distribution generated under the null hypothesis (strict neutrality). According to our simulations, it is unlikely to observe a dN/dS significantly greater than 1 for the $X = 2$ scenario (low number of positively selected sites) for the chosen significance thresholds of 0.05 and 0.01. Contrary to this, in the $X = 100$ scenario we rejected the null hypothesis of strict neutrality in every of the 250 replicates for bins with $r \geq 0.8$. (**C**) We studied how the number of conditionally adaptive loci affected the frequency with which inversion was lost due to drift or genetic load in our simulations. The dot indicates the observed frequency, whereas the vertical lines correspond to the bootstrapped interquartile range. The x-axis indicates the increase in fitness in an individual heterozygous or homozygous for the inverted haplotype.

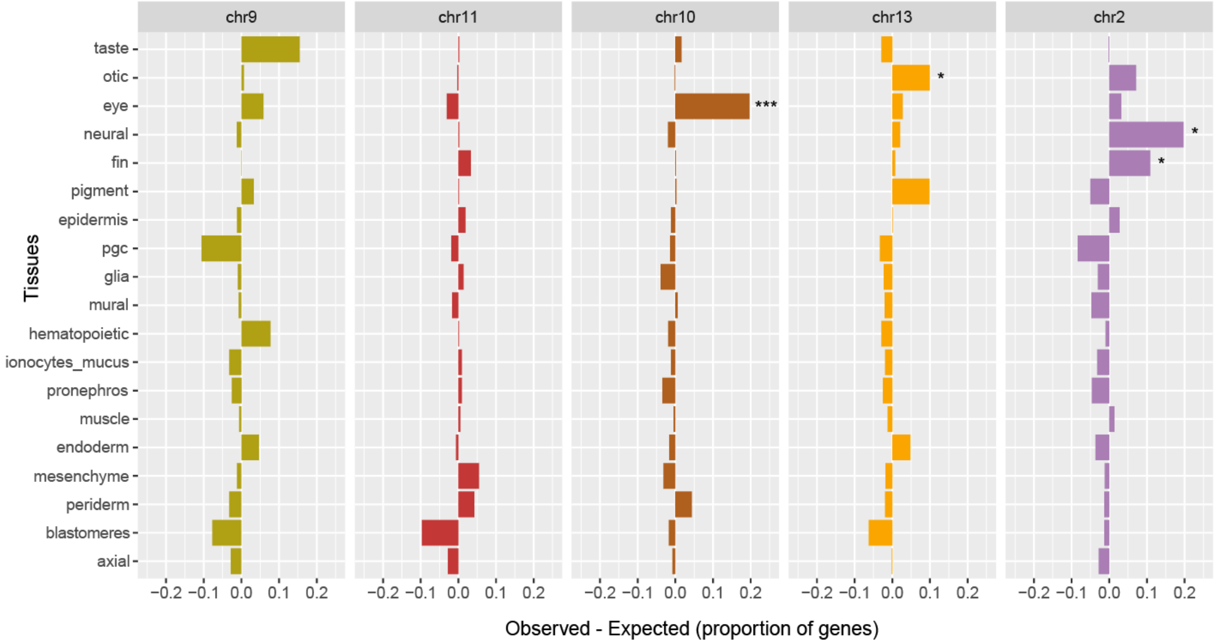

**Fig. S46: Relative expression of genes with nsICS in 5 inversions and 19 tissues.**

Comparison of relative expression of orthologous nsICS genes in different tissues of zebrafish embryo: 5 inversions and 19 tissues. Analysing 441 nsICS genes in 5 inversions, we paid special attention to 315 genes which had an orthologous sequence in the zebrafish single cell expression database "Daniocell". In order to find patterns specific to deep- and shallow water adaptation, we divided 315 target genes into two categories: genes, for which the derived nsICS allele is associated with the inverted haplotype and for which the derived nsICS allele is associated with the non-inverted (ancestral) haplotype. To find tissues in which expression has changed, we compared expression in each group of inversion-associated genes with a set of 12873 genes with matching ID between *Astatotilapia calliptera* reference genome (version 99) and Daniocell database. Tissues which demonstrate significant change of expression are marked with asterisks ($p<0.001$ corresponds to ***, $p<0.01$ corresponds to **, $p<0.05$ corresponds to *, Fisher test with Benjamini-Hochberg multiple test correction). Gene counts and significance are available in table S17.

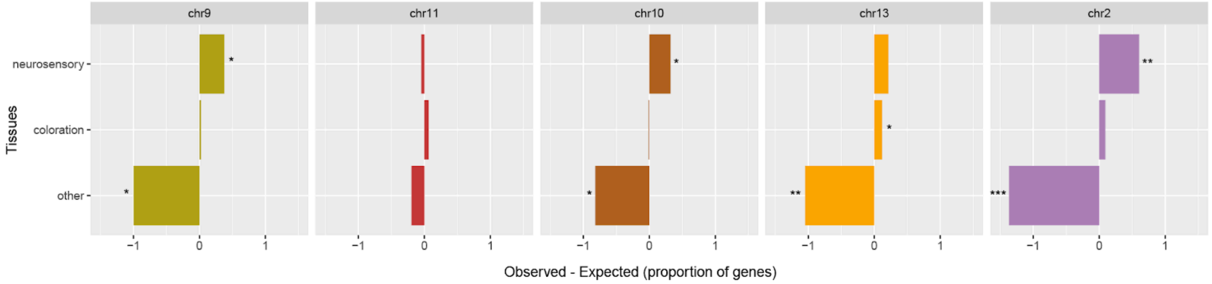

**Fig. S47: Relative expression of genes with nsICS in 5 inversions and 3 grouped tissues.**

Comparison of relative expression of orthologous nsICS genes in different tissues of zebrafish embryo, 5 inversions and 3 grouped tissues. Data and analysis are the same as in fig. S46. Gene counts and significance are available in table S21.

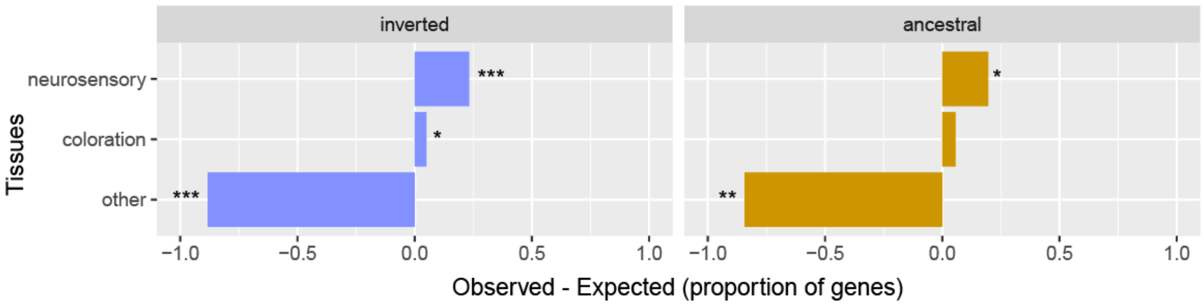

**Fig. S48: Relative expression of genes with nsICS in 2 haplotypes and 3 grouped tissues.**

Comparison of relative expression of orthologous nsICS genes in different tissues of zebrafish embryo: all 5 inversions are grouped, but separated by haplotype; 3 grouped tissues are used. Data and analysis are analogous to fig. S46. The left panel (“inverted”) shows results for genes with positive nsICS (i.e., the derived allele present on the inverted haplotype), while the right panel (“ancestral") shows results for genes with negative nsICS (i.e., the derived allele present on the non-inverted haplotype). Gene counts and significance are available in table S22.

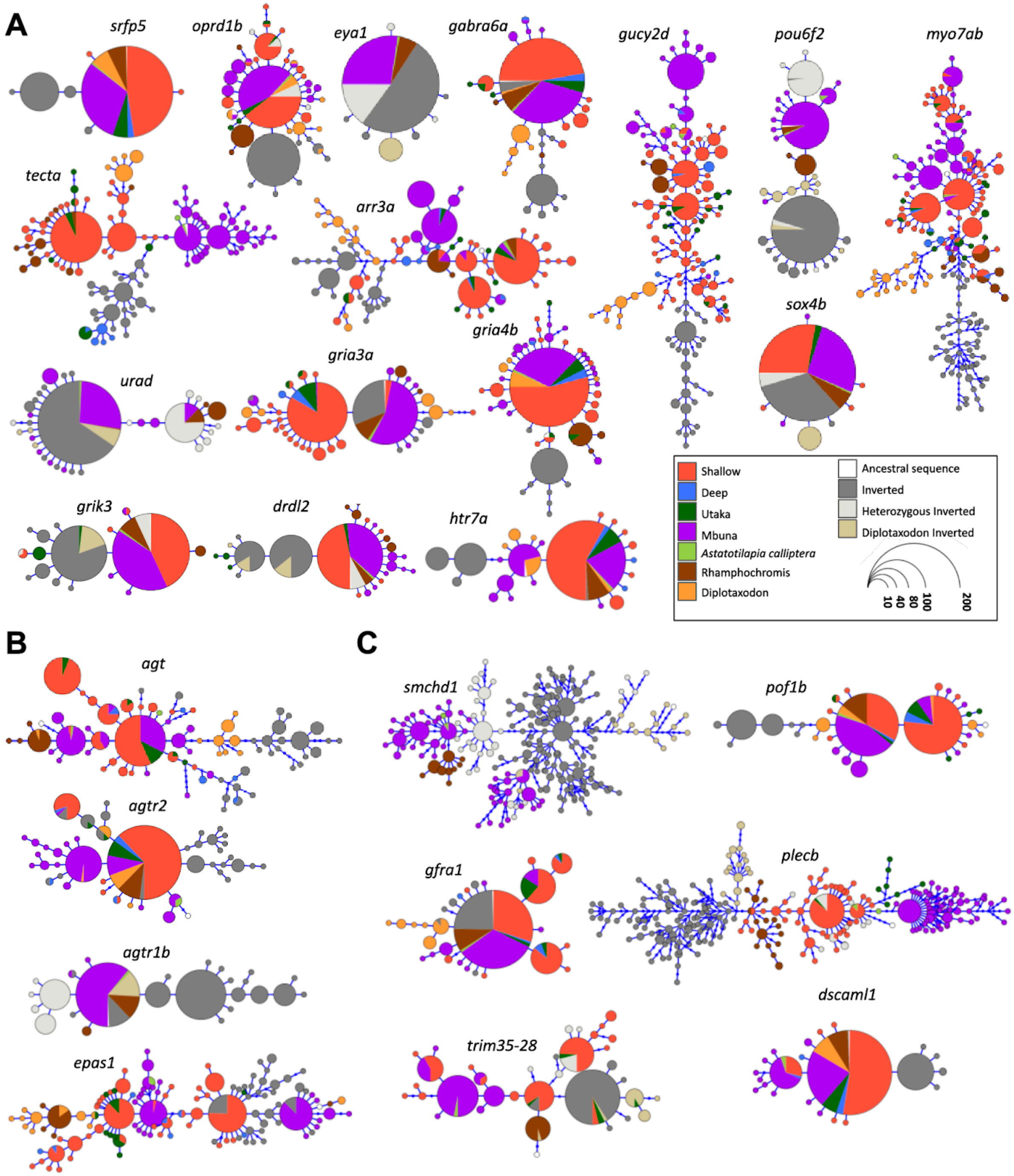

2793

2794 **Fig. S49: Haplotype trees for genes of interest.**

2795 Haplotype trees for genes relevant in (**A**) sensory adaptation and behaviour, (**B**) vascular
2796 system regulation and (**C**) adaptations related to sex and reproduction. Colours signify different
2797 clades and grey represents inverted haplotypes, except *Diplotaxodon*, for which inverted
2798 haplotypes are displayed in their own colour. The inverted haplotype is often separated from the
2799 non-inverted benthic haplotypes in these candidate genes for adaptive evolution.

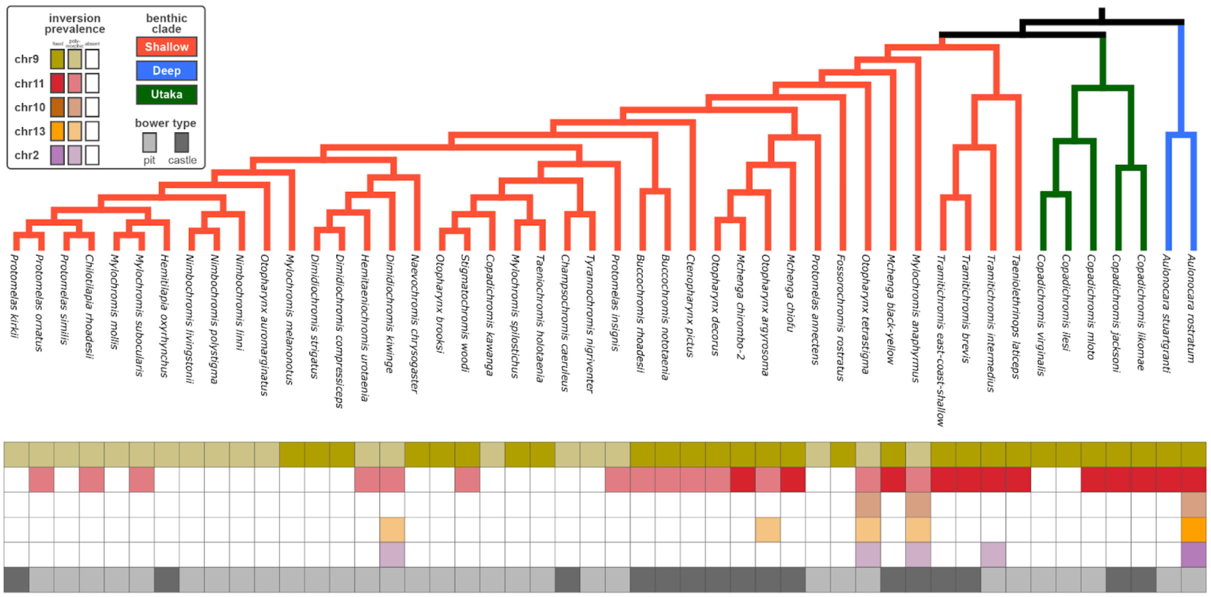

#### Fig. S50: Bower type and inversion state.

To be able to visually examine the correspondence between species' inversion states and their basic bower layout, we obtained per species information. We plotted bower type ('pit' vs. 'castle') and inversion prevalence per species (fixed vs. polymorphic vs. absent) below the relevant subset of our *benthic* phylogeny. See materials and methods for details on bower data and plotting.

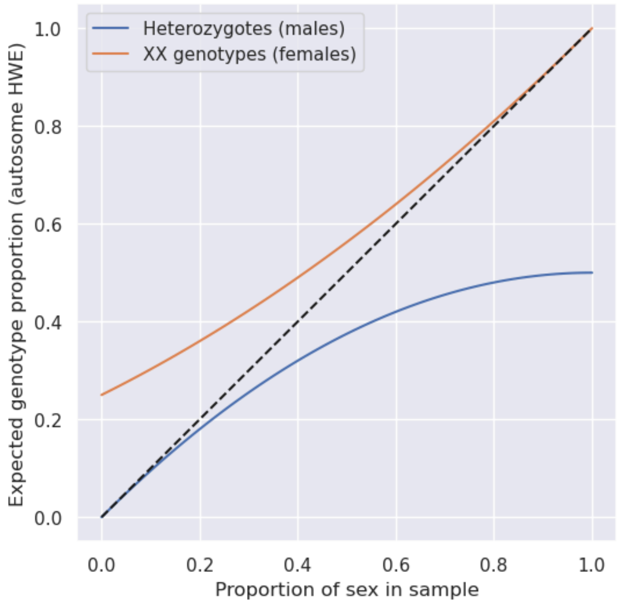

#### Fig. S51: Genotype proportions of a sex locus.

Observed genotype proportions for a sex-locus compared to expectations for an autosomal locus under HWE. The x axis gives observed genotype proportions when a perfectly sex-linked locus is sampled for a given sex-ratio in the sample. The fact that the blue line is always below the diagonal demonstrates that sex-loci show an excess of heterozygotes compared to an autosomal locus under HWE. For example, for a sex locus in a sample with sex ratio of 1/2, 50% heterozygotes are observed, while 37.5% would be expected under autosomal HWE.

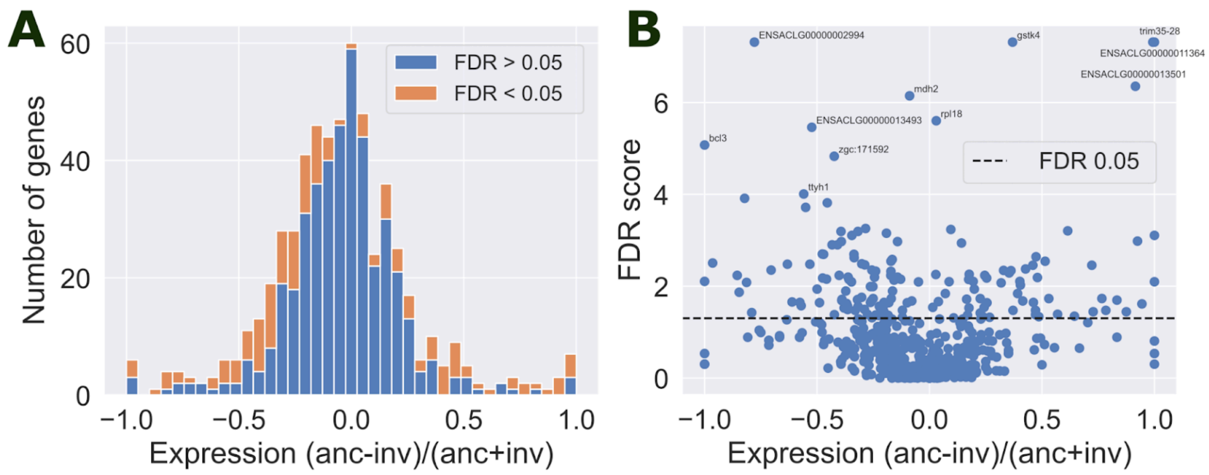

**Fig. S52: Allele specific expression.**

Allele specific expression (ASE) within the chromosome 11 inversion region across five tissues of eleven males of *Copadichromis chrysonotus* all heterozygous for the inversion (see materials and methods). (**A**) Histogram of ASE values. (**B**) ASE values plotted against an FDR score (-log10(p-value)) obtained from applying Benjamini-Hochberg multiple testing correction to Wilcoxon signed-rank test tests on RNAseq read counts. Full data is given in table S24.

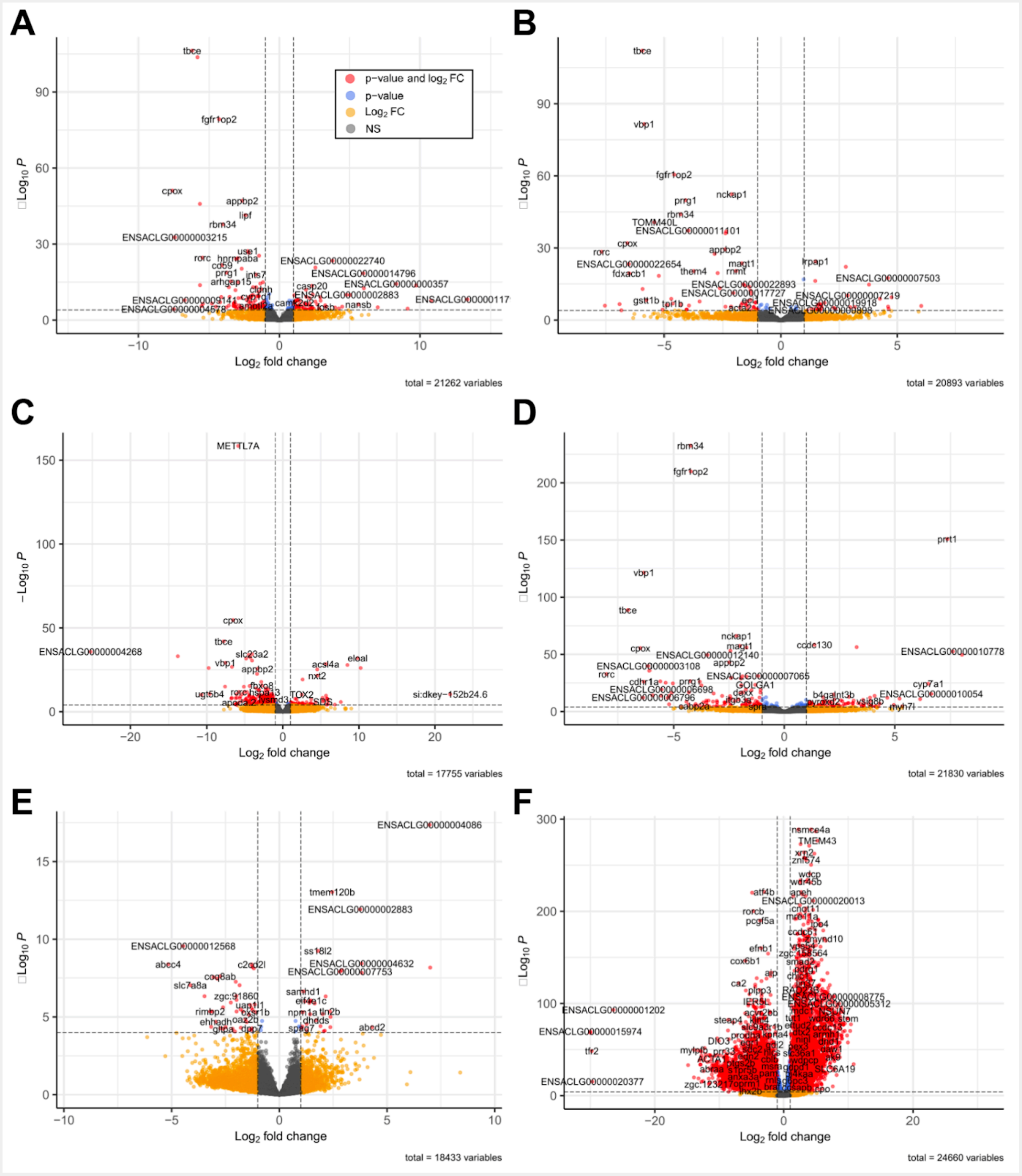

### Fig. S53: Differential gene expression.

(**A**)-(**E**) Differentially expressed genes between *C.mloto* and *C.chrysonotus* by tissue: (**A**) Gills, (**B**) Brain, (**C**) Liver, (**D**) Gonads, (**E**) Muscle). (**F**) Genes, which are differentially expressed in gonads relative to other tissues in both species. $p_{adj}$ threshold = 0.05, abs(log2FC) threshold = 1.

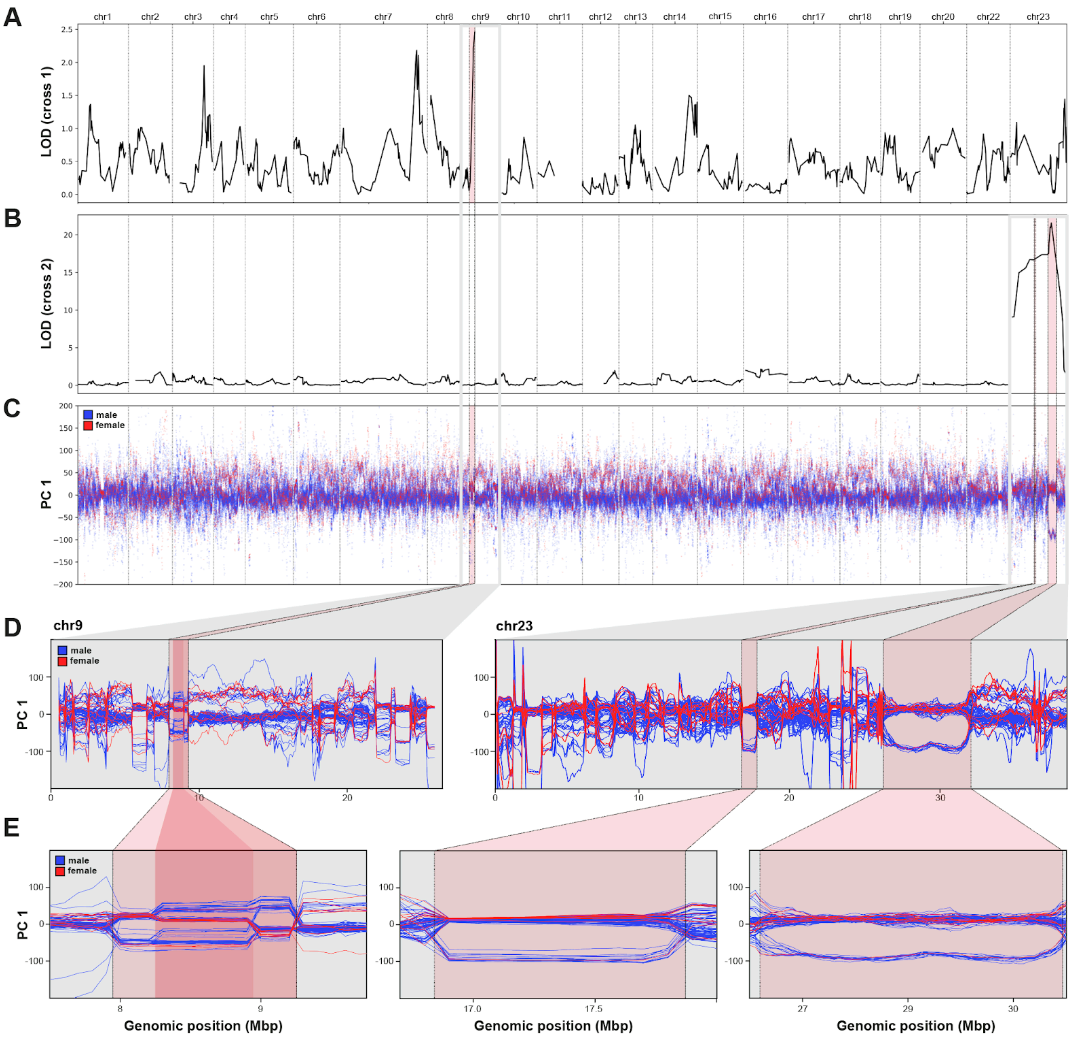

#### Fig. S54: (Putative) sex-linked inversions in the Lake Victoria radiation.

Using existing radiation-wide sequencing data for lake 100 Lake Victoria cichlid species (*177*), we identified three putative inversion regions that all overlap with previously mapped sex determination loci. (**A**) Re-plotted LOD scores (from Feller et al. (*58*)) for sex association in an interspecific QTL cross (*Pundamillia* sp. 'nyererei-like' x *P*. sp. 'pundamilia-like') against genomic position in *P. nyererei*. They identified the association peak on chromosome 9 to represent a ZW system. (**B**) Re-plotted LOD scores (from Feller et al. (*58*)) for sex association in a second interspecific QTL cross (*P. pundamilia* x *P*. sp. 'red-head') against genomic position in *P. nyererei*. They identified the association peak on chromosome 9 to represent an XY system. (**C**) Windowed PC analysis of 100 Lake Victoria species, with each blue line representing a male, and each red line representing a female individual. (**D**) When examining the data from (**C**) chromosome-wise and at higher resolution, we identified three putative inversion regions on chromosomes 9 and 23. (**E**) Zoom in on the putative inversion regions. The region on chromosome 9 appears to be composed of at least two overlapping inversions, forming characteristic strata in which certain inversion genotypes comprise almost exclusively male individuals. The region perfectly matches the QTL peak in (**A**). On chromosome 23, two separate putative inversion regions exist, both falling within the QTL peak in (**B**), and forming characteristic strata in which one inversion genotypes comprises almost exclusively male individuals.

### Tables S1 to S31

#### Table S1: Sample metadata of the main analysis (separate file).

Metadata of the (mostly) wild-caught cichlid samples contained in the variant callset used for the main analysis. Per sample, the file contains genus and species name, clade assignment, catch location (and a more specific sublocation, if available), sex assignment (see materials and methods), sequence ID (used as sample ID in the VCF), sample/individual ID, repository accessions (BioSample and BioProject) and sequencing depth.

**Supplementary file "Table_S1.tsv"**

#### Table S2: Inversion frequency per species (separate file).

Frequency of each focal inversion per species (and non-*benthic* clade), combining PCA-based genotyping of whole genome-sequenced samples (table S7) with PCR-based genotyping of additional samples (table S11). Per *benthic* species (and for joint outgroup clades *Diplotaxodon*, *Rhamphochromis*, *A. calliptera* and *mbuna*), the frequency of each inversion is reported in separate columns. Because of occasional dropouts in the PCR-based typing, the total number of typed samples (PCA + PCR) can vary slightly between inversions for the same species. Therefore, we report the total number of typed individuals ('nind') in separate columns for each inversion. For details of both inversion genotyping approaches, see materials and methods.

**Supplementary file "Table_S2.tsv"**

#### Table S3: Species name abbreviations.

Species names in figure 1 are abbreviated for space reasons. The following abbreviations were used to refer to genera: *Alticorpus*: Al, *Aristochromis*: Ar, *Aulonocara*: Au, *Buccochromis*: Bu, *Champsochromis*: Ca, *Cheilochromis*: Ce, *Chilotilapia*: Ci, *Copadichromis*: Cd, *Corematodus*: Cm, *Ctenopharynx*: Ct, *Cyrtocara*: Cy, *Dimidiochromis*: Di, *Fossorochromis*: Fo, *Hemitilapia*: He, *Hemitaeniochromis*: Ha, *Lethrinops*: Le, *Mchenga*: Mc, *Mylochromis*: My, *Naevochromis*: Na, *Nimbochromis*: Ni, *Otopharynx*: Ot, *Placidochromis*: Pl, *Protomelas*: Pr, *Sciaenochromis*: Sc, *Stigmatochromis*: St, *Taeniochromis*: Tc, *Taeniolethrinops*: Tl, *Tramitichromis*: Tr, *Trematocranus*: Te, *Tyrannochromis*: Ty. The table below displays all species names and corresponding abbreviations used in Fig. 1.

| Species | abbreviated | Species | abbreviated |
|---|---|---|---|
| Alticorpus geoffreyi | Al. geoffreyi | Mylochromis ensatus | My. ensatus |
| Alticorpus macrocleithrum | Al. macrocleithrum | Mylochromis epichoralis | My. epichoralis |
| Alticorpus mentale | Al. mentale | Mylochromis guentheri | My. guentheri |
| Alticorpus peterdaviesi | Al. peterdaviesi | Mylochromis labidodon | My. labidodon |
| Aristochromis christyi | Ar. christyi | Mylochromis melanonotus | My. melanonotus |
| Aulonocara blue-orange | Au. blue O | Mylochromis mollis | My. mollis |
| Aulonocara copper | Au. copper | Mylochromis obtusus | My. obtusus |
| Aulonocara deep | Au. deep | Mylochromis spilostichus | My. spilostichus |
| Aulonocara ethelwynnae | Au. ethelwynnae | Mylochromis subocularis | My. subocularis |
| Aulonocara gold | Au. gold | Naevochromis chrysogaster | Na. chrysogaster |
| Aulonocara maisoni | Au. maisoni | Nimbochromis linni | Ni. linni |
| Aulonocara malembo-orange | Au. malembo O | Nimbochromis livingstonii | Ni. livingstonii |
| Aulonocara minutus | Au. minutus | Nimbochromis polystigma | Ni. polystigma |
| Aulonocara nyassae | Au. nyassae | Nimbochromis venustus | Ni. venustus |
| Aulonocara rostratum | Au. rostratum | Otopharynx argyrosoma | Ot. argyrosoma |
| Aulonocara saulosi | Au. saulosi | Otopharynx auromarginatus | Ot. auromarginatus |
| Aulonocara six-bar | Au. six B | Otopharynx brooksi | Ot. brooksi |
| Aulonocara stonemani | Au. stonemani | Otopharynx brooksi-striped | Ot. brooksi S |
| Aulonocara stuartgranti | Au. stuartgranti | Otopharynx decorus | Ot. decorus |
| Aulonocara stuartgranti-cape-maclear | Au. stuartgranti CM | Otopharynx heterodon-nankumba | Ot. heterodon N |
| Aulonocara yellow | Au. yellow | Otopharynx lithobates | Ot. lithobates |
| Buccochromis heterotaenia | Bu. heterotaenia | Otopharynx panniculus | Ot. panniculus |
| Buccochromis lepturus | Bu. lepturus | Otopharynx selenurus | Ot. selenurus |
| Buccochromis nototaenia | Bu. nototaenia | Otopharynx speciosus | Ot. speciosus |
| Buccochromis rhoadesii | Bu. rhoadesii | Otopharynx styrax | Ot. styrax |
| Buccochromis whitefins | Bu. whitefins | Otopharynx tetraspilus | Ot. tetraspilus |
| Champsochromis caeruleus | Ca. caeruleus | Otopharynx tetrastigma | Ot. tetrastigma |
| Cheilochromis euchilus | Ce. euchilus | Placidochromis acutirostris | Pl. acutirostris |
| Chilotilapia rhoadesii | Ci. rhoadesii | Placidochromis boops | Pl. boops |
| Copadichromis borleyi | Cd. borleyi | Placidochromis electra | Pl. electra |
| Copadichromis chrysonotus | Cd. chrysonotus | Placidochromis elongatus | Pl. elongatus |
| Copadichromis cyaneus | Cd. cyaneus | Placidochromis hennydaviesae | Pl. hennydaviesae |
| Copadichromis ilesi | Cd. ilesi | Placidochromis johnstoni | Pl. johnstoni |
| Copadichromis jacksoni | Cd. jacksoni | Placidochromis longimanus | Pl. longimanus |
| Copadichromis kawanga | Cd. kawanga | Placidochromis mbunoides | Pl. mbunoides |
| Copadichromis likomae | Cd. likomae | Placidochromis milomo | Pl. milomo |
| Copadichromis mloto | Cd. mloto | Placidochromis nkhotakotae | Pl. nkhotakotae |
| Copadichromis pleurostigma | Cd. pleurostigma | Placidochromis obscurus | Pl. obscurus |
| Copadichromis quadrimaculatus | Cd. quadrimaculatus | Placidochromis platyrhynchos | Pl. platyrhynchos |

| | | | |
|---|---|---|---|
| ***Copadichromis trewavasae*** | *Cd. trewavasae* | ***Placidochromis trewavasae*** | *Pl. trewavasae* |
| ***Copadichromis verduyni*** | *Cd. verduyni* | ***Protomelas annectens*** | *Pr. annectens* |
| ***Copadichromis virginalis*** | *Cd. virginalis* | ***Protomelas chilumba*** | *Pr. chilumba* |
| ***Corematodus taeniatus*** | *Cm. taeniatus* | ***Protomelas insignis*** | *Pr. insignis* |
| ***Ctenopharynx intermedius*** | *Ct. intermedius* | ***Protomelas kirkii*** | *Pr. kirkii* |
| ***Ctenopharynx pictus*** | *Ct. pictus* | ***Protomelas ornatus*** | *Pr. ornatus* |
| ***Dimidiochromis compressiceps*** | *Di. compressiceps* | ***Protomelas similis*** | *Pr. similis* |
| ***Dimidiochromis kiwinge*** | *Di. kiwinge* | ***Protomelas taeniolatus*** | *Pr. taeniolatus* |
| ***Dimidiochromis strigatus*** | *Di. strigatus* | ***Protomelas triaenodon*** | *Pr. triaenodon* |
| ***Fossorochromis rostratus*** | *Fo. rostratus* | ***Sciaenochromis benthicola*** | *Sc. benthicola* |
| ***Hemitaeniochromis brachyrhynchus*** | *Ha. brachyrhynchus* | ***Sciaenochromis fryeri*** | *Sc. fryeri* |
| ***Hemitaeniochromis deep*** | *Ha. deep* | ***Sciaenochromis nyassae*** | *Sc. nyassae* |
| ***Hemitaeniochromis urotaenia*** | *Ha. urotaenia* | ***Sciaenochromis sand*** | *Sc. sand* |
| ***Hemitilapia oxyrhynchus*** | *He. oxyrhynchus* | ***Stigmatochromis macrorhynchos*** | *St. macrorhynchos* |
| ***Lethrinops albus*** | *Le. albus* | ***Stigmatochromis melanchros*** | *St. melanchros* |
| ***Lethrinops albus-deep-water*** | *Le. albus DW* | ***Stigmatochromis modestus*** | *St. modestus* |
| ***Lethrinops albus-green-head*** | *Le. albus GH* | ***Stigmatochromis pholidophorus*** | *St. pholidophorus* |
| ***Lethrinops argenteus*** | *Le. argenteus* | ***Stigmatochromis pleurospilus*** | *St. pleurospilus* |
| ***Lethrinops atrilabris*** | *Le. atrilabris* | ***Stigmatochromis woodi*** | *St. woodi* |
| ***Lethrinops blue-Chilumba*** | *Le. blue C* | ***Taeniochromis holotaenia*** | *Tc. holotaenia* |
| ***Lethrinops blue-nose*** | *Le. blue N* | ***Taeniolethrinops furcicauda*** | *Tl. furcicauda* |
| ***Lethrinops deepwater-altus*** | *Le. deepwater A* | ***Taeniolethrinops furcicauda-yellow*** | *Tl. furcicauda Y* |
| ***Lethrinops gossei*** | *Le. gossei* | ***Taeniolethrinops laticeps*** | *Tl. laticeps* |
| ***Lethrinops lethrinus*** | *Le. lethrinus* | ***Taeniolethrinops macrorhynchus*** | *Tl. macrorhynchus* |
| ***Lethrinops longimanus-red-head*** | *Le. longimanus RH* | ***Taeniolethrinops praeorbitalis*** | *Tl. praeorbitalis* |
| ***Lethrinops longipinnis-blue-head*** | *Le. longipinnis BH* | ***Taeniolethrinops praeorbitalis-northern*** | *Tl. praeorbitalis N* |
| ***Lethrinops longipinnis-white-lappets*** | *Le. longipinnis WL* | ***Tramitichromis brevis*** | *Tr. brevis* |
| ***Lethrinops oculatus*** | *Le. oculatus* | ***Tramitichromis east-coast-shallow*** | *Tr. east CS* |
| ***Lethrinops oliveri*** | *Le. oliveri* | ***Tramitichromis intermedius*** | *Tr. intermedius* |
| ***Lethrinops parvidens*** | *Le. parvidens* | ***Tramitichromis kande*** | *Tr. kande* |
| ***Mchenga black-yellow*** | *Mc. black Y* | ***Tramitichromis red-throat*** | *Tr. red T* |
| ***Mchenga chiofu*** | *Mc. chiofu* | ***Trematocranus cape-maclear*** | *Te. cape M* |
| ***Mchenga chirombo-1*** | *Mc. chirombo 1* | ***Trematocranus placodon*** | *Te. placodon* |
| ***Mchenga chirombo-2*** | *Mc. chirombo 2* | ***Tyrannochromis macrostoma*** | *Ty. macrostoma* |
| ***Mylochromis anaphyrmus*** | *My. anaphyrmus* | ***Tyrannochromis nigriventer*** | *Ty. nigriventer* |

### Table S4: Genome assembly.

Information of previously published and newly provided assemblies (**bold**) of Lake Malawi cichlid species. The following abbreviations are used in the sequencing technology column: CLR: PacBio Continuous Long Reads; HiFi: PacBio High Fidelity reads; ONT: Oxford Nanopore simplex reads. Detailed information is provided in the corresponding data repositories.

| **Assembly ID** | fAstCal1.2 | **fAstCal1.6** | **fTroMau2.1** | **fAulStu2.1** | **fDipLim1.1** | **fRhaChi1.1** | fOtoArg1.0 | fCopChr1.0 |
|---|---|---|---|---|---|---|---|---|
| **Assembly metadata** | | | | | | | | |
| **Species** | *Astatotilapia calliptera* | *Astatotilapia calliptera* | *Tropheops* sp. 'mauve' | *Aulonocara stuartgranti* | *Diplotaxodon limnothrissa* | *Rhamphochr.* sp. 'Chilingali' | *Otopharynx argyrosoma* | *Copadichromis chrysonotus* |
| **Clade** | *A. calliptera* | *A. calliptera* | *Mbuna* | *Benthic (deep)* | *Diplotaxodon* | *Rhampho-chromis* | *Benthic (shallow)* | *Benthic (utaka)* |
| **Sequencing technology (assembler)** | CLR (Falcon) | CLR (Falcon) | CLR (Falcon) | ONT duplex (Hifiasm) | HiFi (Hifiasm) | CLR (Falcon) | ONT (Shasta) | ONT (Shasta) |
| **Scaffolding technology (scaffolding software)** | Linkage map + Bionano | Hi-C (yahs) | Hi-C (yahs) | Hi-C (yahs) | Hi-C (yahs) | Hi-C (yahs) | - | - |
| **Assembly level** | chromosome | chromosome | chromosome | chromosome | chromosome | chromosome | contig | contig |
| **NCBI accession** | GCA_900246225.3 | <fill> | <fill> | <fill> | PRJEB77457 | <fill> | <fill> | <fill> |
| **Contigs** | | | | | | | | |
| **bases** | 879258901 | 879258901 | 912692380 | 945378724 | 932535246 | 901598841 | 875605137 | 864498723 |
| **sequences** | 739 | 808 | 273 | 54 | 2683 | 640 | 2861 | 6225 |
| **N50** | 4438245 | 4317830 | 8390905 | 37447771 | 599000 | 4280269 | 2499340 | 564332 |
| **L50** | 58 | 59 | 34 | 11 | 453 | 66 | 106 | 462 |
| **N90** | 786390 | 692411 | 1873351 | 13398811 | 176028 | 1046806 | 388416 | 96394 |
| **L90** | 236 | 248 | 125 | 26 | 1544 | 233 | 414 | 1793 |
| **Scaffolds** | | | | | | | | |
| **bases** | 880445564 | 879324101 | 912725380 | 945383024 | 932998246 | 901684441 | - | - |
| **sequences** | 249 | 482 | 108 | 23 | 368 | 212 | - | - |
| **N50** | 38669361 | 36155743 | 38071527 | 40335072 | 39087811 | 39184602 | - | - |
| **L50** | 10 | 11 | 11 | 10 | 11 | 10 | - | - |
| **N90** | 31467755 | 29460845 | 31003463 | 35632062 | 31715461 | 33687932 | - | - |
| **L90** | 20 | 22 | 21 | 20 | 21 | 20 | - | - |

#### Table S5: Sample metadata for species cross (separate file).

Metadata of samples from an interspecific cross between *A. calliptera* and *Au. stuartgranti*. Per sample, the file identifies the generation (G0, F1, F2, F3), sex, sequence ID, repository accessions and sequencing depth.

**Supplementary file "Table_S5.tsv"**

#### Table S6: Approximated centromere locations.

Approximated centromere locations in the fAstCal1.2 genome assembly, inferred as described in the methods. Genomic coordinates are provided in bp, and the level of certainty along with the supporting evidence is indicated per chromosome.

| chrom | start | end | evidence | certainty level |
|---|---|---|---|---|
| chr1 | 5500000 | 7000000 | flanks,gaps,wpca | medium |
| chr2 | 16823543 | 16889383 | blat,flanks,gaps,wga,wpca | high |
| chr3 | 5800000 | 6700000 | flanks,gaps | medium |
| chr4 | 27700000 | 28000000 | flanks,gaps,wga,wpca | high |
| chr5 | 35500000 | 37000000 | blat,gaps,wpca | medium |
| chr6 | 17000000 | 18000000 | flanks | low |
| chr7 | 10766357 | 10766456 | gaps,flanks,wga,wpca | high |
| chr8 | 22000000 | 25827708 | gaps,flanks,wpca | low |
| chr9 | 5600000 | 6200000 | blat,flanks,gaps,wga,wpca | high |
| chr10 | 6700000 | 7700000 | blat,flanks,gaps,wga,wpca | medium |
| chr11 | 1 | 500000 | flanks,wga | medium |
| chr12 | 8000000 | 10000000 | blat,flanks,gaps,wga,wpca | low |
| chr13 | 500000 | 1500000 | blat,gaps,flanks,wga,wpca | high |
| chr14 | 33600000 | 33800000 | blat,flanks,wga,wpca | high |
| chr15 | 27425631 | 27425730 | flanks,gaps,wga | high |
| chr16 | 9000000 | 11000000 | blat,flanks,wga | low |
| chr17 | 3000000 | 4000000 | flanks,gaps,wga,wpca | medium |
| chr18 | 32000000 | 33000000 | blat,gaps,flanks,wga,wpca | medium |
| chr19 | 30700000 | 30863130 | blat,flanks,wga | medium |

#### Table S7: Inversion genotypes for whole genome-sequenced individuals (separate file).

Inversion genotype of the focal inversions for all 1,375 whole genome-sequenced samples used in the main analysis (table S1). Per sample, the file provides the sequence ID (connecting it to table S1 and to the variant callset). For each inversion (chr2, chr9, chr10, chr11, chr13) the inversion genotype is reported (see materials and methods) and encoded as one of: 0 (homozygous non-inverted), 1 (heterozygous), 2 (homozygous inverted).

**Supplementary file "Table_S7.tsv"**

#### Table S8: Inversion breakpoint regions.

Inversion breakpoint regions determined using local windowed PC analyses. 50-80 kbp regions around the left (start) and right (end) breakpoint were approximated using high resolution windowed PC analyses. For example on chr9, start_outer=9800000 and start_inner=9850000 define the estimated 50 kbp region in which the start (left breakpoint) of the chromosome 9 inversion is located in the fAstCal1.2 reference assembly, and start_midpoint=9825000 defined the center of these coordinates. All coordinates are in bp.

| chrom | start_midpoint | end_midpoint | start_outer | start_inner | end_inner | end_outer |
|---|---|---|---|---|---|---|
| chr9 | 12025000 | 28860000 | 12000000 | 12050000 | 28820000 | 28900000 |
| chr11 | 6770000 | 28710000 | 6730000 | 6810000 | 28680000 | 28740000 |
| chr10 | 11325000 | 29285000 | 11300000 | 11350000 | 29250000 | 29320000 |
| chr13 | 9810000 | 29570000 | 9780000 | 9840000 | 29540000 | 29600000 |
| chr2 | 9825000 | 32510000 | 9800000 | 9850000 | 32480000 | 32540000 |

#### Table S9: TE assay primer sets.

Sequences and properties of the primer sets used in the inversion genotyping PCR assay. Melting temperature ('™'), GC content ('GC%'), the paired primer ('PrimerPartner') and amplicon size in the fAstCal1.2 reference genome ('size') are reported.

| PrimerName | Sequence | Tm | GC% | PrimerPartner | Size |
|---|---|---|---|---|---|
| chr9_28197353_F | ACCATGACAGGTTTTCACCCT | 59 | 48 | | |
| chr9_28197353_R | ACTCTAACAAACCACCCGCA | 60 | 48 | chr9_28197353_F | 585 |
| chr9_28197353_ins_R | CAGGGTTTCATGCTACTACTGC | 59 | 50 | chr9_28197353_F | 300 |
| chr11_6948751_F | CCCTTCATAGCCGGTCAGTTT | 60 | 52 | | |
| chr11_6948751_R | GTTTGTGGTCGCATGTGTCTT | 60 | 48 | chr11_6948751_F | 599 |
| chr11_6948751_ins_R | ATGTCCGCTGCTAGTTCAGG | 60 | 55 | chr11_6948751_F | 280 |
| chr10_11347003_F | GCTTTGCATCTCATCTCAGCA | 59 | 48 | | |
| chr10_11347003_R | CCTGCACCTCTCAAGAACCA | 60 | 55 | chr10_11347003_R | 531 |
| chr10_11347003_ins_R | TGGGTAACTTTTAACCACAACTTT | 57 | 33 | chr10_11347003_F | 289 |
| chr13_29501513_F | CTGCCAACAGACAAAACCCA | 59 | 50 | | |
| chr13_29501513_R | TCCGGTCCAGGATTCAGCAT | 61 | 55 | chr13_29501513_F | 443 |
| chr13_29501513_ins_R | TAAAATGCACTGAGTGGTCAGC | 59 | 45 | chr13_29501513_F | 271 |
| chr2_32484955_F | ATCCTACTCAGGCCACATCAGT | 60 | 50 | | |
| chr2_32484955_R | CACCAGTAAGGAAGCCCACG | 61 | 60 | chr2_32484955_F | 525 |
| chr2_32484955_ins_R | AAACCTTGCAGACGAAGGTG | 59 | 50 | chr2_32484955_F | 278 |

#### Table S10: Sample metadata for PCR-typed individuals (separate file).

Metadata of 401 samples PCR-typed for inversion state. Per sample, the file contains genus and species name, catch location and sex assignment (see materials and methods).

**Supplementary file “Table_S10.tsv”**

#### Table S11: Inversion genotypes for PCR-typed individuals (separate file).

Inversion genotype of the focal 5 inversions for all 401 PCR-typed samples. Per sample, the file provides the sample ID connecting it with the samples' metadata (table S10). For each inversion (chr2, chr9, chr10, chr11, chr13) the inversion genotype is reported (see materials and methods) and encoded as one of: 0 (homozygous non-inverted), 1 (heterozygous), 2 (homozygous inverted) or NA (if the PCR run failed twice).

**Supplementary file "Table_S11.tsv"**

#### Table S12: Sample metadata for outgroup dataset (separate file).

Non-Malawi specimens used in the gene flow analysis for the investigation of the origin of the chromosome 9 haplotype. Per sample, the file contains the genus and species name, assignment to a taxonomic group ('group'), sample number, catch location, sex assignment, sequence ID (used as sample ID in the VCF), sample/individual ID, repository accessions (BioSample and BioProject) and sequencing depth.

**Supplementary file "Table_S12.tsv"**

#### Table S13: Excess allele sharing (ABBA-BABA) signals of non-inverted benthic chromosome 9 haplotype with outgroup species (separate file).

Output of Dsuite Dtrios (_BBAA.txt file) subsetted for comparisons for which P1 and P2 are the Malawi-like and the "foreign" haplotype in chromosome 9 heterozygous individuals (or vice versa) and P3 is a non-Malawi species (see materials and methods section *Investigation of chromosome 9 haplotype origin*). Note that what is referred to as chr9het_malawi and chr9het_foreign in the materials and methods is called chr9hets_mand chr9hets_o, respectively, in the table.

**Supplementary file "Table_S13.tsv"**

#### Table S14: Depth records of benthic species (separate file).

Summary of depth data for a set of species of the *benthic* subradiation used in this study, derived from our own records (bottom trawling and artisanal fisheries). For each species, the minimum, maximum, and mean depth in metres across all data entries are included. For the bottom trawling data, an entry represents a trawling session, for which depth was approximated as the average depth between the start and end of the session. Species entries for which no data were available are left blank.

**Supplementary file "Table_S14.tsv"**

#### Table S15: Counts of synonymous and nonsynonymous substitutions in highest bins.

Counts of synonymous and nonsynonymous substitutions in highest widening bins, which include sites with inversion correlation from r_ancestral to -1 (for non-inverted haplotype), from r_inverted to 1 (for inverted haplotype).

| | r_ancestral | N_ancestral | S_ancestral | dNdS_ancestral | r_inverted | N_inverted | S_inverted | dNdS_inverted |
|---|---|---|---|---|---|---|---|---|
| **chr9** | -0.989 | 48 | 45 | 0.463 | 0.879 | 31 | 11 | 1.225 |
| **chr11** | -0.929 | 71 | 47 | 0.656 | 0.969 | 44 | 13 | 1.471 |
| **chr10** | -0.969 | 19 | 1 | 8.260 | 0.989 | 59 | 4 | 6.413 |
| **chr13** | -0.959 | 13 | 3 | 1.884 | 0.989 | 28 | 2 | 6.086 |
| **chr2** | -0.979 | 9 | 6 | 0.652 | 0.989 | 20 | 6 | 1.449 |

#### Table S16: Polymorphism sharing between inversion haplotypes

Proportion of variants for which both inverted and non-inverted haplotypes are polymorphic with at least 1, 2 or 4 copies of the minor allele. The left three columns show proportions for inversion correlated SNPs (ICS), while the right three columns show proportions for inversion region SNPs with minor allele frequency > 10%.

| | ICS | | | SNPs MAF > 0.1 | | |
|---|---|---|---|---|---|---|
| **Minimum minor AC for each orientation** | **1** | **2** | **4** | **1** | **2** | **4** |
| **chr2** | 0.72 | 0.48 | 0.19 | 0.61 | 0.51 | 0.44 |
| **chr10** | 0.37 | 0.16 | 0.05 | 0.55 | 0.46 | 0.41 |
| **chr11** | 0.72 | 0.46 | 0.16 | 0.67 | 0.53 | 0.44 |
| **chr13** | 0.64 | 0.34 | 0.11 | 0.61 | 0.48 | 0.41 |

#### Table S17: Tissue groupings used in single cell expression analysis.

Tissues presented in the Daniocell database were grouped into major physiological categories for further analysis. 'Tissue group' indicates how we summarised them into broader categories.

| Tissue group | Tissues name in Daniocell database |
|---|---|
| neurosensory | neural, eye, otic_lateral_line, taste_olfactory |
| coloration | epidermis, pigment, fin |
| other | axial, blastomeres, periderm, mesenchyme, endoderm, muscle, pronephros, ionocytes_mucus, hematopoietic, mural, glial, pgc |

#### Table S18: Relative expression of genes with nsICS in 19 tissues.

Summary table for gene functional analysis, based on zebrafish single cell expression (separate file). Data from this table is used for fig. S46. Counts of target genes which were expressed in specific groups of tissues are summarised by ancestral or inverted haplotype (‘counts’). The sum of all gene expression counts for genes of interest is provided in ‘count_sums’. Additionally, analogous counts and sums for a set of random genes are provided in columns ‘rand_counts’ and ‘rand_count_sums’ (see also Materials and methods section).

**Supplementary file “Table_S18.tsv”**

#### Table S19: Candidate gene tissue specific expression (separate file).

Counts of tissue-specific single cell expression of 315 nsICS genes, gathered from the Daniocell database (*44*, *154*) (see materials and methods). Each gene repeated according to the number of nsICS mutations in it.

**Supplementary file "Table S19.tsv"**

#### Table S20: Background gene tissue specific expression (separate file).

Counts of tissue-specific single cell expression of 12873 genes, orthologous between *Astatotilapia calliptera* and zebrafish. Data gathered from the Daniocell database (*44*, *154*) (see materials and methods).

**Supplementary file "Table_S20.tsv"**

#### Table S21: Relative expression of genes with nsICS in 3 grouped tissues.

Summary table for gene functional analysis, based on zebrafish single cell expression (separate file). Data from this table is used for fig. S47. Counts of target genes which were expressed in specific groups of tissues are summarised by ancestral or inverted haplotype ('counts'). The sum of all gene expression counts for genes of interest is provided in 'count_sums'. Additionally, analogous counts and sums for a set of random genes are provided in columns 'rand_counts' and 'rand_count_sums' (see also Materials and methods section).

**Supplementary file "Table_S21.tsv"**

#### Table S22: Relative expression of genes with nsICS across haplotypes.

Summary table for gene functional analysis, based on zebrafish single cell expression (separate file). Data from this table is used for fig. S48. Counts of target genes which were expressed in specific groups of tissues are summarised by ancestral or inverted haplotype (‘counts’). The sum of all gene expression counts for genes of interest is provided in ‘count_sums’. Additionally, analogous counts and sums for a set of random genes are provided in columns ‘rand_counts’ and ‘rand_count_sums’ (see also Materials and methods section).

**Supplementary file “Table_S22.tsv”**

#### Table S23: Gene ontology (GO) enrichment results (separate file).

Gene ontology terms (Fisher exact test $p<0.05$) in high inversion correlated SNPs ("ICS"; $|r|>=0.99$, $p<10^{-230}$) per inversion.

**Supplementary file "Table_S23.tsv"**

#### Table S24: Summary of selection and functional evidence for genes in inversion regions (separate file).

This table contains all genes that fall inside the five focal inversion regions. ‘GENE INFO’ contains gene IDs, gene coordinates and number or nonsynonymous mutations with ICS score > 200 by absolute value. ‘ICS>200’ and ‘ICS>300’ sections contain per gene counts of nonsynonymous mutations with high inversion correlation score (either 200 or 300, respectively, positive/negative columns correspond to mutations with derived allele correlated with inverted/ancestral inversion state). ‘ICS TOP WINDOWS (0.5%)’ and ‘ICS TOP WINDOWS (0.1%)’ sections signify if the gene falls into the top 0.5% or 0.1% windows of highest average ICS on the respective chromosome. ‘GO ENRICHMENT’ lists if the gene has been assigned enriched GO terms. ‘DGE’ and ‘ASE’ sections describe differentially expressed genes and genes with allele specific expression. “NOTES” contains additional information including short descriptions of gene functions and literature references.

**Supplementary file “Table_S24.tsv”**

#### Table S25: nsICS density

This file contains numbers of nsICS mutations, normalized by gene length. Contains gene name (gene_name field), number of nsICS mutations per gene (nonsyn_mut_count), gene (gene_length), and number of nsICS mutations per nucleotide (density).

**Supplementary file "Table_S25.tsv"**

#### Table S26: Neuroreceptor gene list.

List of neuroreceptors, involved in social and affiliative behaviour which was used to evaluate overrepresentation of neuroreceptors in the inversion regions.

| Receptor type | Gene name |
|---|---|
| glutamate | gria1, gria2, gria3, gria4 |
| NDMA type | grin1, grin2a, grin2b, grin2c, grin2d, grin3a, grin3b |
| glutamatergic | grik1, grik2, grik3, grik4, grik5 |
| oxytocin | oxt, oxtr, oxtrl |
| arginine vasopressin/vasotocin | avp, avpr1a, avpr2a |
| opioid | oprm1, oprk1, oprd1b |
| dopamine | drd1, drd1b, drd2a, drd2l, drd3, drd4a, drd4b, drd6b |
| serotonin | htr1a, htr1b, htr1ab, htr1f, htr2a, htr2b, htr2cl1, htr3a, htra3a, htr4, htr5ab, htr6, htr7a |

#### Table S27: Bower architecture per species.

List of species assigned to bower types ‘pit’ and ‘castle’. Only species with clear assignment to one of these two basic types were considered, species that build more complex bowers were ignored in this analysis.

| **Bower type** | **Species** |
|---|---|
| **pit** | *Aulonocara rostratum*, *Aulonocara stuartgranti*, *Chilotilapia rhoadesii*, *Copadichromis ilesi*, *Copadichromis kawanga*, *Copadichromis mloto*, *Copadichromis virginalis*, *Dimidiochromis compressiceps*, *Dimidiochromis kiwinge*, *Dimidiochromis strigatus*, *Fossorochromis rostratus*, *Hemitaeniochromis urotaenia*, *Lethrinops macrochir*, *Mylochromis melanonotus*, *Mylochromis mollis*, *Mylochromis spilostichus*, *Mylochromis subocularis*, *Naevochromis chrysogaster*, *Nimbochromis linni*, *Nimbochromis livingstonii*, *Nimbochromis polystigma*, *Otopharynx auromarginatus*,*Otopharynx brooksi*, *Otopharynx tetrastigma*, *Protomelas annectens*, *Protomelas fenestratus*, *Protomelas insignis*, *Protomelas marginatus*, *Protomelas ornatus*, *Protomelas similis*, *Protomelas spilopterus*, *Stigmatochromis woodi*, *Taeniochromis holotaenia*, *Taeniolethrinops laticeps*, *Tramitichromis intermedius*, *Tyrannochromis nigriventer* |
| **castle** | *Buccochromis* nototaenia, *Buccochromis rhoadesii*, *Champsochromis caeruleus*, *Copadichromis jacksoni*, *Copadichromis likomae*, *Ctenopharynx pictus*, *Hemitilapia oxyrhynchus*, *Lethrinops furcifer*, *Lethrinops parvidens*, *Mchenga* sp. ‘black-yellow’, *Mchenga* sp. ‘Chiofu’, *Mchenga* sp. ‘Chirombo-1’, *Mchenga* sp. ‘Chirombo-2’, *Mylochromis anaphyrmus*, *Otopharynx argyrosoma*, *Otopharynx decorus*, *Protomelas kirkii*, *Taeniolethrinops furcicauda*, *Tramitichromis brevis*, *Tramitichromis* sp. ‘east-coast-shallow’ |

#### Table S28: Association between inversions and sex.

Input data used for Fisher’s exact tests for association between the five focal inversions and sex of individuals from species with at least one heterozygous sample (see materials and methods). Per inversion, the following sample counts are reported: Total number of females/males, number of females/males without the inversion, number of females/males heterozygous for the inversion, number of females/males homozygous for the inversion. Furthermore, the output of the test statistic (p-value and odd ratio) is reported per inversion.

| **inversion** | **chr9** | | **chr11** | | **chr10** | | **chr13** | | **chr2** | |
|---|---|---|---|---|---|---|---|---|---|---|
| **sex** | **F** | **M** | **F** | **M** | **F** | **M** | **F** | **M** | **F** | **M** |
| **total** | 162 | 167 | 131 | 152 | 8 | 59 | 128 | 139 | 59 | 74 |
| **non-inverted** | 0 | 0 | 9 | 39 | 1 | 3 | 90 | 86 | 37 | 44 |
| **heterozygous** | 14 | 66 | 46 | 90 | 3 | 49 | 15 | 28 | 11 | 15 |
| **inverted** | 148 | 101 | 76 | 23 | 4 | 7 | 23 | 25 | 11 | 15 |
| **p-value** | $3.7\times10^{-11}$ | | $5.3\times10^{-11}$ | | 0.0143 | | 0.287 | | 1.0 | |
| **odd ratio** | 0.145 | | 0.155 | | 0.107 | | 0.582 | | 1.0 | |

#### Table S29: tblastn analysis of INDELS in *Aulonocara stuartgranti*.

Table of results from tblastn analysis of INDELS against known sex determining gene protein sequences.

| Type | fAulStu2.1 sequence location | gene | Identity (%) | E-val | gene function |
|---|---|---|---|---|---|
| Deletion | chr6:39163238 | tgfbr1b | 74.194 | 2.07E-08 | control of cell growth, cell proliferation, cell differentiation, and apoptosis |
| Deletion | chr17:19274331 | tgfbr1b | 90.909 | 2.45E-06 | control of cell growth, cell proliferation, cell differentiation, and apoptosis |
| Insertion | chr14:37381200 | tdrd1 | 48.837 | 2.65E-13 | suppression of transposable elements during spermatogenesis |
| Deletion | chr6:5669829 | cbx2 | 41.86 | 1.31E-07 | stimulate the male pathway and concurrently inhibit the female pathway |

#### Table S30: Tissue expression of artificial ICS (19 tissues).

This table summarizes the number of chromosomes which had significant overrepresentation of genes expressed in one of 19 tissues in the analysis of artificial nsICS obtained from in-silico expansion of a random haplotype (see text S3). Fisher test, Benjamini-Hochberg false discovery rate correction.

| Tissues | Chromosomes tested | Significant |
|---|---|---|
| taste | 176 | 1 |
| otic | 176 | 0 |
| eye | 176 | 1 |
| neural | 176 | 4 |
| fin | 176 | 2 |
| pigment | 176 | 2 |
| epidermis | 176 | 0 |
| pgc | 176 | 1 |
| glia | 176 | 6 |
| mural | 176 | 0 |
| hematopoietic | 176 | 3 |
| ionocytes_mucus | 176 | 1 |
| pronephros | 176 | 2 |
| muscle | 176 | 2 |
| endoderm | 176 | 2 |
| mesenchyme | 176 | 0 |
| periderm | 176 | 2 |
| blastomeres | 176 | 0 |
| axial | 176 | 2 |

#### Table S31: Tissue expression of artificial ICS (3 tissues).

This table summarizes the number of chromosomes which had significant overrepresentation of genes expressed in one of 3 tissue groups in the analysis of artificial nsICS obtained from in-silico expansion of a random haplotype (see text S3). Fisher test, Benjamini-Hochberg false discovery rate correction.

| Tissues | Chromosomes tested | Significant |
|---|---|---|
| coloration | 176 | 1 |
| neurosensory | 176 | 2 |
| other | 176 | 1 |

### Data S1 to S8

#### Data S1: Phylogenetic tree containing all 684 benthic individuals.

Phylogenetic tree reconstruction of benthic samples based on trees reconstructed in 100 kbp windows with IQtree2 and summarised with ASTRAL III (see materials and methods). Branch lengths are given in coalescent units, branch supports measured as local posterior probabilities.

**Data_S1.nwk**

#### Data S2: Phylogenetic tree containing 93 taxa of all studied clades.

Phylogenetic tree reconstruction of a subset of samples from all clades based on trees reconstructed in 100 kbp windows with IQtree2 and summarised with ASTRAL III (see materials and methods). Branch lengths are given in coalescent units, branch supports measured as local posterior probabilities.

**Data_S2.nwk**

#### Data S3: Inversion states per species per sex

Inversion states per species per sex

**Data_S3.xlsx**

#### Data S4: Phylogenetic relationships within the chromosome 9 inversion.

High resolution version of fig. S29.

**Data_S4.pdf**

#### Data S5: Phylogenetic relationships within the chromosome 11 inversion.

High resolution version of fig. S30.

**Data_S5.pdf**

#### Data S6: Phylogenetic relationships within the chromosome 10 inversion.

High resolution version of fig. S31.

**Data_S6.pdf**

#### Data S7: Phylogenetic relationships within the chromosome 13 inversion.

High resolution version of fig. S32.

**Data_S7.pdf**

#### Data S8: Phylogenetic relationships within the chromosome 2 inversion.

High resolution version of fig. S33.

**Data_S8.pdf**

100. A. Rhie, S. A. McCarthy, O. Fedrigo, J. Damas, G. Formenti, S. Koren, M. Uliano-Silva, W. Chow, A. Fungtammasan, J. Kim, C. Lee, B. J. Ko, M. Chaisson, G. L. Gedman, L. J. Cantin, F. Thibaud-Nissen, L. Haggerty, I. Bista, M. Smith, B. Haase, J. Mountcastle, S. Winkler, S. Paez, J. Howard, S. C. Vernes, T. M. Lama, F. Grutzner, W. C. Warren, C. N. Balakrishnan, D. Burt, J. M. George, M. T. Biegler, D. Iorns, A. Digby, D. Eason, B. Robertson, T. Edwards, M. Wilkinson, G. Turner, A. Meyer, A. F. Kautt, P. Franchini, H. W. Detrich 3rd, H. Svardal, M. Wagner, G. J. P. Naylor, M. Pippel, M. Malinsky, M. Mooney, M. Simbirsky, B. T. Hannigan, T. Pesout, M. Houck,

A. Misuraca, S. B. Kingan, R. Hall, Z. Kronenberg, I. Sović, C. Dunn, Z. Ning, A. Hastie, J. Lee, S. Selvaraj, R. E. Green, N. H. Putnam, I. Gut, J. Ghurye, E. Garrison, Y. Sims, J. Collins, S. Pelan, J. Torrance, A. Tracey, J. Wood, R. E. Dagnew, D. Guan, S. E. London, D. F. Clayton, C. V. Mello, S. R. Friedrich, P. V. Lovell, E. Osipova, F. O. Al-Ajli, S. Secomandi, H. Kim, C. Theofanopoulou, M. Hiller, Y. Zhou, R. S. Harris, K. D. Makova, P. Medvedev, J. Hoffman, P. Masterson, K. Clark, F. Martin, K. Howe, P. Flicek, B. P. Walenz, W. Kwak, H. Clawson, M. Diekhans, L. Nassar, B. Paten, R. H. S. Kraus, A. J. Crawford, M. T. P. Gilbert, G. Zhang, B. Venkatesh, R. W. Murphy, K.-P. Koepfli, B. Shapiro, W. E. Johnson, F. Di Palma, T. Marques-Bonet, E. C. Teeling, T. Warnow, J. M. Graves, O. A. Ryder, D. Haussler, S. J. O'Brien, J. Korlach, H. A. Lewin, K. Howe, E. W. Myers, R. Durbin, A. M. Phillippy, E. D. Jarvis, Towards complete and error-free genome assemblies of all vertebrate species. *Nature* **592**, 737–746 (2021).

117. A. Smit, R. Hubley, P. Green, RepeatMasker software program (computer program), ver. 3.1.8. Seattle: Institute for Systems Biology. (2007).

133. X. Yi, Y. Liang, E. Huerta-Sanchez, X. Jin, Z. X. P. Cuo, J. E. Pool, X. Xu, H. Jiang, N. Vinckenbosch, T. S. Korneliussen, H. Zheng, T. Liu, W. He, K. Li, R. Luo, X. Nie, H. Wu, M. Zhao, H. Cao, J. Zou, Y. Shan, S. Li, Q. Yang, Asan, P. Ni, G. Tian, J. Xu, X. Liu, T. Jiang, R. Wu, G. Zhou, M. Tang, J. Qin, T. Wang, S. Feng, G. Li, Huasang, J. Luosang, W. Wang, F. Chen, Y. Wang, X. Zheng, Z. Li, Z. Bianba, G. Yang, X. Wang, S. Tang, G. Gao, Y. Chen, Z. Luo, L. Gusang, Z. Cao, Q. Zhang, W. Ouyang, X. Ren, H. Liang, H. Zheng, Y. Huang, J. Li, L. Bolund, K. Kristiansen, Y. Li, Y. Zhang, X. Zhang, R. Li, S. Li, H. Yang, R. Nielsen, J. Wang, J. Wang, Sequencing of 50 human exomes reveals adaptation to high altitude. *Science* **329**, 75–78 (2010).

153. O. Seehausen, Lake Victoria rock cichlids : taxonomy, ecology, and distribution. *(No Title)* (1996).

159. A. L. Tan, S. Mohanty, J. Guo, A. C. Lekven, B. B. Riley, Pax2a, Sp5a and Sp5l act downstream

of Fgf and Wnt to coordinate sensory-neural patterning in the inner ear. *Dev Biol* **492**, 139–153 (2022).
