## Supplementary figures and images for "Introgression dynamics of sex-linked chromosomal inversions shape the Malawi cichlid adaptive radiation"

### Data_S4.pdf

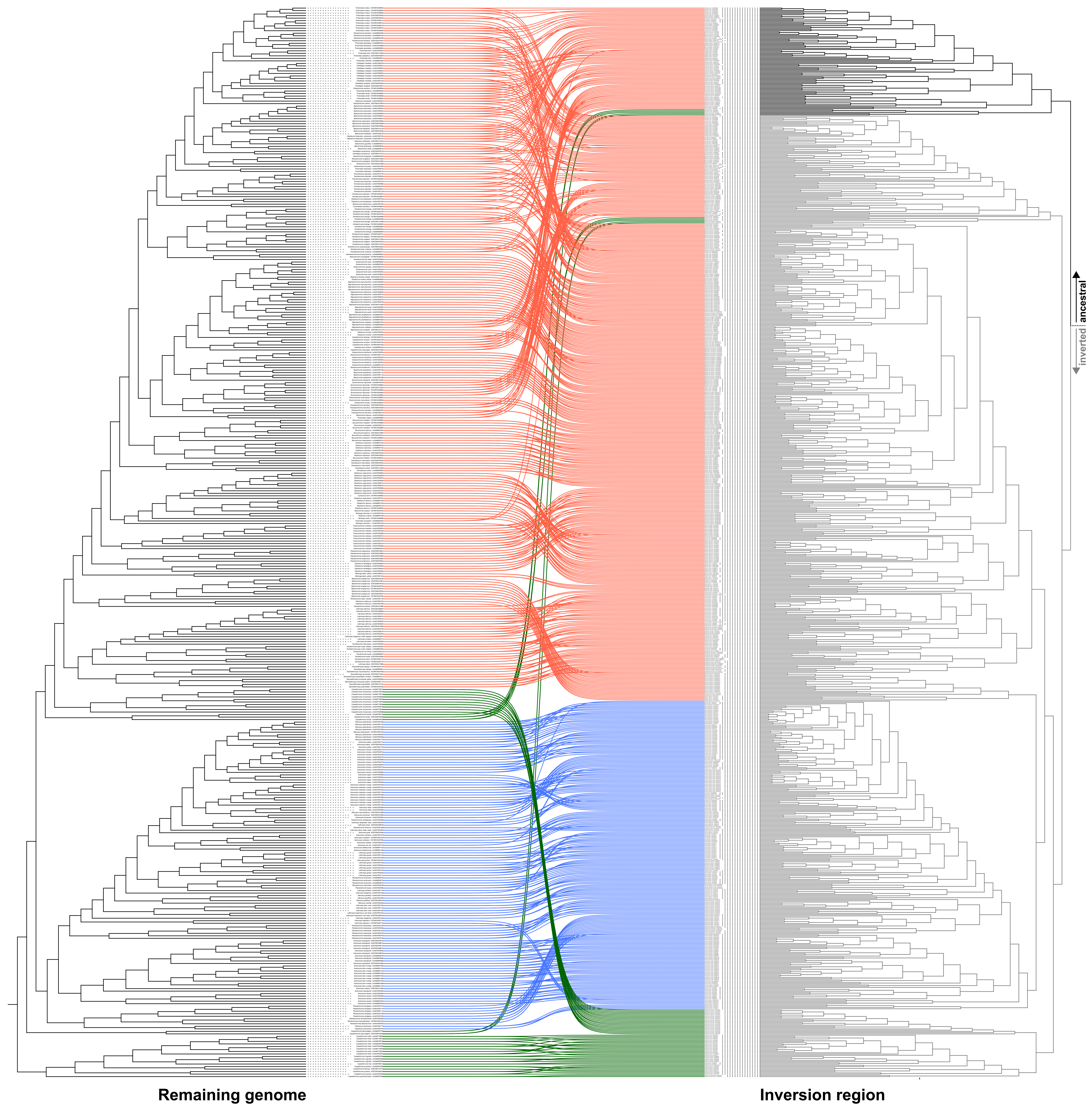

### Data_S5.pdf

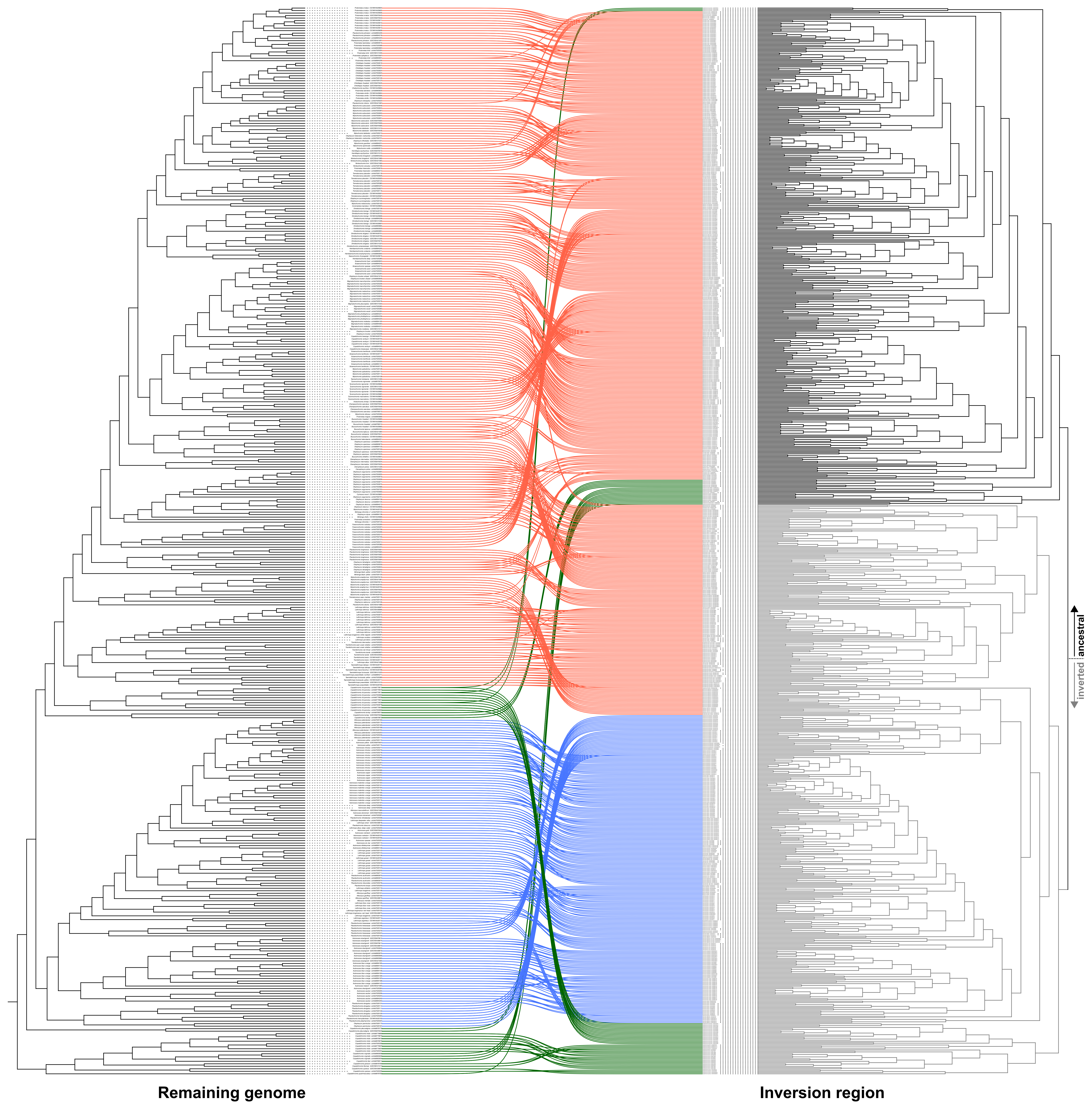

### Data_S6.pdf

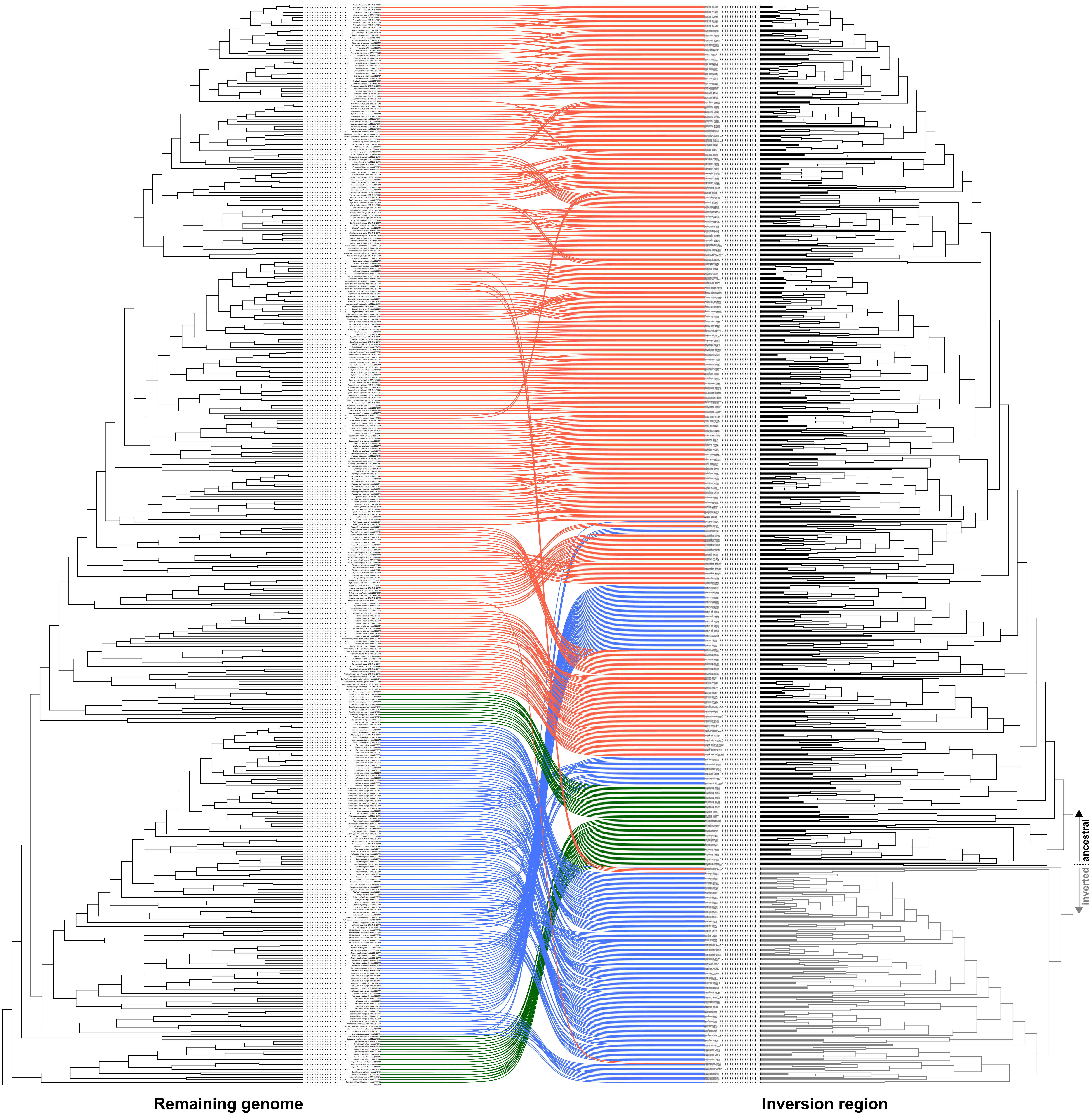

### Data_S7.pdf

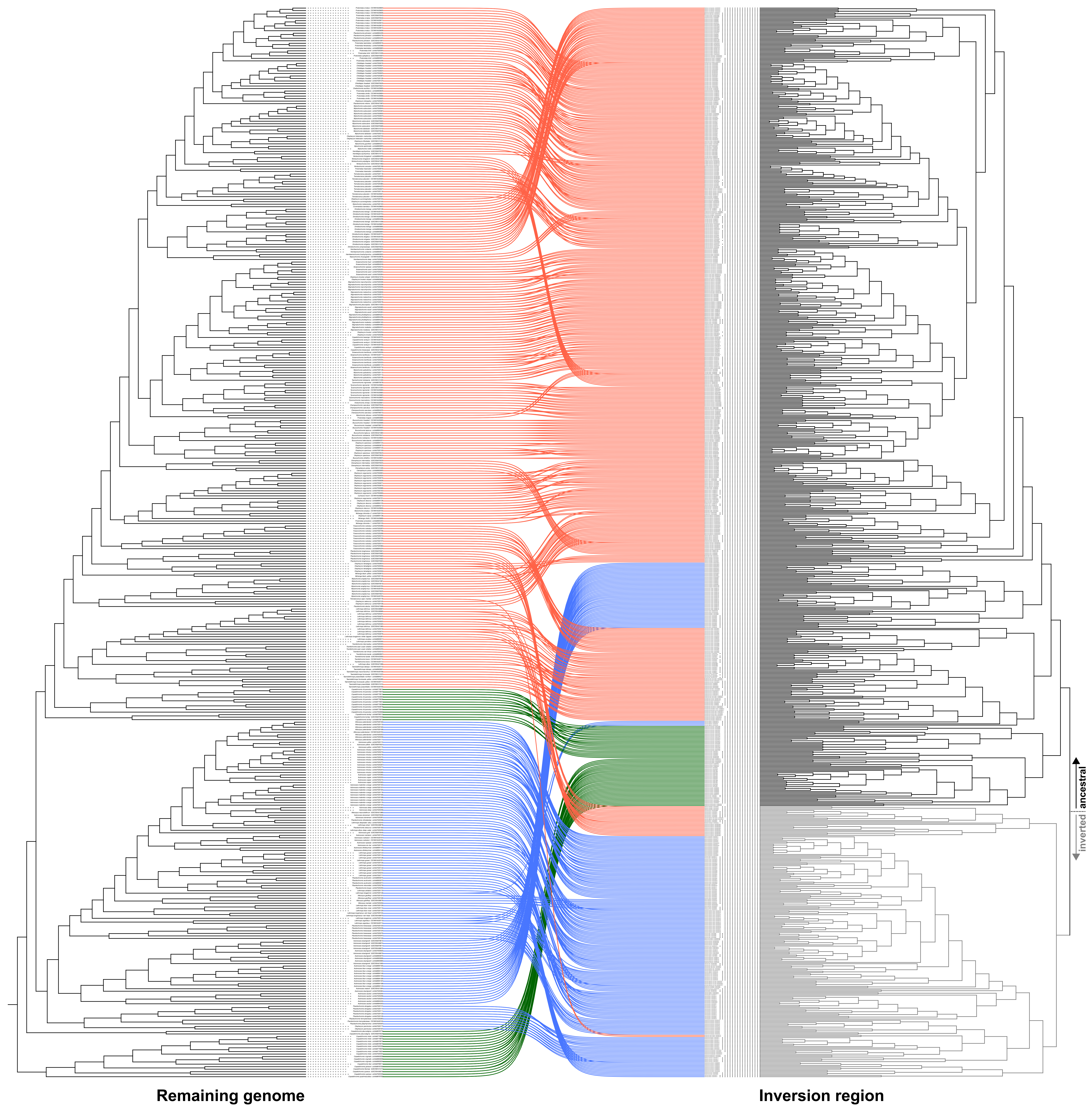

### Data_S8.pdf

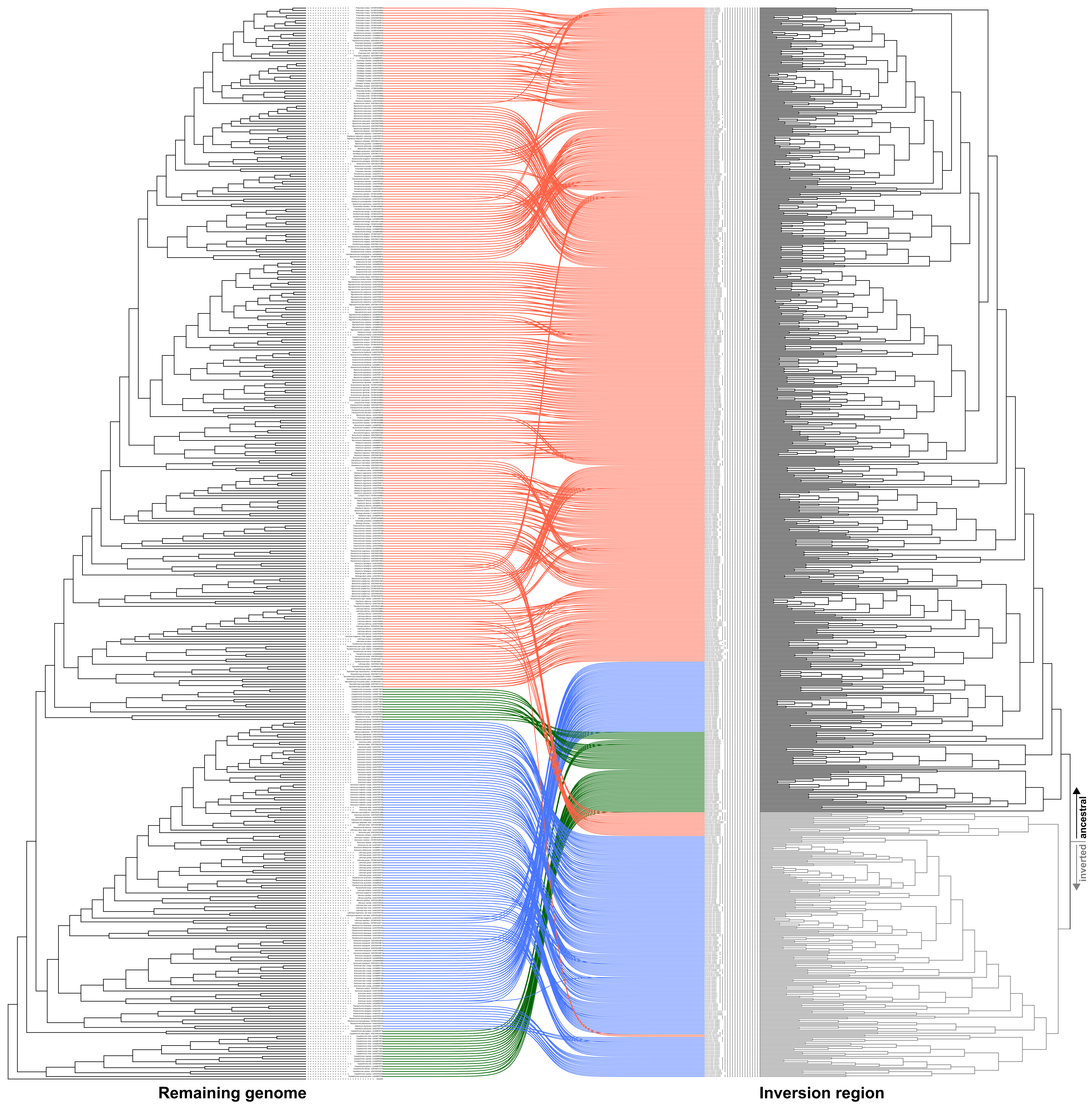
